# Altered immune and treatment response gene expression signatures among poverty-exposed children with B-ALL

**DOI:** 10.64898/2026.01.27.702019

**Authors:** Amy Guillaumet-Adkins, Noori Sotudeh, Sayalee Potdar, Tushara Vijaykumar, Monica Nair, Valeriya Dimitrova, Julia Frede, Yana Pikman, Marian Harris, Andrew E. Place, Lewis B. Silverman, Jens G. Lohr, Kira Bona, Birgit Knoechel

## Abstract

Children diagnosed with cancer typically receive standardized treatment regimens. Despite highly protocolized care, children living in poverty experience a greater risk of cancer relapse and higher mortality compared to their more affluent peers.^1,2^ Acute lymphoblastic leukemia (ALL) is the most prevalent childhood cancer, and children with ALL exposed to poverty are more likely to experience early relapse.^3^ Using single-cell RNA sequencing to analyze leukemic blasts and their microenvironment at diagnosis we found that poverty-exposed patients with standard-risk B-ALL exhibit transcriptional signatures of steroid resistance at time of diagnosis. Additionally, we observe increased expression of inflammatory signatures in myeloid cells and reduced effector signatures in CD8+ T-cells in children with B-ALL living in poverty. Further investigation of the mechanisms underlying these associations may identify opportunities for risk-adapted therapeutic strategies to improve disease outcomes in pediatric ALL.

## INTRODUCTION

Acute lymphoblastic leukemia (ALL) is the most common cancer of childhood, and 20% of children will suffer relapsed disease.^4–6^ Despite improvements in therapy, only 50% of children who experience relapse will be long-term survivors.^6^ Elucidating mechanisms of treatment resistance among populations at risk of relapse is essential to identify opportunities to improve pediatric ALL outcomes.

Early ALL relapse has been defined as relapse <36 months in complete remission^6^ and is associated with particularly poor survival. Recent data have identified children exposed to poverty as a cohort at significantly higher risk of early relapse compared to unexposed children, despite uniform treatment on multi-center clinical trials. Among children treated on Dana-Farber Cancer Institute (DFCI) ALL Consortium trials who experienced leukemia relapse, 92% of poverty-exposed children experienced early relapse compared to 48% of unexposed children. ^3,7^

Poverty is associated with profound health disparities in the US in diseases ranging from asthma to cancer.^8–10^ While structural and sociobehavioral mechanisms underlying these associations—including access to care and adherence—have been delineated, they only partly explain observed disparities.^11^ Emerging evidence suggests that gene-environment interactions may contribute to health outcome disparities in asthma, obesity and cardiovascular disease—and help to explain the influence of stress on disease outcomes.^12,13^ Early childhood exposure to poverty is associated with “toxic stress,” defined by excessive activation of stress regulatory systems that impacts neuroendocrine stress response axes including upregulation of the hypothalamic-pituitary-axis, elevated cortisol and cytokine responses to stress, and altered metabolism,^14–16^ yet no studies have been performed to assess whether similar responses might be at play among children with cancer. Understanding the mechanistic pathways underlying these associations has the potential to inform both therapeutic targets and risk-adapted therapy.

Single-cell analyses have emerged as a powerful way to gain information about heterogeneity within cancer cell populations and have illustrated the importance of co-existing diverse genetic, epigenetic or transcriptional subpopulations for treatment response.^17,18^ We hypothesized that single-cell transcriptomics would allow to capture leukemia-intrinsic and immune microenvironmental differences in pathways associated with drug resistance between leukemia and normal immune cells from poverty-exposed and unexposed children with ALL. To address these hypotheses, we employed full-length single-cell RNA-sequencing of leukemic blasts and microenvironmental cells from poverty-exposed and unexposed children with B-cell ALL (B-ALL). Notably, we identified several differentially enriched pathways in leukemia cells from poverty-exposed and unexposed patients including steroid responsiveness and signatures consistent with altered immune responses in the leukemia microenvironment that may be associated with treatment resistance.

## RESULTS

### Single-cell transcriptomic profiling delineates malignant from normal cells in low-risk pediatric B-ALL

We assembled a cohort of poverty-exposed (N=7) (defined as both parent-reported low-income [annual household income <200% FPL] and at least one unmet basic resource need [HMH; food, utility or housing insecurity) and unexposed (N=7) children with *de novo* B-cell ALL who had consented to biobanking of diagnostic bone marrow specimens to assess transcriptional profiles of lymphoblasts and microenvironmental cell populations at diagnosis. To minimize heterogeneity, we excluded patients with high-risk immunophenotype, *KMT2A* or *BCR-ABL1* gene rearrangements, or high-risk cytogenetic abnormalities (low hypodiploidy; t(17;19); iAMP21). The remaining biospecimens included B-cell immunophenotype, *ETV6-RUNX1* translocations or hyperdiploid karyotypes—two genetic features frequently associated with standard risk prognosis. Clinical, immunophenotype and molecular characteristics are listed in Table 1.

**Table 1.**
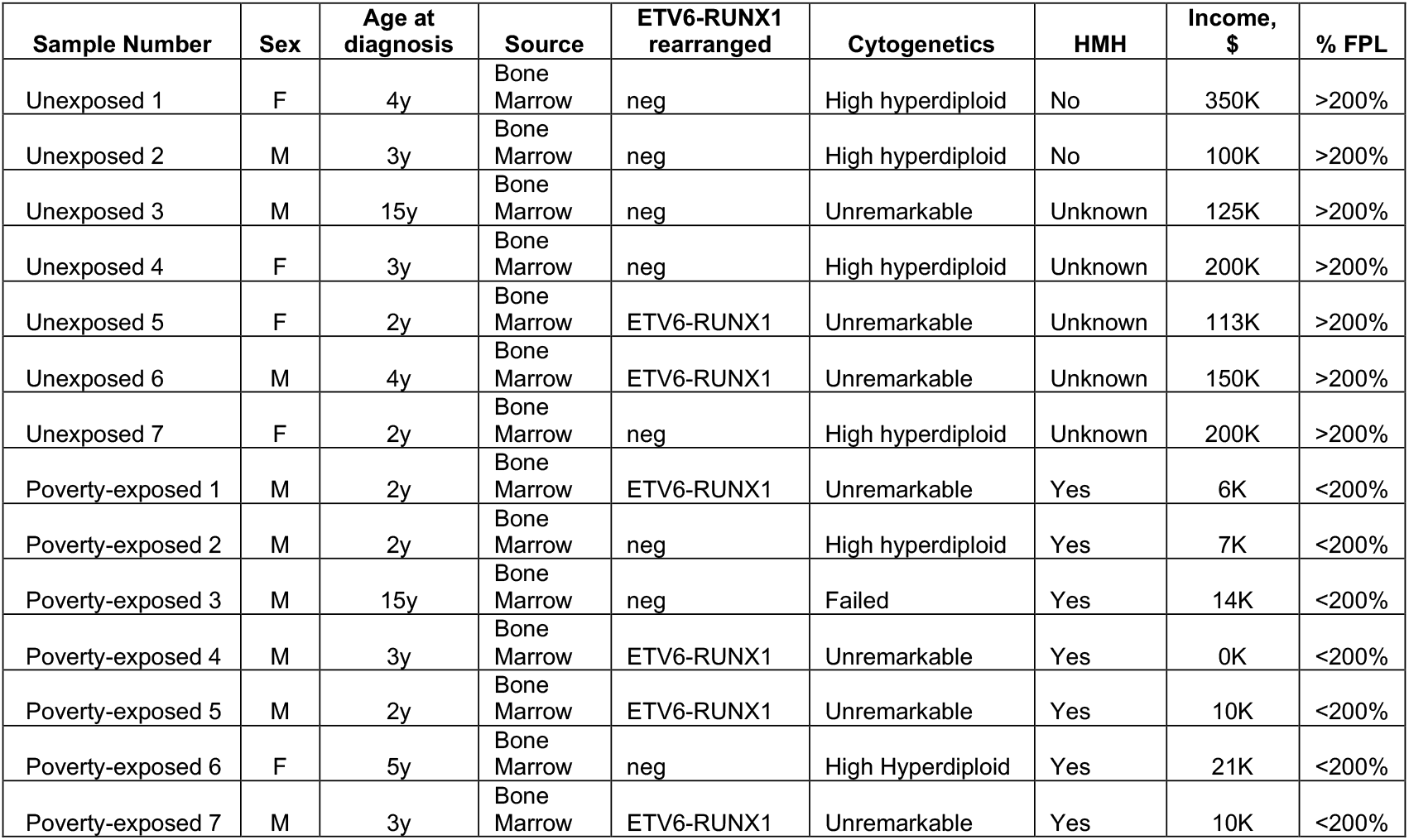
Sociodemographic, clinical, and molecular characteristics at time of diagnosis for N=14 children with standard-risk de novo B-cell ALL. Poverty was defined *a priori* as <u>both</u> parent-reported low-income (annual household income <200% Federal Poverty Level [FPL]) and the presence of household material hardship ([HMH]; food, housing, or utility insecurity).

| Sample Number | Sex | Age at diagnosis | Source | ETV6-RUNX1 rearranged | Cytogenetics | HMH | Income, \$ | % FPL |
| --- | --- | --- | --- | --- | --- | --- | --- | --- |
| Unexposed 1 | F | 4y | Bone Marrow | neg | High hyperdiploid | No | 350K | >200% |
| Unexposed 2 | M | 3y | Bone Marrow | neg | High hyperdiploid | No | 100K | >200% |
| Unexposed 3 | M | 15y | Bone Marrow | neg | Unremarkable | Unknown | 125K | >200% |
| Unexposed 4 | F | 3y | Bone Marrow | neg | High hyperdiploid | Unknown | 200K | >200% |
| Unexposed 5 | F | 2y | Bone Marrow | ETV6-RUNX1 | Unremarkable | Unknown | 113K | >200% |
| Unexposed 6 | M | 4y | Bone Marrow | ETV6-RUNX1 | Unremarkable | Unknown | 150K | >200% |
| Unexposed 7 | F | 2y | Bone Marrow | neg | High hyperdiploid | Unknown | 200K | >200% |
| Poverty-exposed 1 | M | 2y | Bone Marrow | ETV6-RUNX1 | Unremarkable | Yes | 6K | <200% |
| Poverty-exposed 2 | M | 2y | Bone Marrow | neg | High hyperdiploid | Yes | 7K | <200% |
| Poverty-exposed 3 | M | 15y | Bone Marrow | neg | Failed | Yes | 14K | <200% |
| Poverty-exposed 4 | M | 3y | Bone Marrow | ETV6-RUNX1 | Unremarkable | Yes | 0K | <200% |
| Poverty-exposed 5 | M | 2y | Bone Marrow | ETV6-RUNX1 | Unremarkable | Yes | 10K | <200% |
| Poverty-exposed 6 | F | 5y | Bone Marrow | neg | High Hyperdiploid | Yes | 21K | <200% |
| Poverty-exposed 7 | M | 3y | Bone Marrow | ETV6-RUNX1 | Unremarkable | Yes | 10K | <200% |

We enriched for leukemia cells from bone marrow by sorting on CD45low expressing blasts and also collected normal T-cells, B-cells, myeloid and NK cells by sorting on appropriate surface markers (CD45high CD3+ T-cells, CD45high. CD19+ B-cells, CD45high CD14+ myeloid cells; Figure 1A; see methods). Percentage of each sorted cell type is listed in Table 2.

**Figure 1.**
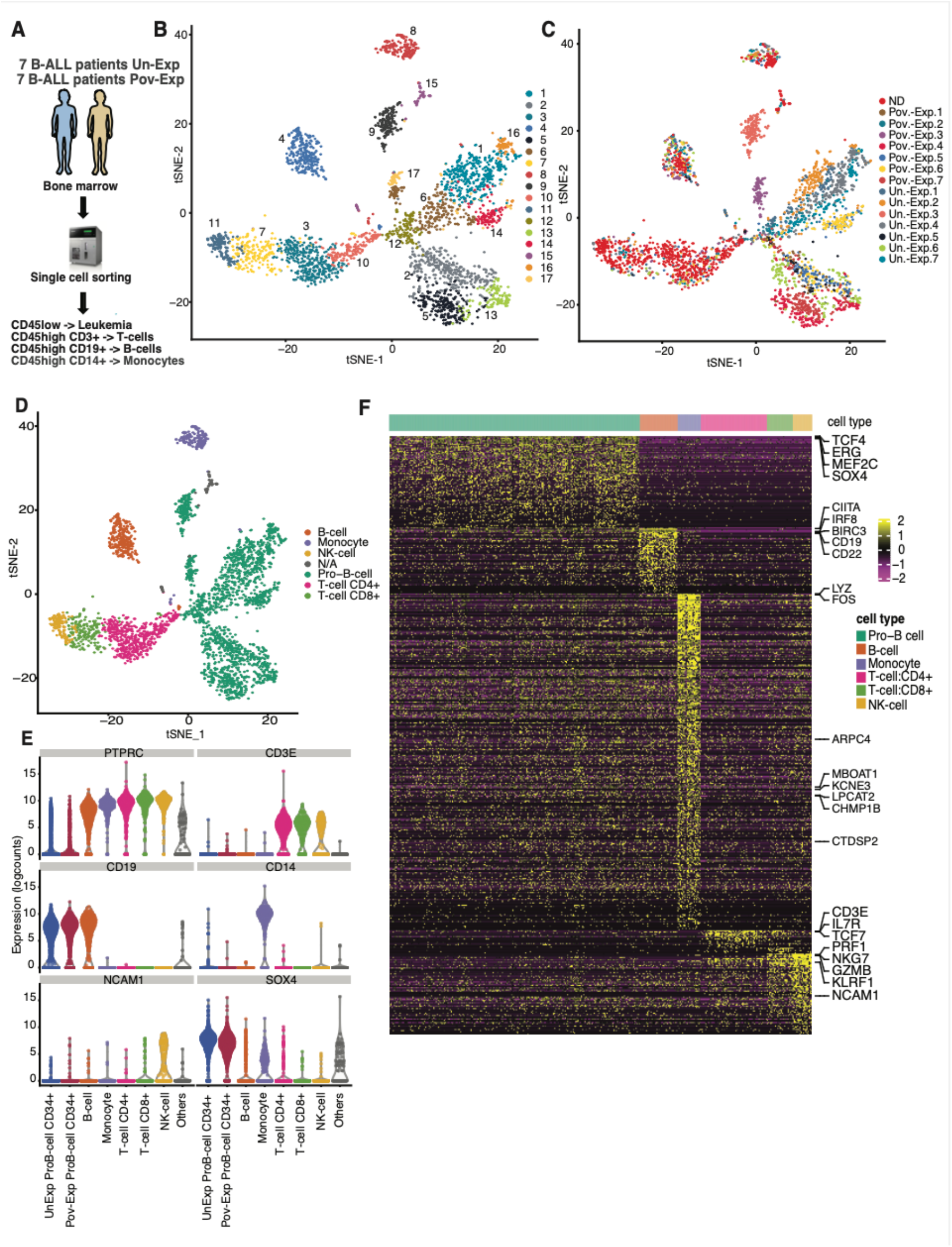
Single-cell transcriptome analysis of pediatric B-ALL in poverty-exposed versus unexposed children. (A) Schematic representation depicting the cohort, sample collection, processing, and sorting of cells for single-cell transcriptional profiling using SMART-seq2 protocol. (B) t-stochastic network embedding (t-SNE) of single-cell gene expression reveals patient-specific and non-specific clustering (ND = normal donor; pov.-exp. = poverty-exposed, un-exp. = unexposed). (C) PAGODA2 analysis demonstrating 17 clusters. (D) t-SNE annotated by celltype as assigned by singleR. (NA = Not Available) (E) Violin plots of canonical marker genes for white blood cells (CD45+), T-cells, B-cells, monocytes, NK-cells, and B-ALL blasts. (F) Expression of marker genes for each cell type is depicted as a heatmap.

After single-cell sorting, we performed full-length single-cell transcriptome analysis of 3,166 malignant (852 “poverty-exposed” and 936 “unexposed”) and 1,378 normal immune cells (T-cells, B-cells and myeloid cells) using the Smartseq2 protocol for full-length RNAseq.^19^ For comparison, we also included normal donor cells from 6 normal donors from published datasets in our analyses (GSE16191, GSE162337).^18,20^ As these normal donors were all from adult individuals, we verified that main cell types and proportions were similar for pediatric individuals by comparing with public pediatric datasets downloaded from GSE132509 (Figure S2).^21^

After several single-cell RNA-seq quality control filtering steps (see Methods and Figure S1), we performed t-Stochastic Neighbor Embedding (t-SNE)^22^ clustering visualization to explore cellular compositions, which revealed 17 distinct cell clusters (Figure 1B). Seven clusters (clusters 4, 7, 8, 10, 11, 15 and 16) were shared between patients and normal donors, thus presumably representing normal immune cells, whereas the remaining clusters (1, 2, 5, 9, 13, 14, and 17) were specific to B-ALL patients (Figure 1C). Lineage annotation of the single-cell profiles with singleR^23^ based on BLUEPRINT^24^ and Human Cell Atlas datasets^25 26^ of immune cell populations revealed that shared clusters indeed represented CD4+ T-cells, CD8+ T-cells, NK-cells, B-cells, and myeloid cells, respectively; whereas leukemic cells fell into clusters annotated as Pro-B cells, consistent with their developmental state (Figure 1D, Figure S3). Additionally, differential expression analysis of well-established canonical lineage marker genes distinguished normal from leukemic clusters (Figure 1E, F). For example, leukemic clusters 1, 2, 5, 9, 13, 14 and 17 expressed *TCF4, ERG, MEF2C* and *SOX4*, all of which are transcription factors with known function in B-ALL.^27–29^ Cluster 4 contained B-cells as demonstrated by expression of *IRF8, CIITA, BIRC3, CD19* and *CD22*. T-cells localized into clusters 3, 7, and 10 with strong expression of *IL7R, CD3G, CD3D*, and *TCF7*. Cluster 8 contained myeloid cells that express *LYZ* and *FOS*. NK cells with expression of the NK markers *NKG7, PRF1* and *GZMB* were found in cluster 11 (Figure 1 C, D).

To ensure that clusters 1, 2, 5, 9, 13, 14, and 17 were indeed malignant, we called arm level copy number variations (CNV) in normal and presumed malignant clusters using InferCNV (see methods) CNVs were limited to leukemic clusters, with copy number events being most prominent in hyperdiploid B-ALL samples (Figure S4).

### Leukemia cells fall into distinct clusters based on underlying genetic characteristics and poverty exposure

Interestingly, while some leukemic clusters were patient-specific, other clusters contained cells of several patients (Figure 1B, C). To understand leukemic cluster formation, we first tested whether underlying genetic events might be causal for cluster formation. Leukemic cells that were hyperdiploid or carried ETV6-RUNX1 translocations clustered separately in t-SNE (Table 1; Figure 2A). Aberrant developmental hierarchies have been described in many cancers including in acute leukemia and have been associated with treatment failure.^17^ To address whether leukemic cells from poverty-exposed or unexposed patients may demonstrate aberrant differentiation states, we performed Louvain clustering^30^ focusing on genes involved in hematopoietic progenitor and B-lymphoid differentiation using validated signatures from the Human Cell Atlas (Table S1, see methods). This analysis revealed 7 clusters (Figure 2B). Leukemia cells from poverty-exposed patients were enriched in clusters 3, 4, 5, 7 while leukemic cells from unexposed patients dominated in clusters 1, 2 and 6 (Figure 2C, D). Of note, all 7 clusters contained cells from both genetic subtypes, even though some were enriched in cells from either hyperdiploid or ETV6-RUNX1 translocated leukemias (i.e., clusters 2 and 4 enriched in hyperdiploid, clusters 3 and 5 enriched in ETV6-RUNX1 translocated specimens (Figure S5), suggesting that genetic events alone were not solely responsible for cluster formation.

**Figure 2.**
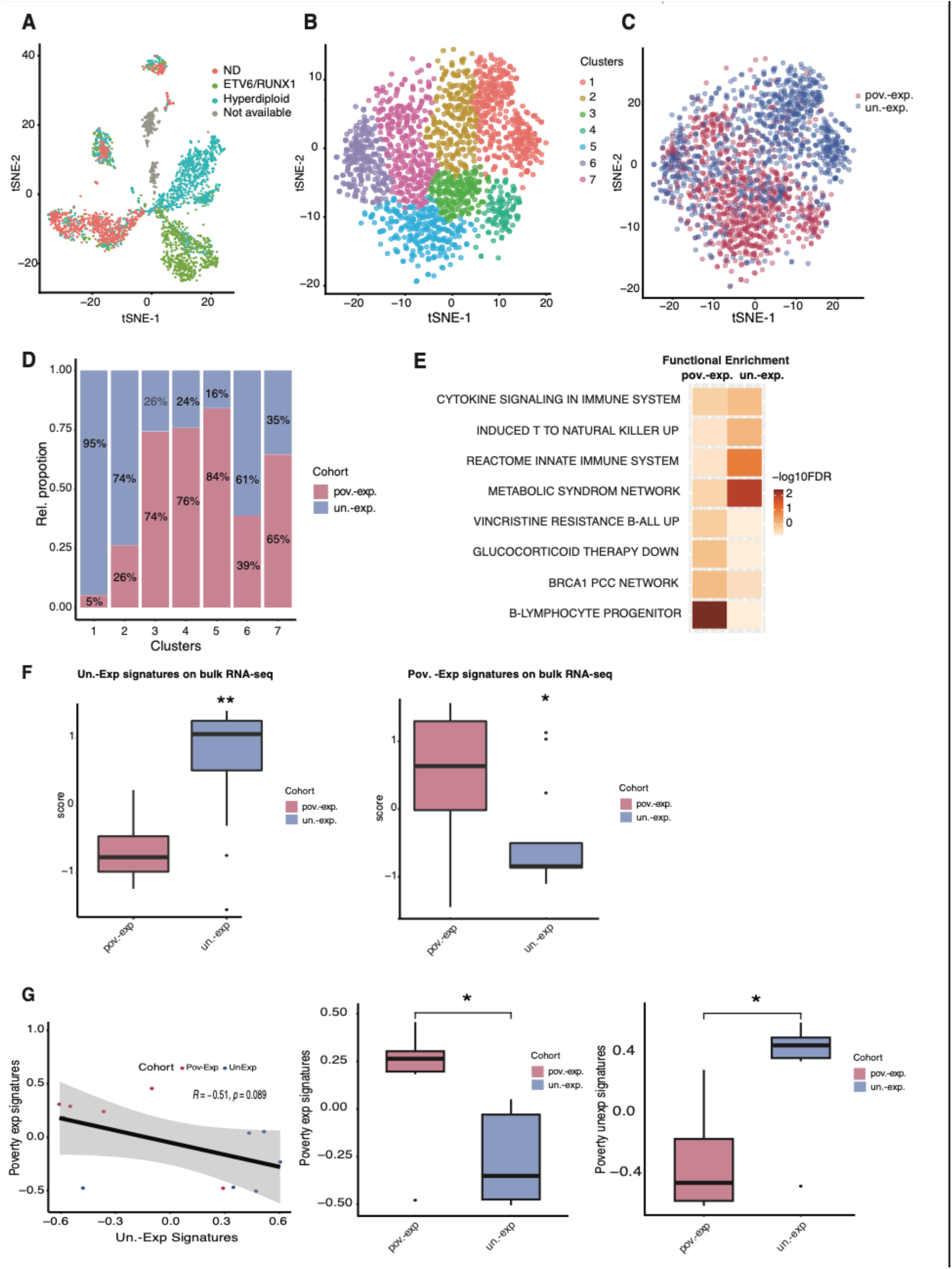
Leukemia blasts from poverty-exposed pediatric B-ALL patients carry distinct transcriptome features. (A) t-SNE annotated by known cytogenetic abnormalities (ND – normal donor). (B) t-SNE plot of malignant cells clustered based on expression of known progenitor and lymphoid differentiation signatures (see methods). (C) t-SNE annotated by poverty-exposure (pov.-exp. = poverty-exposed, un-exp. = unexposed). (D) Relative percentage of single leukemic cells from poverty-exposed and unexposed patients depicted stacked bar plot. (E) GSEA analysis comparing clusters enriched in single cells from poverty-exposed or unexposed patients (poverty-exposed = clusters 3, 4, 5 and 7, unexposed = clusters 1, 2, and 6). (F) Expression of poverty-exposed and unexposed signatures (see methods) in bulk RNA-seq (n = 14; \**p*<0.05, *\*\*p*<0.01). using Wilcoxon rank test). (G) Representative correlation of poverty exposed and unexposed signatures (left), and bar plots with collapsed poverty exposed and unexposed signatures in 6 randomly sampled patients per group (see methods).

To further characterize these clusters, we performed marker gene and geneset enrichment analysis, comparing clusters enriched in poverty-exposed to those enriched in unexposed specimens. Notably, clusters enriched in poverty-exposed leukemic cells (clusters 3, 4, 5 and 7) demonstrated expression of genes with known relation to treatment response in B-ALL, including genes associated with hematopoietic progenitors (*FOXO1, SMARCA4, JUN, ARID5B, STAG3* and *NPY*), resistance to vincristine (*SOX11, DSC2, KCNN1 ABHD3*, and *MTF2*), and glucocorticoid therapy (*SSPB2, PCNA, IGLL1, ADGRAA3, STMN1* and *PCCA*). In contrast, clusters enriched in unexposed leukemic cells (clusters 1, 2, and 6) were characterized by expression of signatures related to metabolism (*PSAP, COTL2, LOW, CYBB, GLIPR1, ALDH3B1, MTHFD1*) and immune activation (*CD74, ISG20, GBP4*, and *PSME2*) (Figure 2E, Table S2). These analyses reveal distinct heterogenous signatures related to inflammation, metabolism and treatment resistance differentially expressed in leukemic cells from poverty-exposed versus unexposed patients.

We next evaluated whether these signatures might also be differentially expressed in bulk RNA-seq of leukemic cells from poverty-exposed and unexposed patients. To this end, we generated bulk RNA-seq data of the same 7 poverty-exposed and 7 unexposed patients on whom we had performed single-cell transcriptome analysis. We collapsed expressed genes of all signatures enriched in poverty-exposed single-cell clusters or unexposed single-cell clusters, respectively, and computed an average expression score (see Methods; supplemental Table S3). Indeed, poverty-exposed signatures were enriched in the poverty-exposed cohort compared to the unexposed cohort, whereas the unexposed signatures were enriched in the unexposed compared to the poverty-exposed cohort (Figure 2F). We confirmed this trend using CIBERSORTx^31^ for deconvolution. Although poverty-exposed signatures were detected in all patients, they were enriched in the poverty-exposed cohort, whereas unexposed signatures were enriched in the unexposed cohort. (Figure S6). We further investigated expression of the poverty-exposed versus unexposed signature within patients who had ETV6 translocated or hyperdiploid B-ALL, respectively. Expression of the poverty-exposed signature was significantly higher in poverty-exposed patients within both groups (Figure S7). We next investigated how these collapsed signatures performed in an independent dataset from our institution. To this end, we analyzed public bulk RNA-seq data downloaded from GSE181157^32^ for which we had poverty-exposure annotation (Supplemental Figure S8, Table S5). Notably, the collapsed poverty-exposed signature was anti-correlated with the unexposed signature and higher expressed in poverty-exposed patients compared to unexposed patients and vice versa (Figure 2G).

Taken together, these analyses suggest that leukemia-intrinsic transcriptional circuits differ in diagnostic B-lymphoblasts from poverty-exposed children compared to non-poverty-exposed children with enrichment of signatures associated with drug resistance in poverty-exposed children.

### Modulation of myeloid microenvironment in poverty-exposed children with B-ALL

Chronic inflammation has been implicated in adverse outcomes associated with low-socioeconomic status in several childhood diseases, including asthma and obesity.^33 34^ To investigate whether the observed differences in inflammatory signatures in leukemic blasts from poverty-exposed versus unexposed B-ALL patients might be associated with changes in the immune microenvironment, we identified the composition and transcriptional landscape of the leukemia microenvironment in the same cohort of patients, focusing on B-cells, T-cells, and myeloid cells as well-defined immune cells. There was no difference in immune cell population frequency between poverty-exposed or unexposed patients (Table S4).

Non-classical monocytes expressing *CD16* (*FCGR3A*) have recently been shown to be enriched in B-ALL at time of diagnosis and at relapse.^35^ Indeed, transcriptional analysis revealed that while normal donor myeloid cells strongly express *CD14*, myeloid cells from B-ALL patients, expressed lower levels of *CD14* with concurrently higher expression of *CD16* (*FCGR3A*) (Figure 3A, left). Notably, myeloid cells from unexposed patients showed stronger expression of *CD16* (*FCGR3A*) than those from poverty-exposed patients (Figure 3A, right).

**Figure 3.**
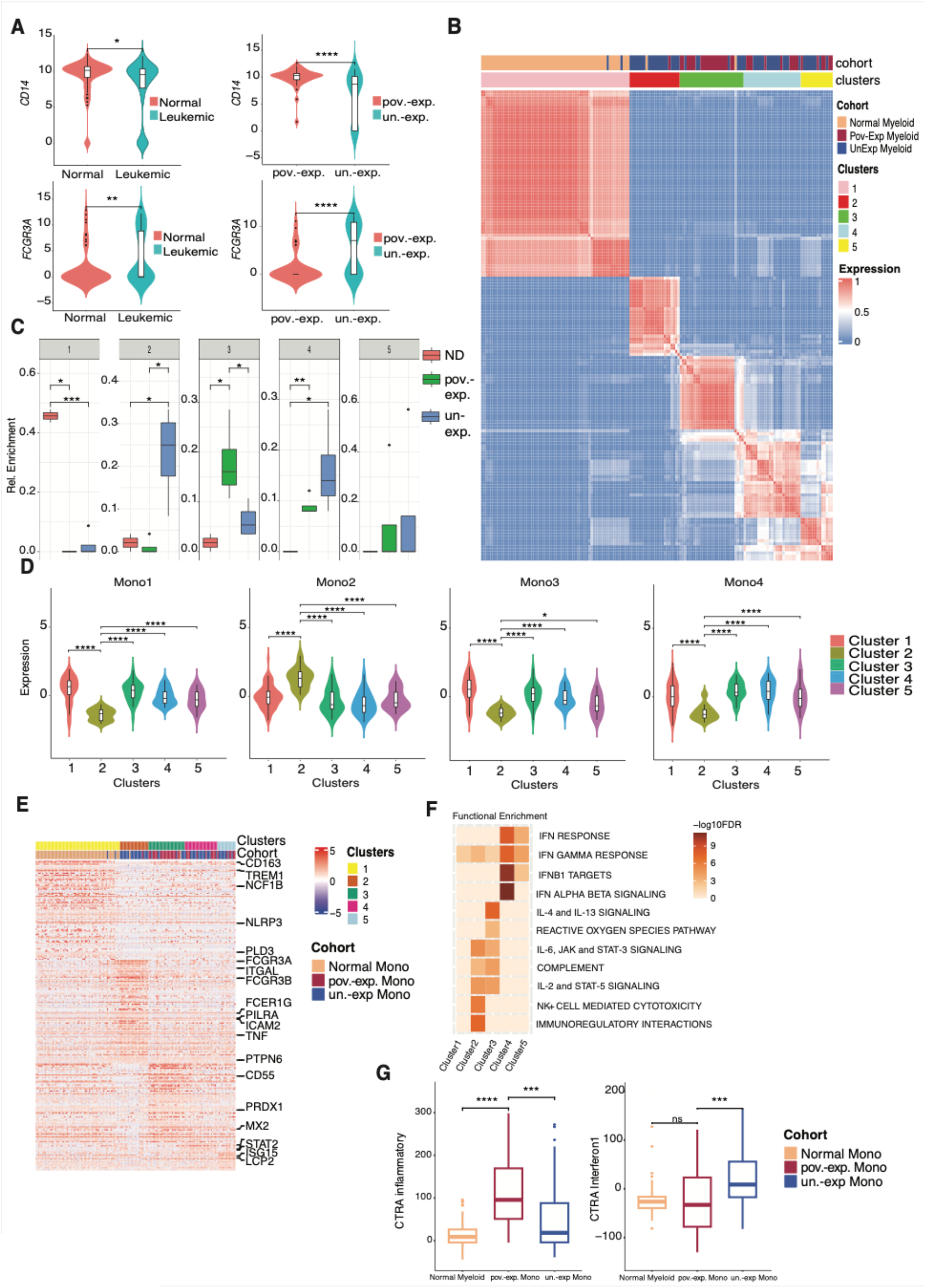
Distinct monocyte population in poverty-exposed versus unexposed patients. (A) Violin plots demonstrating expression of *CD14* and *CD16* (*FCGFR3A*) in normal donor (normal), monocytes from leukemia patients or poverty-exposed and unexposed patients, respectively (\**p*<0.05, *\*\*p*<0.01, *\*\*\*\*p*<0.0001 by two-sided Wilcoxon test). (B) Consensus clustering of myeloid cells using SC3. (C) Bar plot depicting the relative proportion of each cluster in normal donors, poverty-exposed or unexposed patients (\**p*<0.05, *\*\*p*<0.01, *\*\*\*p*<0.001 by two-sided Wilcoxon test). (D) Canonical monocyte expression signatures derived from per cluster^37^ (\**p*<0.05, *\*\*\*\*p*<0.0001 by two-sided Wilcoxon test)). (E) Heatmap depicting the expression of marker genes per cluster. (F) Functional enrichment of different clusters using FSGEA (see methods). (G) Box plots showing expression of inflammatory and interferon CTRA signatures (*\*\*\*p*<0.001 by two-sided Wilcoxon test).

To further investigate possible transcriptional differences in myeloid cells from poverty-exposed and unexposed leukemia patients, we employed SC3 clustering^36^ (see Methods) and identified 5 myeloid clusters (Figure 3B), with cluster 1 being dominated by normal donor myeloid cells (Figure 3B,C). Interestingly, while the remaining 4 clusters contained mostly myeloid cells from leukemic patients, clusters 2 and 4 were dominated by unexposed patients, whereas cluster 3 was enriched in myeloid cells from poverty-exposed patients (Figure 3B, C). To further characterize these clusters, we first looked at validated myeloid expression signatures, including expression signatures for classical monocytes (Mono1), and non-classical monocytes (Mono2).^37^ As expected, cluster 1, which contained the largest fraction of normal donor myeloid cells showed strong expression for the classical monocyte signature Mono1 (Figure 3D). Cluster 2 (enriched in unexposed patients) was dominated by expression of the non-classical monocyte signature Mono2 and likely represents non-classical monocytes that had been previously shown to be enriched in B-ALL.^35^ In contrast, clusters 3 (enriched in poverty-exposed patients) and 4 (enriched in unexposed patients) demonstrated expression of classical monocyte signatures, but in addition expressed the Mono3 signature (associated with differentiation and trafficking in cycling monocytes^38^), and Mono4 signature (associated with cytotoxicity),^37^ respectively (Figure 3D). Cluster 5, which consists of poverty-exposed and unexposed myeloid cells, did not show strong enrichment for any of the 4 monocyte signatures.

To further characterize differences between these clusters, we performed marker gene and geneset enrichment analysis (Figure 3E, F; Table S3). This analysis confirmed that genes associated with classical monocytes such as *TREM1, CD163, ALDH1A1, NLRP3, LYZ*, and *PLD3* were strongly expressed in cluster 1. Cluster 2 expressed the non-classical monocyte signature genes *CX3CR1, SIGLEC10* and *CYTIP*, and in addition expressed genes involved in inflammatory response (*FCGR3A, SAT1, GCH1, LYN*, and *LST1*), immune regulation (*LILRB2, IFITM1, PILRA, ITGAL* and *ICAM2)*, and cytotoxicity (*FCGR3B, TNF, FCER1G*, and *PTPN6)*.

Interestingly, myeloid cells in cluster 3—which were enriched in poverty-exposed patients— expressed signatures associated with cytokine signaling (*IL1B, SOCS3, TNFRSF1A, IL4R, SELL, IL4R* and *IL6R*), complement (*CD55, CLU, GCA* and *S100A12)*, and reactive oxygen species (ROS) pathway (*PRDX1, TXN* and *FES*). In contrast, cluster 4, enriched in myeloid cells from unexposed patients was enriched for genes associated with interferon alpha and beta signaling (*OAS3, OAS1, STAT2, MX2, RSAD2* and *PARP9*). Lastly, cluster 5, shared by both poverty-exposed and unexposed patients, demonstrated enrichment in interferon gamma response, but to lesser extent than cluster 4 (*LCP2, STAT1, IFI44*, and *ISG15)*.

Signaling alterations in myeloid cells have been associated with response to social adversity.^39, 40^ To evaluate whether similar mechanisms may be at play in pediatric leukemia patients we assessed expression of the conserved transcriptional response to adversity (CTRA).^41^ Indeed, we observed higher expression of the inflammatory CTRA signature in myeloid cells from patients exposed to poverty compared to unexposed which showed higher expression of the interferon CTRA signature (Figure 3G, see methods).

These findings indicate that myeloid cells from children exposed to poverty exhibit altered immune responses including expression of the inflammatory CTRA signature previously described in health adversity. This contrasts with enrichment of interferon response signatures including the interferon CTRA signatures that was expressed at higher levels in myeloid cells from B-ALL patients not exposed to poverty (Figure 3G).

### Alterations in T- and B-cell profiles among poverty-exposed children with B-ALL

We next explored differences in mature lymphoid cell composition between poverty-exposed and unexposed B-ALL patients. Overall percentages of B-cells, CD4+ and CD8+ T-cells from leukemia patients were comparable to normal donor and did not differ between poverty-exposed and unexposed patients (Table S4).

To explore if B-cell subtypes might differ between poverty-exposed and unexposed patients, we clustered B-cells using Louvain in t-SNE space and identified 5 clusters (Figure 4A, B). SingleR annotation using validated human cell atlas annotations showed that these clusters included mostly naïve, immature and memory B-cells, respectively, with clusters 2 and 4 being enriched in naïve B-cells, and clusters 1 and 5 in memory B-cells (Figure 4C). Notably, while cluster 2 representing naïve B-cells was enriched in normal donor B-cells, the remaining clusters contained B-cells from both poverty-exposed and unexposed patients (Figure 4D). Thus, representation of B-cell subsets was overall similar between poverty-exposed and unexposed patients with B-ALL.

**Figure 4.**
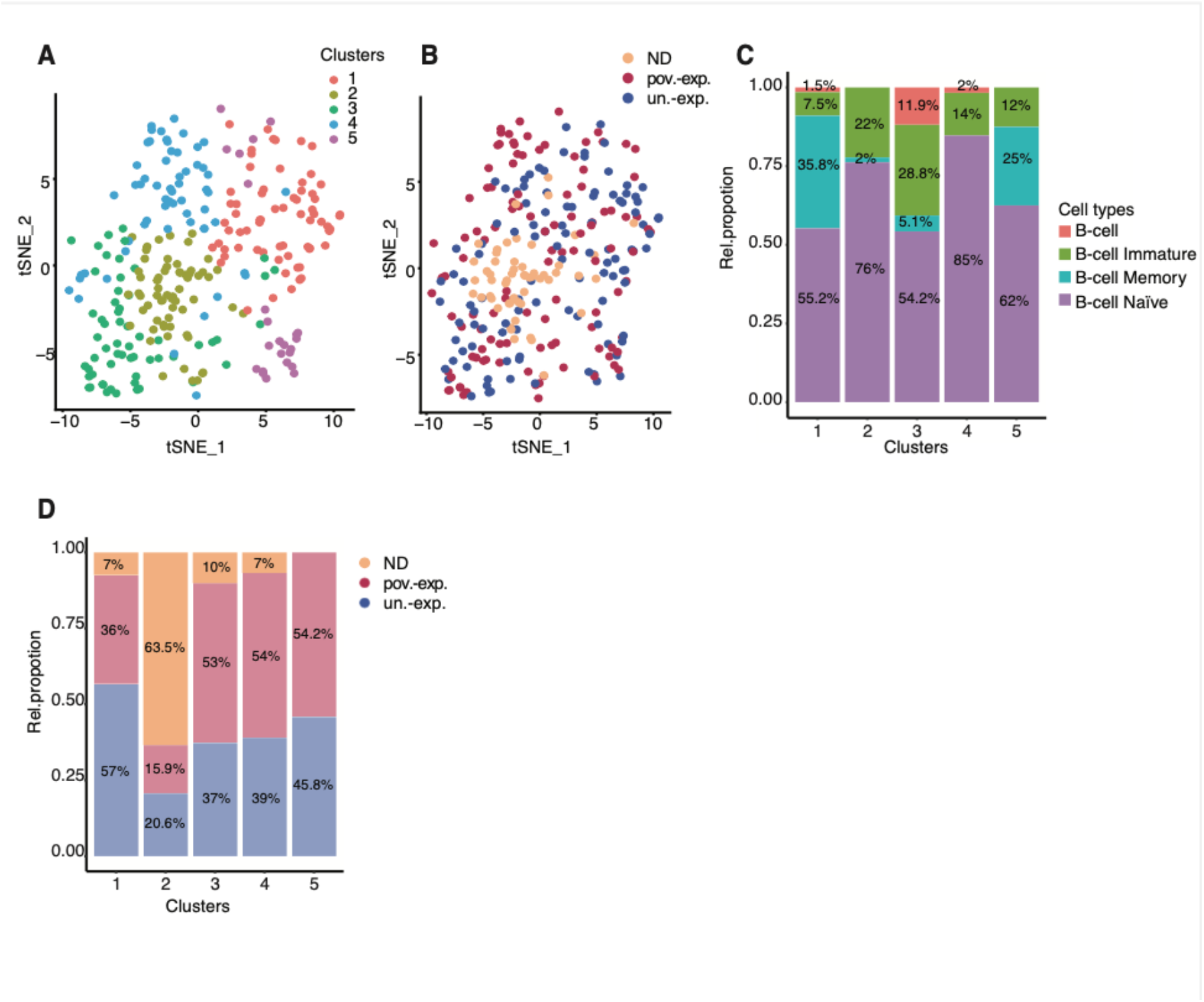
B-cell populations in poverty-exposed and unexposed children with B-ALL compared to normal donors. (A) t-SNE plot depicting 5 B-cell clusters by Louvain clustering. (B) t-SNE plot depicting distribution of B-cells from normal donors (ND), poverty-exposed (pov.-exp.) and unexposed (un-exp.) children with B-ALL. (C) Bar plot depicting relative proportions of B-cell subtypes as annotated by SingleR per cluster. (D) Bar plot depicting relative proportions of B-cells from normal donors (ND), poverty-exposed (pov.-exp.) and unexposed (un-exp.) children with B-ALL across 5 clusters.

Next, we focused on T-cells and first separated CD4+ from CD8+ T-cells using validated human cell atlas annotations (Figure 5A). We then performed Louvain clustering of CD4+ annotated T-cells and identified 4 clusters (Figure 5B, C). Further annotation using singleR and marker gene and geneset enrichment analysis demonstrated that cluster 1 was dominated by CD4+ central memory cells, clusters 2 and 3 consisted of naïve CD4+ T-cells, and cluster 4 contained effector memory CD4+ T-cells (Figure 5D, E). Notably, while normal donor T-cells were enriched in cluster 1, all CD4+ T-cell clusters contained cells from poverty-exposed and unexposed B-ALL patients. Poverty-exposed patients demonstrated slightly higher frequencies of CD4+ T-cells in clusters 2 and 3, while unexposed patients contained slightly more cells in cluster (Figure 5F). Thus, while overall percentages of CD4+ T-cells were comparable, poverty-exposed patients demonstrated relatively more naïve and central memory CD4+ T-cells and fewer effector memory CD4 T-cells (Figure S9).

**Figure 5.**
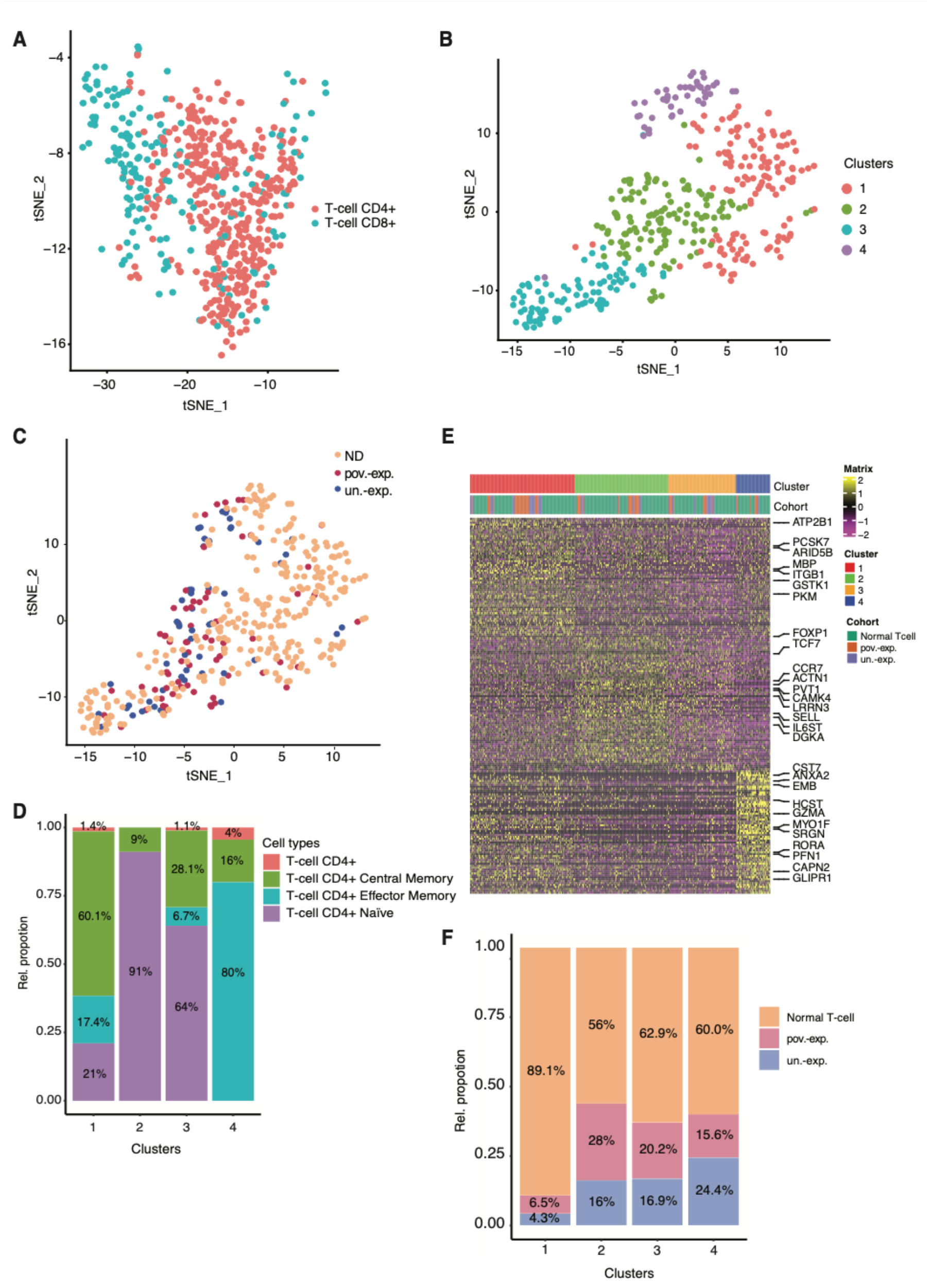
CD4+ T-cell populations in poverty-exposed and unexposed children with B-ALL compared to normal donors. (A)t-SNE plot depicting T-cells annotated as CD4+ or CD8+ T-cells, respectively, using SingleR. (B)t-SNE of CD4+ T-cells separating into 4 clusters by Louvain clustering. (C) t-SNE plot depicting distribution of CD4+ T-cells from normal donors (ND), poverty-exposed (pov.-exp.) and unexposed (un-exp.) children with B-ALL. (D) Bar plot depicting relative proportions of CD4+ T-cell subtypes as annotated by SingleR per cluster (E) Heatmap depicting expression of marker genes of CD4+ T-cells per cluster. (F) Bar plot depicting relative proportions of CD4+ T-cells from normal donors (ND), poverty-exposed (pov.-exp.) and unexposed (un-exp.) children with B-ALL across 4 clusters.

CD8+ T-cells separated into 2 clusters by Louvain, with cluster 1 containing slightly higher frequencies of CD8+ T-cells from unexposed patients, and cluster 2 being enriched in CD8+ T-cells from poverty-exposed patients (Figure 6A, B). By marker gene analysis cluster 2 cells expressed genes associated with naïve T-cells, such as *CCR7, TCF7, LEF1* and *SELL*. In contrast, cells in cluster 1 expressed gene signatures associated with T-cell dysfunction including the T-cell exhaustion marker *TIGIT* and *IL10RA*, in addition to containing genes reflecting T-cell activation, such as *GZMA, GMZB* and *PRF1* (Figure 6C). To further investigate whether endogenous CD8+ T-cells might show differences in T-cell exhaustion, we computed the average expression of genes that had been previously validated as expressed in naïve, cytotoxic, and dysfunctional T-cells (see Methods). Interestingly, while CD8+ T-cells from unexposed patients expressed signature scores at similar levels as normal donor T-cells, T-cells from poverty-exposed patients showed relatively higher expression of naïve and dysfunctional T-cell signatures (Figure 6D). Thus, these analyses support an imbalance of naïve, activated, and dysfunctional T-cells in poverty-exposed patients with B-ALL and suggest that anti-tumor immune responses may be impaired.

**Figure 6.**
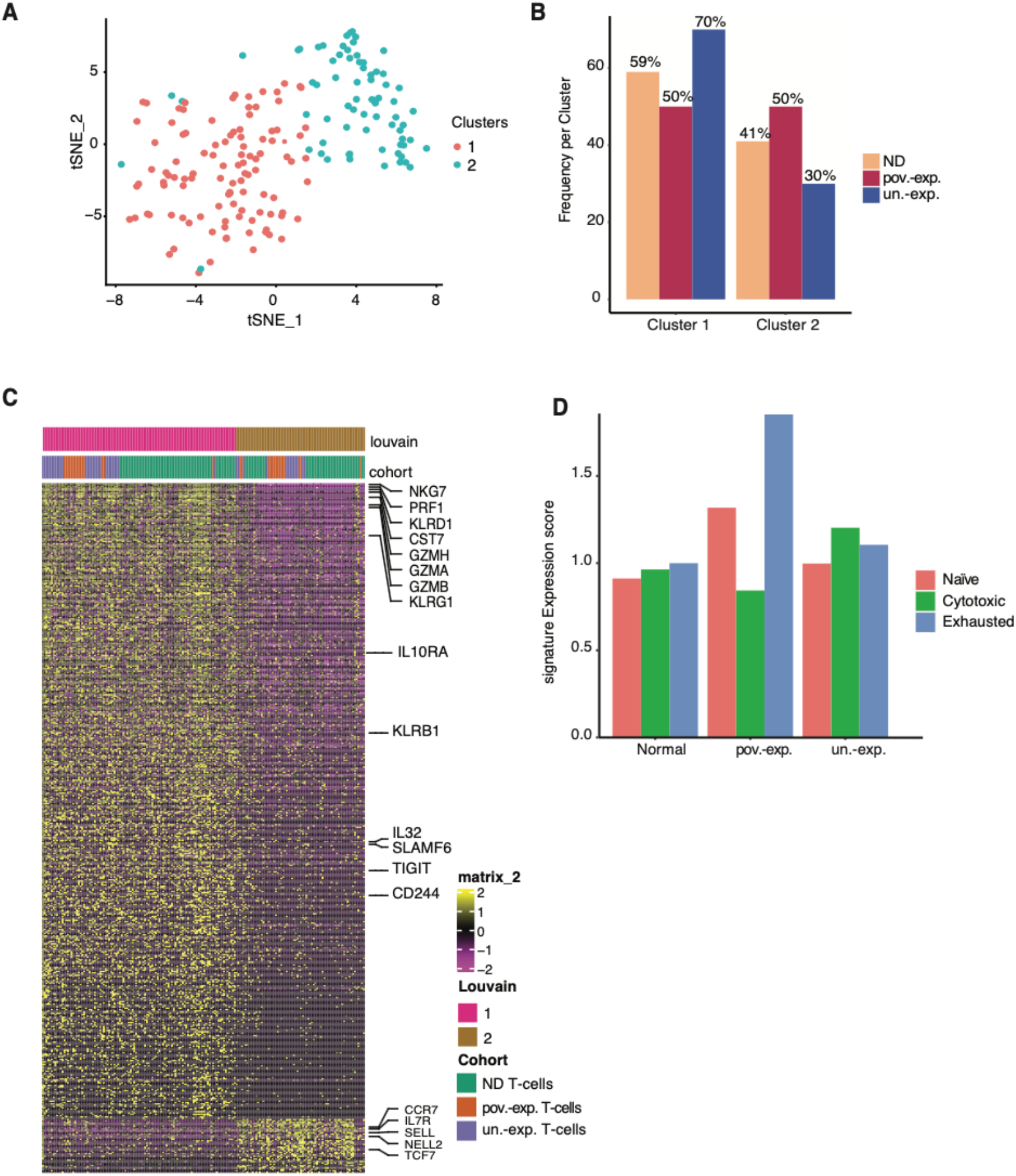
Distinct CD8+ T-cell subpopulations in poverty-exposed compared to unexposed children with B-ALL. (A) t-SNE plot of CD8+ T-cells separating into 2 clusters by Louvain clustering. (B) Bar plot depicting relative proportions of CD8+ T-cells from normal donors (ND), poverty-exposed (pov.-exp.) and unexposed (un-exp.) children with B-ALL across 2 clusters. (C) Heatmap depicting marker gene expression of CD8+ T-cells per cluster. (D) Gene expression signature score for naïve, cytotoxic and exhaustion CD8+ T-cells in poverty-exposed (pov.-exp.) and unexposed (un-exp.) children with B-ALL compared to normal donor (see methods).

## DISCUSSION

Using single-cell RNA-seq of leukemic blasts and their immune microenvironment in a cohort of children with B-ALL, we identified several differentially enriched pathways in leukemia cells from poverty-exposed and unexposed patients including steroid responsiveness and signatures consistent with altered immune responses in the leukemia microenvironment that may be associated with treatment resistance.

These data point toward several potential mechanisms of adverse treatment response that may explain observed disparities in relapse and survival among children with B-ALL living in poverty. First, our data suggest that poverty-exposed patients with B-ALL have fewer leukemia cells susceptible to steroids at diagnosis. Steroids are a core component of ALL chemotherapy and poor response to steroids is a well-accepted mechanism of treatment failure in ALL.^42^ Poverty exposure is associated with “toxic stress,” defined by the National Research Council as the excessive activation of stress regulatory systems in the absence of adequate buffering.^12^ Toxic stress physiology includes dysregulation of the hypothalamic-pituitary-adrenocortical axis and the sympathetic-adrenomedullary system that result in perturbations in stress hormones, elevated inflammatory cytokine responses, and altered cellular metabolism.^12 14 43^ It is intriguing to speculate that elevated levels of endogenous cortisol may potentially contribute to inherent steroid resistance of leukemic blasts. Notably, prior studies in adult acute myeloid leukemia patients with low SES undergoing hematopoietic stem cell transplantation have demonstrated higher expression of glucocorticoid receptor signaling signatures in PBMCs pre-transplant compared to non-exposed patients.

Prior studies have demonstrated the coexistence of toxic stress and chronic inflammation in asthma, cardiovascular disease, and depression, among other chronic diseases,^8 11 12^ and it is unknown how this might affect the development of cancer or its immune surveillance. Related, prior work in solid tumor mouse-models suggest that interventions targeting adrenergic receptor responses of the microenvironment—such as beta-blockade—may impact chemotherapy responsiveness.

Inflammatory signatures in tumor cells, including leukemias, have been linked to inferior treatment response—hypothesized to be due in part to immune dysfunction in the tumor microenvironment that prevents a host-immune response against the tumor.^44 45^ Mechanisms of immune dysfunction are well established in many solid tumors and emerging data suggest an important role for aberrant immune activation in ALL.^18 44^ There is sparse knowledge with regards to the composition and functional characteristics of the tumor immune microenvironment among patients who experience poverty and other adverse social determinants of health. Our results point towards unique immune derangements that occur in children with B-ALL in the context of poverty exposures, notably an increased expression of inflammatory signatures in myeloid cells and a relative depletion of CD8 effector cells. Importantly, myeloid cells from poverty-exposed B-ALL patients also express inflammatory signatures that have previously been associated with adversity in other disease contexts,^39, 40^ arguing that these alterations might represent universal changes that occur in the setting of hardship and may offer unique therapeutic possibilities. For example, interventions targeting adrenergic receptor responses through beta-blockade have been shown to reduce chemotherapy resistance in select solid tumors^46^, yet possible effects on the immune microenvironment have not been explored.

Our data are limited by a small sample size, including 7 children in each cohort, and thus exploratory in nature. Future replication of these findings in a larger cohort is necessary and will allow investigation of poverty-associated signatures stratified by genetic sub-type that is not possible in our sample. More detailed studies are needed to better understand the relative contributions of myeloid cells and CD8-T-cells and their interactions with leukemic cells in the context of poverty-exposure employing additional single-cell sequencing approaches that allow profiling of a larger number of cells such as the multimodal approaches available on the 10X platform. Additionally, further studies are needed to determine if the observed leukemia-intrinsic and microenvironmental alterations are limited to children with B-ALL or if they also occur in the setting of genomic high-risk ALL and their relevance for other pediatric cancers.

It is notable that at least 1 in 5 children with ALL lives in poverty at the time of diagnosis, and these children disproportionately experience treatment failure in the form of early relapse of their disease.^3^ Future work to unravel tumor microenvironmental and epigenetic state changes associated with poverty-exposure may identify opportunities for therapeutic targeting and resistance-adapted chemotherapy relevant to a strikingly high proportion of children with ALL.

## ACKNOWLEDGEMENTS

We thank the Flow Cytometry Core at Dana-Farber Cancer Institute for their assistance, the Pediatric Oncology Outcomes research team and the Clinical Research Coordinators ALL study team at Dana-Farber Cancer Institute for their help with primary patient samples. This work was supported by a Helen Trailblazers award from Dana-Farber Cancer Institute/Pussycat Foundation (to K.B, B.K.), and a National Cancer Institute – DF/HCC Nodal Award (2P30CA006516-52) (to K.B., B.K).

## AUTHOR CONTRIBUTIONS

K.B. and B.K. conceptualized this study. A.G, N.S., K.B. and B.K. co-wrote the manuscript. A.G. performed experiments and analyzed the data. N.S. performed data analysis. S.P, T.V, M.N, V.D. and J.F. executed and/or analyzed the experiments. Y.P, M.H, A.E.P, L.B.S, and J.G.L provided scientific input for study design and/or data analysis. All authors reviewed and edited the manuscript.

## REFERENCES

1. Bona K, London WB, Guo D, Frank DA, Wolfe J. Trajectory of Material Hardship and Income Poverty in Families of Children Undergoing Chemotherapy: A Prospective Cohort Study. Pediatr Blood Cancer. 2016;63(1):105–111. doi:10.1002/pbc.25762

2. Gupta S, Dai Y, Chen Z, et al. Racial and ethnic disparities in childhood and young adult acute lymphocytic leukaemia: secondary analyses of eight Children’s Oncology Group cohort trials. Lancet Haematol. 2023;10(2):e129–e141. doi:10.1016/S2352-3026(22)00371-4

3. Bona K, Blonquist TM, Neuberg DS, Silverman LB, Wolfe J. Impact of Socioeconomic Status on Timing of Relapse and Overall Survival for Children Treated on Dana-Farber Cancer Institute ALL Consortium Protocols (2000-2010). Pediatr Blood Cancer. 2016;63(6):1012–1018. doi:10.1002/pbc.25928

4. Inaba H, Pui CH. Advances in the Diagnosis and Treatment of Pediatric Acute Lymphoblastic Leukemia. J Clin Med. 2021;10(9):1926. doi:10.3390/jcm10091926

5. Ward E, DeSantis C, Robbins A, Kohler B, Jemal A. Childhood and adolescent cancer statistics, 2014. CA Cancer J Clin. 2014;64(2):83–103. doi:10.3322/caac.21219

6. Hunger SP, Raetz EA. How I treat relapsed acute lymphoblastic leukemia in the pediatric population. Blood. 2020;136(16):1803–1812. doi:10.1182/blood.2019004043

7. Vrooman LM, Blonquist TM, Stevenson KE, et al. Efficacy and Toxicity of Pegaspargase and Calaspargase Pegol in Childhood Acute Lymphoblastic Leukemia: Results of DFCI 11-001. JCO. 2021;39(31):3496–3505. doi:10.1200/JCO.20.03692

8. Singh GK, Jemal A. Socioeconomic and Racial/Ethnic Disparities in Cancer Mortality, Incidence, and Survival in the United States, 1950-2014: Over Six Decades of Changing Patterns and Widening Inequalities. J Environ Public Health. 2017;2017:2819372. doi:10.1155/2017/2819372

9. Chen E, Miller GE. Socioeconomic status and health: mediating and moderating factors. Annu Rev Clin Psychol. 2013;9:723–749. doi:10.1146/annurev-clinpsy-050212-185634

10. Adler NE, Newman K. Socioeconomic disparities in health: pathways and policies. Health Aff (Millwood). 2002;21(2):60–76. doi:10.1377/hlthaff.21.2.60

11. Braveman P, Gottlieb L. The social determinants of health: it’s time to consider the causes of the causes. Public Health Rep. 2014;129 Suppl 2:19–31. doi:10.1177/00333549141291S206

12. Seeman M, Stein Merkin S, Karlamangla A, Koretz B, Seeman T. Social status and biological dysregulation: the ‘status syndrome’ and allostatic load. Soc Sci Med. 2014;118:143–151. doi:10.1016/j.socscimed.2014.08.002

13. Shonkoff JP. Leveraging the biology of adversity to address the roots of disparities in health and development. Proc Natl Acad Sci U S A. 2012;109 Suppl 2(Suppl 2):17302–17307. doi:10.1073/pnas.1121259109

14. Blair C, Raver CC. Poverty, Stress, and Brain Development: New Directions for Prevention and Intervention. Acad Pediatr. 2016;16(3 Suppl):S30–36. doi:10.1016/j.acap.2016.01.010

15. Schiavone S, Colaianna M, Curtis L. Impact of early life stress on the pathogenesis of mental disorders: relation to brain oxidative stress. Curr Pharm Des. 2015;21(11):1404–1412. doi:10.2174/1381612821666150105143358

16. Lupien SJ, McEwen BS, Gunnar MR, Heim C. Effects of stress throughout the lifespan on the brain, behaviour and cognition. Nat Rev Neurosci. 2009;10(6):434–445. doi:10.1038/nrn2639

17. van Galen P, Hovestadt V, Wadsworth M, et al. Single-cell RNA-seq reveals AML hierarchies relevant to disease progression and immunity. Cell. 2019;176(6):1265-1281.e24. doi:10.1016/j.cell.2019.01.031

18. Anand P, Guillaumet-Adkins A, Dimitrova V, et al. Single-cell RNA-seq reveals developmental plasticity with coexisting oncogenic states and immune evasion programs in ETP-ALL. Blood. 2021;137(18):2463–2480. doi:10.1182/blood.2019004547

19. Picelli S, Faridani OR, Björklund AK, Winberg G, Sagasser S, Sandberg R. Full-length RNA-seq from single cells using Smart-seq2. Nat Protoc. 2014;9(1):171–181. doi:10.1038/nprot.2014.006

20. Frede J, Anand P, Sotudeh N, et al. Dynamic transcriptional reprogramming leads to immunotherapeutic vulnerabilities in myeloma. Nat Cell Biol. 2021;23(11):1199–1211. doi:10.1038/s41556-021-00766-y

21. Caron M, St-Onge P, Sontag T, et al. Single-cell analysis of childhood leukemia reveals a link between developmental states and ribosomal protein expression as a source of intra-individual heterogeneity. Sci Rep. 2020;10(1):8079. doi:10.1038/s41598-020-64929-x

22. Kobak D, Berens P. The art of using t-SNE for single-cell transcriptomics. Nat Commun. 2019;10(1):5416. doi:10.1038/s41467-019-13056-x

23. Aran D, Looney AP, Liu L, et al. Reference-based analysis of lung single-cell sequencing reveals a transitional profibrotic macrophage. Nat Immunol. 2019;20(2):163–172. doi:10.1038/s41590-018-0276-y

24. Fernández JM, de la Torre V, Richardson D, et al. The BLUEPRINT Data Analysis Portal. Cell Syst. 2016;3(5):491-495.e5. doi:10.1016/j.cels.2016.10.021

25. Hay SB, Ferchen K, Chetal K, Grimes HL, Salomonis N. The Human Cell Atlas bone marrow single-cell interactive web portal. Exp Hematol. 2018;68:51–61. doi:10.1016/j.exphem.2018.09.004

26. Regev A, Teichmann SA, Lander ES, et al. The Human Cell Atlas. eLife. 6:e27041. doi:10.7554/eLife.27041

27. Ramezani-Rad P, Geng H, Hurtz C, et al. SOX4 enables oncogenic survival signals in acute lymphoblastic leukemia. Blood. 2013;121(1):148–155. doi:10.1182/blood-2012-05-428938

28. Wu Z, Zhang F, Liu C, et al. Whole transcriptome sequencing reveals a TCF4-ZNF384 fusion in acute lymphoblastic leukemia. Frontiers in Oncology. 2022;12. Accessed July 13, 2023. https://www.frontiersin.org/articles/10.3389/fonc.2022.900054

29. Zhang J, McCastlain K, Yoshihara H, et al. Deregulation of DUX4 and ERG in acute lymphoblastic leukemia. Nat Genet. 2016;48(12):1481–1489. doi:10.1038/ng.3691

30. Blondel VD, Guillaume JL, Lambiotte R, Lefebvre E. Fast unfolding of communities in large networks. J Stat Mech. 2008;2008(10):P10008. doi:10.1088/1742-5468/2008/10/P10008

31. Newman AM, Steen CB, Liu CL, et al. Determining cell type abundance and expression from bulk tissues with digital cytometry. Nat Biotechnol. 2019;37(7):773–782. doi:10.1038/s41587-019-0114-2

32. Tran TH, Langlois S, Meloche C, et al. Whole-transcriptome analysis in acute lymphoblastic leukemia: a report from the DFCI ALL Consortium Protocol 16-001. Blood Adv. 2022;6(4):1329–1341. doi:10.1182/bloodadvances.2021005634

33. Trivedi M, Denton E. Asthma in Children and Adults—What Are the Differences and What Can They Tell us About Asthma? Front Pediatr. 2019;7:256. doi:10.3389/fped.2019.00256

34. Furman D, Campisi J, Verdin E, et al. Chronic inflammation in the etiology of disease across the life span. Nat Med. 2019;25(12):1822–1832. doi:10.1038/s41591-019-0675-0

35. Witkowski MT, Dolgalev I, Evensen NA, et al. Extensive Remodeling of the Immune Microenvironment in B Cell Acute Lymphoblastic Leukemia. Cancer Cell. 2020;37(6):867-882.e12. doi:10.1016/j.ccell.2020.04.015

36. Kiselev VY, Kirschner K, Schaub MT, et al. SC3: consensus clustering of single-cell RNA-seq data. Nat Methods. 2017;14(5):483–486. doi:10.1038/nmeth.4236

37. Villani AC, Satija R, Reynolds G, et al. Single-cell RNA-seq reveals new types of human blood dendritic cells, monocytes, and progenitors. Science. 2017;356(6335):eaah4573. doi:10.1126/science.aah4573

38. Olingy CE, Dinh HQ, Hedrick CC. Monocyte heterogeneity and functions in cancer. J Leukoc Biol. 2019;106(2):309–322. doi:10.1002/JLB.4RI0818-311R

39. Taylor MR, Cole SW, Strom J, et al. Unfavorable transcriptome profiles and social disadvantage in hematopoietic cell transplantation: a CIBMTR analysis. Blood Adv. 2023;7(22):6830–6838. doi:10.1182/bloodadvances.2023010746

40. Cole SW. The Conserved Transcriptional Response to Adversity. Curr Opin Behav Sci. 2019;28:31–37. doi:10.1016/j.cobeha.2019.01.008

41. Knight JM, Rizzo JD, Hari P, et al. Propranolol inhibits molecular risk markers in HCT recipients: a phase 2 randomized controlled biomarker trial. Blood Adv. 2020;4(3):467–476. doi:10.1182/bloodadvances.2019000765

42. Pui CH, Yang JJ, Hunger SP, et al. Childhood Acute Lymphoblastic Leukemia: Progress Through Collaboration. J Clin Oncol. 2015;33(27):2938–2948. doi:10.1200/JCO.2014.59.1636

43. Johnson SB, Riley AW, Granger DA, Riis J. The science of early life toxic stress for pediatric practice and advocacy. Pediatrics. 2013;131(2):319–327. doi:10.1542/peds.2012-0469

44. Witkowski MT, Kousteni S, Aifantis I. Mapping and targeting of the leukemic microenvironment. J Exp Med. 2020;217(2):e20190589. doi:10.1084/jem.20190589

45. Chu Y, Dai E, Li Y, et al. Pan-cancer T cell atlas links a cellular stress response state to immunotherapy resistance. Nat Med. 2023;29(6):1550–1562. doi:10.1038/s41591-023-02371-y

46. Manoleras AV, Sloan EK, Chang A. The sympathetic nervous system shapes the tumor microenvironment to impair chemotherapy response. Front Oncol. 2024;14:1460493. doi:10.3389/fonc.2024.1460493

