## Supplemental for "Altered immune and treatment response gene expression signatures among poverty-exposed children with B-ALL"

**SUPPLEMENTAL MATERIALS**

**METHODS**

**Primary samples and data**

Banked diagnostic bone marrow samples from children with *de novo* B-cell ALL diagnosed between 2005 and 2018 who had consented to biobanking for future research (Dana-Farber Cancer Institute (DFCI) protocols 06-078, 05-001 [NCT00400946], 11-001 [NCT01574274]) were utilized in this analysis. Parent-reported socioeconomic data from concurrent poverty-focused research protocols (DFCI 11-198 and 16-544) were utilized to annotate biospecimens. Patient demographics and tumor characteristics (immunophenotype, cytogenetics, and targeted DNA sequencing results) were abstracted from study protocol data and the electronic medical record. This study was approved by the DFCI Institutional Review Board.

**Poverty-exposure**

Poverty was the primary exposure of interest and was defined *a priori* at the household-level utilizing both income-poverty and unmet resource needs. Specifically, patients were considered poverty-exposed if they had concurrent parent reported low-income (annual household income <200% of the Federal Poverty Level[1] (FPL) and household material hardship (HMH; food, housing, or transportation insecurity)[1] at time of diagnosis. Those with reported income >200% FPL were considered unexposed, regardless of whether HMH data were available.

**Cohort assembly**

To minimize heterogeneity, we excluded biospecimens from patients with high-risk immunophenotypic or genetic abnormalities at diagnosis including T-cell immunophenotype, *KMT2A* or *BCR-ABL1* gene rearrangements, or high-risk cytogenetic abnormalities (low hypodiploidy; t[17;19], iAMP21). Analytic specimens were thus restricted to lymphoblasts with B-cell immunophenotype, *ETV6-RUNX1* translocations or a hyperdiploid karyotype—two genetic features frequently associated with standard risk prognosis—or normal FISH and cytogenetic testing. Biospecimens were analyzed in a deidentified fashion.

**Flow cytometry**

Mononuclear cells (PBMCs) were isolated using Ficoll-Paque PLUS from GE Healthcare according to manufacturer's protocol. B-ALL samples were stained for CD45 FITC (Thermo Fisher Scientific Cat# 11-0459-41, RRID: AB_10854279), T-cells for CD3 PerCP-Cy5.5 (Thermo Fisher Scientific Cat# 45-0037-41, RRID:AB_10548354), monocytes for CD14 APC-Cy7 (BD Biosciences Cat# 557831, RRID:AB_396889), B-cells for CD19 PE (BioLegend Cat# 302208, RRID:AB_314238). Cells were stained with DAPI (1 µg/mL, Sigma Aldrich) to exclude dead cells. Single-cells were sorted into 96 well plates containing TCL buffer (Qiagen) using a Sony SH800 sorter. Minipools of 100 leukemic cells from each patient were sorted in the same way and used for bulk RNA-seq. Number of sorted and sequenced cells for each population are listed in Supplemental Table 4.

**scRNA-seq library preparation and sequencing**

Full-length single-cell RNA-seq libraries were prepared using the SMART-seq2 protocol.[2] RNA was purified by RNAClean XP beads (Beckman Coulter) and reverse transcribed using Maxima RNAse H-minus (Thermo Fisher Scientific) in the presence of oligo-dT30VN, template-switching oligonucleotides, and betaine. cDNA was then amplified using the KAPA Hifi Hotstart ReadyMix (Kappa Biosystems) and ISPCR primers with the following protocol: 98°C - 3 min, 24 x [98°C - 15s, 67°C - 20s, 72°C – 6min], 72°C - 5 min. After purification with Agencourt Ampure XP beads (Beckmann Coulter), product size distribution was assessed on a Bioanalyzer using a High

Sensitivity DNA Kit (Agilent Technologies). The quantity of amplified cDNA was determined by Qubit (Thermo Fisher Scientific). 0.15 ng of the product was fragmented and indexed using Nextera XT and Nextera PCR primers (Illumina). Following another purification step, quality control was assessed by Bioanalyzer High Sensitivity DNA Kit. Libraries were pooled and quantified using Qubit dsDNA High Sensitivity reagents. Paired-end sequencing was performed on a NextSeq 500 (Illumina) with an aimed sequencing depth of 1 million reads per single-cell.

**Statistical analysis**

**Processing of scRNA-seq reads**

Following sequencing reads were trimmed using trimmomatic and aligned to the hg19 version of the genome using STAR aligner with following parameters '-- twopassMode Basic --alignIntronMax 100000 --alignMatesGapMax 100000 --alignSJDBoverhangMin 10 --alignSJstitchMismatchNmax 5 -1 5 5'.[3,4] Raw counts and normalized TPM values were obtained from the aligned bam file using HTSeq and RSEM, respectively.[5,6]

All statistical analyses were conducted using R (version 4.0.4) and R packages available at BioConductor (https://www.bioconductor.org). To test for relationships between different groups, the stat_compare_means function from the ggpubr (Version 0.6.0) package was utilized. *P*-values were adjusted for false discovery rate (FDR) across all comparisons using Holm-Bonferroni. An adjusted *P*-value equal or lower than 0.05 and a log 2-fold change of more or equal to 0.58 was considered significant. A parametric student's t tests or One-Way ANOVA was used to compare the means of 2 groups or multiple groups, respectively. All data represent the mean ± standard deviation (SD). * $p < 0.05$; ** $p < 0.01$; *** $p < 0.001$, **** $p < 0.0001$, were considered statistically significant.

**Quality filtering of scRNA-seq data**

To filter out low quality cells from our dataset, we used four different parameters – i) library size, ii) number of genes detected, iii) percentage of reads mapping to house-keeping genes, and iv) percentage of reads mapping to mitochondrial genes (Figure S1A). QC parameters were calculated for each cell using *addPerCellQC* function from *scater.*[7]Then, outlier cells were identified for all defined metrics, based on the median absolute deviation (MAD) by calculation of the median of each metric across all samples. 1,347 single cells (from 4,513 samples) with median absolute deviations (M.A.D's) of more than 3 were identified as low quality cells (Figure S1B). Cells that were detected as outliers were removed from the dataset (Figure S1 B-C). This resulted in retention of 3,166 high-quality cells for further downstream analyses (Figure S1C). On average, around 3,618 genes were detected in 25% of cells (Figure S1C). The variance in the dataset was used to systematically investigate the contribution of various technical factors and batch effects. The proportion of variance due to cohort, total number of genes detected, percentage of housekeeping and mitochondrial genes and library size were found to be low (Figure S1D).

**Clustering of scRNA-seq profiles and identification of cell types**

For comparison, analyses included data from 719 normal donor cells (after quality control) derived from published datasets.[8,9] Clustering of high-quality cells was performed using the Louvain algorithm for t-SNE representation.[10] To rule out the possibility of clusters being purely driven by cell-cycle, each individual cell was analyzed for expression of G1, G2M and S phase markers to predict the cell-cycle phase using Seurat, [11] and cell cycle phases (S.Score, G2M.Score) were regressed out from the dataset using the Seurat ScaleData function.

Cell types were initially inferred using the Human primary cell atlas (HPCA) panel in the Bioconductor package SingleR (https://github.com/dviraran/SingleR),[12] cell identity was confirmed using the *findMarker* function in the scran package,[13] and clusters were annotated as $CD4^+$ T-cells, $CD8^+$ T-cells, NK cells, myeloid, B- or Pro-B-cells (Figure 1). Cluster 15 included marker genes from several other cell types and was therefore labelled as "not annotated" (N/A).

For Figure 2, cells were clustered based on expression of signatures for hematopoietic stem cells (HSC, 25 genes), common lymphoid progenitors (CLP, 31 genes), pro-B (169 genes), immature B (19 genes) and naïve-B cells (33 genes) that were derived from the literature (Table S1),[9] using AUCell[14] (Version 1.12.0 R-package).

For Figure 3, single-cell consensus clustering SC3[15] was used with ks set to 5.

For comparison of pediatric versus adult bone marrow composition we downloaded single cell RNA-seq data from pediatric bone marrow from GSE132509.[16,17] We used Read10x()[27] function to read and create a raw count matrix and then integrated the count matrix of the pediatric dataset with our adult raw count matrix. To annotate the cell types in the created dataset we used the Azimuth R package. Subsequently, visualization was carried out using the ggboxplot function from the ggpubr and ggplot2 packages in R.

**Detection of copy number variations from scRNA-seq and cytogenetics**

The copy number variants of individual cells were inferred from the scRNA-seq data using InferCNV (https://github.com/broadinstitute/inferCNV) using normal donor B-cells (see above) as reference.

**Gene set enrichment analysis (GSEA)**

We performed GSEA using fgsea (version 1.18.0 R-package).[18] We first annotated genes (produced by findMarker function) with Entrez ID using EnsDb.Hsapiens.v86 (version 2.99.0 R-package). Then, gene lists were ranked by log2fold (0.58) change or FDR values before subjecting to fgseaMultilevel function. FDR-adjusted $P < 0.05$ was considered significant. Genes for which there was no Entrez IDs were excluded.

**Scoring of expression signatures in bulk and single-cells**

To generate poverty-exposed versus unexposed signatures, we compiled signatures by intersecting marker genes with expressed genes of enriched GSEA signatures (Figure 2E) and scored their expression in bulk using AUCell[14] (Version 1.12.0), using aucMaxRank set to default value of 0.05. Monocyte subsets were scored employing AUCell[14] for published myeloid expression signatures[19,20,21] and published conserved signatures of adversity. Upregulated genes: *IL1A*, *IL1B, IL6, IL8/CXCL8, TNF, PTGS1, PTGS2, FOS, FOSB, FOSL1, FOSL2, JUN, JUNB, JUND, NFKB1, NFKB2, REL, RELA*, and *RELB*. Downregulated genes: *GBP1, IFI27, IFI27L1-2, IFI30, IFI35, IFI44, IFI44L, IFI6, IFIH1, IFIT1-3, IFIT5, IFIT1B, IFITM1-3, IFITM4P, IFITM5, IFNB1, IRF2, IRF7-8, MX1-2, OAS1-3, OASL, JCHAIN,* and *IGLL1*.

**T-cell activation scores**

T-cell exhaustion, naïve and cytotoxic scores were calculated using an average relative expression of key marker genes from the literature.[22]The exhaustion score was defined as difference between average relative expression of exhaustion markers – *PDCD1, TIGIT, LAG3, HAVCR2, CTLA4* and naïve markers – *CCR7, TCF7, LEF1 and SELL*. The cytotoxic score was defined as difference between average relative expression of cytotoxic markers – *NKG7, CCL4, CST7, PRF1, GZMA, GZMB, IFNG, CCL3* and naïve markers.

**Scoring relative proportion of poverty-exposed and unexposed signatures in bulk RNA-seq**

Output for bulk RNA-seq was generated in *FASTQ* format (http://www.bioinformatics.babraham.ac.uk/projects/fastqc) mapped to the human genome GRCh37 (hg19). Gene expression at transcript-level resolution was calculated using *RSEM* (v1.2.31) after alignment with STAR[3] (and duplicates marked with *Picard* (https://broadinstitute.github.io/picard/). Processed bulk RNA-seq read counts from patient samples undergoing treatment on 16-001 were obtained from GSE181157, and used to calculate correlation of the collapsed poverty-exposed and unexposed signatures.[23] We calculated correlations for gene set variation scores using the GSVA R package[24] to compute the statistical significance differences for the two gene set signatures of poverty-exposed and unexposed by transforming the gene expression into enrichment scores (Fig 2G). Given the imbalance of the two groups, we randomly sampled n=6 per group, repeating random sampling 10x (Supplemental Figure S8). To analyze relative enrichment of poverty-exposed/unexposed signatures we used the CIBERSORTx web portal (https://cibersort.stanford.edu),[25] using a custom reference matrix file.

**Data Availability Statement**
Sequencing data have been deposited in GEO (reference number GSE237888). Codes used for all analyses are available upon request from the corresponding author.

**SUPPLEMENTAL DATA**

**Figure S1. Quality filtering of scRNA-seq dataset**. (A) Distribution of features in the unfiltered dataset – (i) library size per cell, (ii) number of genes detected in each cell, (iii) percentage of counts mapping to house-keeping genes and (iv) percentage of counts mapping to mitochondrial genes in each cell in all cells sequenced. (B) The distribution of the features after filtering out the cells detected to be outlier. (C) Scatter plot depicting the quality of data from the remaining 3,166 cells through expression frequency and mean read counts per gene. (D) the density plot depicting the contribution of various technical factors contributing to the total variation observed in entire dataset.

**Figure S2. Cell Type Proportion Adult vs Pediatric.** (A) B-cell type proportions. (B) T-cell type proportions. (C) Monocyte proportions.

**Figure S3. Inferred human cell atlas annotations.** Lineage annotation of the single-cell profiles according to Human Cell Atlas (HCA) dataset.

**Figure S4. Copy Number Variations (CNV) called in single leukemia or normal cells.** The copy number variants were inferred from the scRNA-seq data using InferCNV. The detected CNVs were compared to the cytogenetics report obtain as clinical routine (Table 1).

**Figure S5. Genetic features in leukemic single cell clusters.** (A) t-SNE plot color-coded based on presence of genetic feature (hyperdiploid or ETV6-RUNX1 translocated). (B) Bar plot representing relative proportions of genetic features across the 7 leukemic clusters.

**Figure S6. Expression of poverty-exposed and unexposed signatures in bulk leukemia RNA-seq.** Signature scores of poverty-exposed and unexposed pathways using CIBERSORTx (see methods).

**Figure S7. ETV6-RUNX1 and hyperdiploid vs poverty exposed and non-poverty exposed signatures.** ETV6-RUNX1 (A) unexposed and (B) poverty exposed. Hyperdiploid (C) unexposed. (D) poverty exposed.

**Figure S8. Expression of poverty-exposed vs unexposed gene expression signatures in bulk RNA-seq.** (A) Top 5 correlation iterations between exposed vs unexposed signature. (B) Top 5 boxplots iterations depicting unexposed vs unexposed signatures in unexposed vs exposed B-ALL patients (see methods).

**Figure S9. CD4+ T-cell populations in B-ALL patients compared to normal donors.**
Bar plot of relative proportion CD4+ T-cell subtypes in poverty-exposed, unexposed B-ALL patients and normal donors.

### SUPPLEMENTAL TABLES

**Supplemental Table S1. Progenitor signatures from Human Cell Atlas (HCA).** Genes included in signatures for HSC, CLP, pro-B, immature B-cell, and naïve B-cells.

**Supplemental Table S2. Marker genes of clusters upregulated in poverty-exposed or unexposed leukemia cells**. Log2FC poverty-exposed to unexposed clusters.

**Supplemental Table S3. Poverty-exposed and unexposed signatures.** List of genes included in poverty-exposed and unexposed signature.

**Supplemental Table S4. Percentages and sorted number of leukemia and immune cells in poverty-exposed and unexposed B-ALL patients.** Flow percentages of $CD45^{low}$ leukemia cells, $CD45^{high}CD3^{+}$ T-cells, $CD45^{high}CD19^{+}$ B-cells and $CD34^{high}CD14^{+}$ myeloid cells in bone marrow from poverty-exposed and unexposed children with B-ALL at time of diagnosis.

**Supplemental Table S5. List of poverty-exposed vs unexposed B-ALL samples from GSE181157.**[23]

### REFERENCES

1. Bona K, London WB, Guo D, Frank DA, Wolfe J. Trajectory of Material Hardship and Income Poverty in Families of Children Undergoing Chemotherapy: A Prospective Cohort Study. *Pediatr Blood Cancer*. 2016;63(1):105-111. doi:10.1002/pbc.25762
2. Picelli S, Faridani OR, Björklund AK, Winberg G, Sagasser S, Sandberg R. Full-length RNA-seq from single cells using Smart-seq2. *Nat Protoc*. 2014;9(1):171-181. doi:10.1038/nprot.2014.006
3. Dobin A, Davis CA, Schlesinger F, et al. STAR: ultrafast universal RNA-seq aligner. *Bioinformatics*. 2013;29(1):15-21. doi:10.1093/bioinformatics/bts635
4. Bolger AM, Lohse M, Usadel B. Trimmomatic: a flexible trimmer for Illumina sequence data. *Bioinformatics*. 2014;30(15):2114-2120. doi:10.1093/bioinformatics/btu170
5. Anders S, Pyl PT, Huber W. HTSeq--a Python framework to work with high-throughput sequencing data. *Bioinformatics*. 2015;31(2):166-169. doi:10.1093/bioinformatics/btu638
6. Li B, Dewey CN. RSEM: accurate transcript quantification from RNA-Seq data with or without a reference genome. *BMC Bioinformatics*. 2011;12:323. doi:10.1186/1471-2105-12-323
7. McCarthy D, Campbell K, Lun A, Wills Q. *scater:*pre-processing, quality control, normalisation and visualisation of single-cell RNA-seq data in. Preprint posted online 2016. doi:10.1101/069633
8. Anand P, Guillaumet-Adkins A, Dimitrova V, et al. Single-cell RNA-seq reveals developmental plasticity with coexisting oncogenic states and immune evasion programs in ETP-ALL. *Blood*. 2021;137(18):2463-2480. doi:10.1182/blood.2019004547
9. Frede J, Anand P, Sotudeh N, et al. Dynamic transcriptional reprogramming leads to immunotherapeutic vulnerabilities in myeloma. *Nat Cell Biol*. 2021;23(11):1199-1211. doi:10.1038/s41556-021-00766-y
10. Kobak D, Berens P. The art of using t-SNE for single-cell transcriptomics. *Nat Commun*. 2019;10(1):5416. doi:10.1038/s41467-019-13056-x

11. Stuart T, Butler A, Hoffman P, et al. Comprehensive Integration of Single-Cell Data. *Cell*. 2019;177(7):1888-1902.e21. doi:10.1016/j.cell.2019.05.031
12. Aran D, Looney AP, Liu L, et al. Reference-based analysis of lung single-cell sequencing reveals a transitional profibrotic macrophage. *Nat Immunol*. 2019;20(2):163-172. doi:10.1038/s41590-018-0276-y
13. Lun ATL, McCarthy DJ, Marioni JC. A step-by-step workflow for low-level analysis of single-cell RNA-seq data with Bioconductor. *F1000Research*. Preprint posted online 31 October 2016. doi:10.12688/f1000research.9501.2
14. Aibar S, González-Blas CB, Moerman T, et al. SCENIC: single-cell regulatory network inference and clustering. *Nat Methods*. 2017;14(11):1083-1086. doi:10.1038/nmeth.4463
15. Kiselev VY, Kirschner K, Schaub MT, et al. SC3: consensus clustering of single-cell RNA-seq data. *Nat Methods*. 2017;14(5):483-486. doi:10.1038/nmeth.4236
16. Caron M, St-Onge P, Sontag T, et al. Single-cell analysis of childhood leukemia reveals a link between developmental states and ribosomal protein expression as a source of intra-individual heterogeneity. *Sci Rep*. 2020;10(1):8079. doi:10.1038/s41598-020-64929-x
17. Knight JM, Rizzo JD, Hari P, et al. Propranolol inhibits molecular risk markers in HCT recipients: a phase 2 randomized controlled biomarker trial. *Blood Adv*. 2020;4(3):467-476. doi:10.1182/bloodadvances.2019000765
18. Korotkevich G, Sukhov V, Budin N, Shpak B, Artyomov M, Sergushichev A. Fast gene set enrichment analysi. Preprint posted online 2016. doi:10.1101/060012
19. Villani AC, Satija R, Reynolds G, et al. Single-cell RNA-seq reveals new types of human blood dendritic cells, monocytes, and progenitors. *Science*. 2017;356(6335):eaah4573. doi:10.1126/science.aah4573
20. Taylor MR, Cole SW, Strom J, et al. Unfavorable transcriptome profiles and social disadvantage in hematopoietic cell transplantation: a CIBMTR analysis. *Blood Adv*. 2023;7(22):6830-6838. doi:10.1182/bloodadvances.2023010746
21. Cole SW. The Conserved Transcriptional Response to Adversity. *Curr Opin Behav Sci*. 2019;28:31-37. doi:10.1016/j.cobeha.2019.01.008
22. Heng TS, Painter MW; Immunological Genome Project Consortium. The Immunological Genome Project: networks of gene expression in immune cells. Nat Immunol. 2008 Oct;9(10):1091-4. doi: 10.1038/ni1008-1091. PMID: 18800157.
23. Tran TH, Langlois S, Meloche C, et al. Whole-transcriptome analysis in acute lymphoblastic leukemia: a report from the DFCI ALL Consortium Protocol 16-001. *Blood Adv*. 2022;6(4):1329-1341. doi:10.1182/bloodadvances.2021005634
24. Hänzelmann S, Castelo R, Guinney J. GSVA: gene set variation analysis for microarray and RNA-seq data. *BMC Bioinformatics*. 2013;14:7. doi:10.1186/1471-2105-14-7
25. Newman AM, Steen CB, Liu CL, et al. Determining cell type abundance and expression from bulk tissues with digital cytometry. *Nat Biotechnol*. 2019;37(7):773-782. doi:10.1038/s41587-019-0114-2

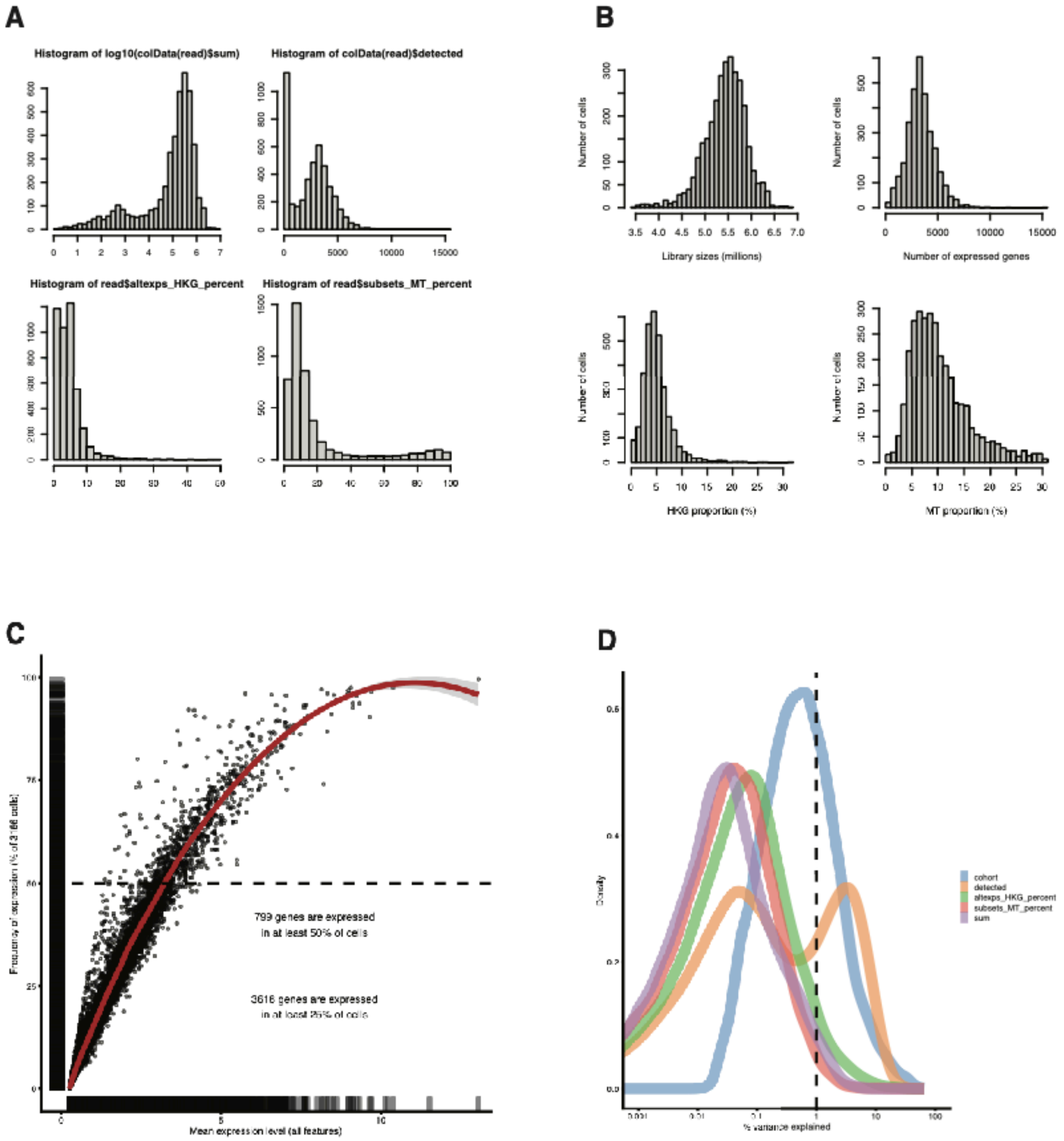


Supp Figure S1

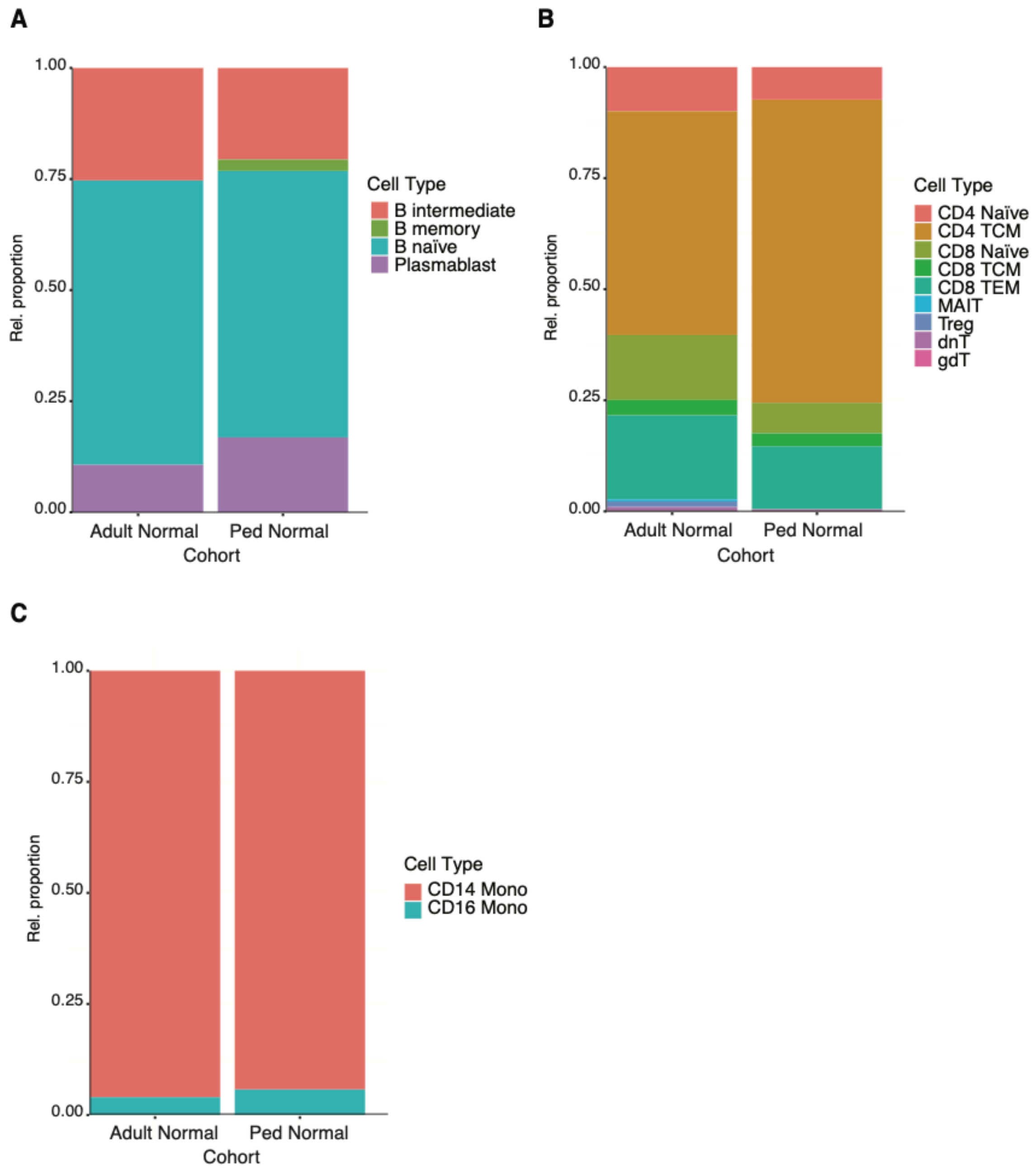


**Suppl Figure S2**

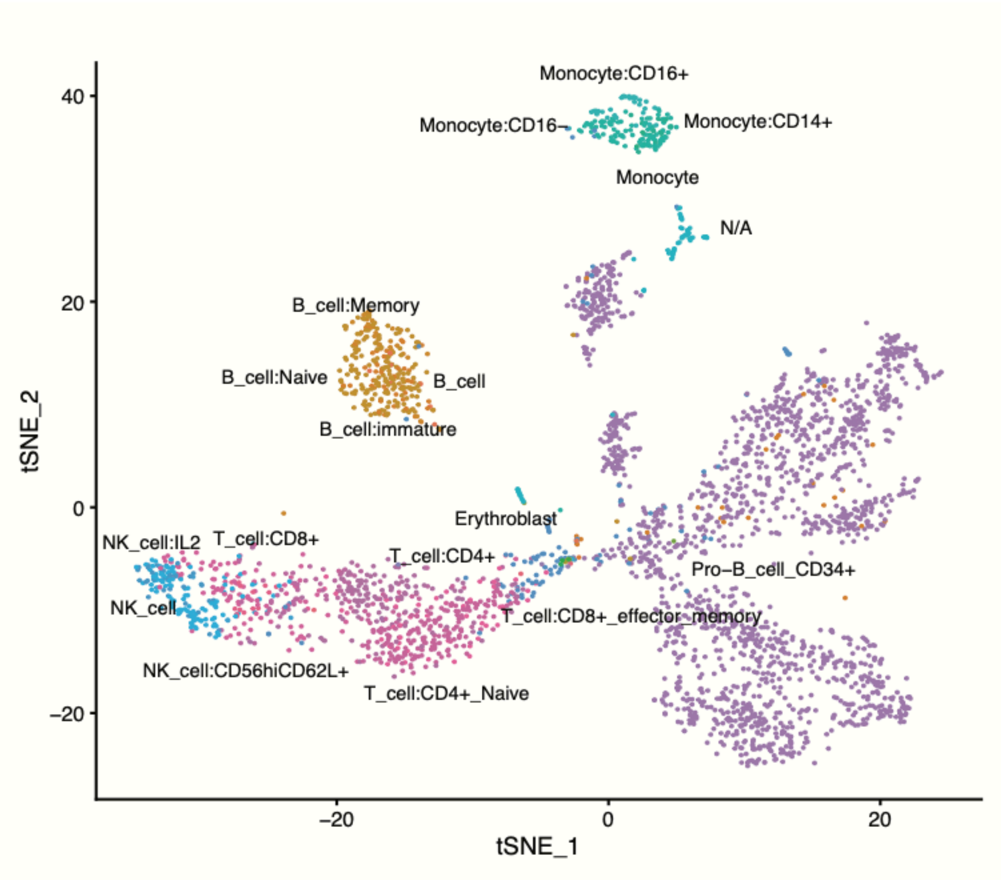


**Suppl Figure S3**

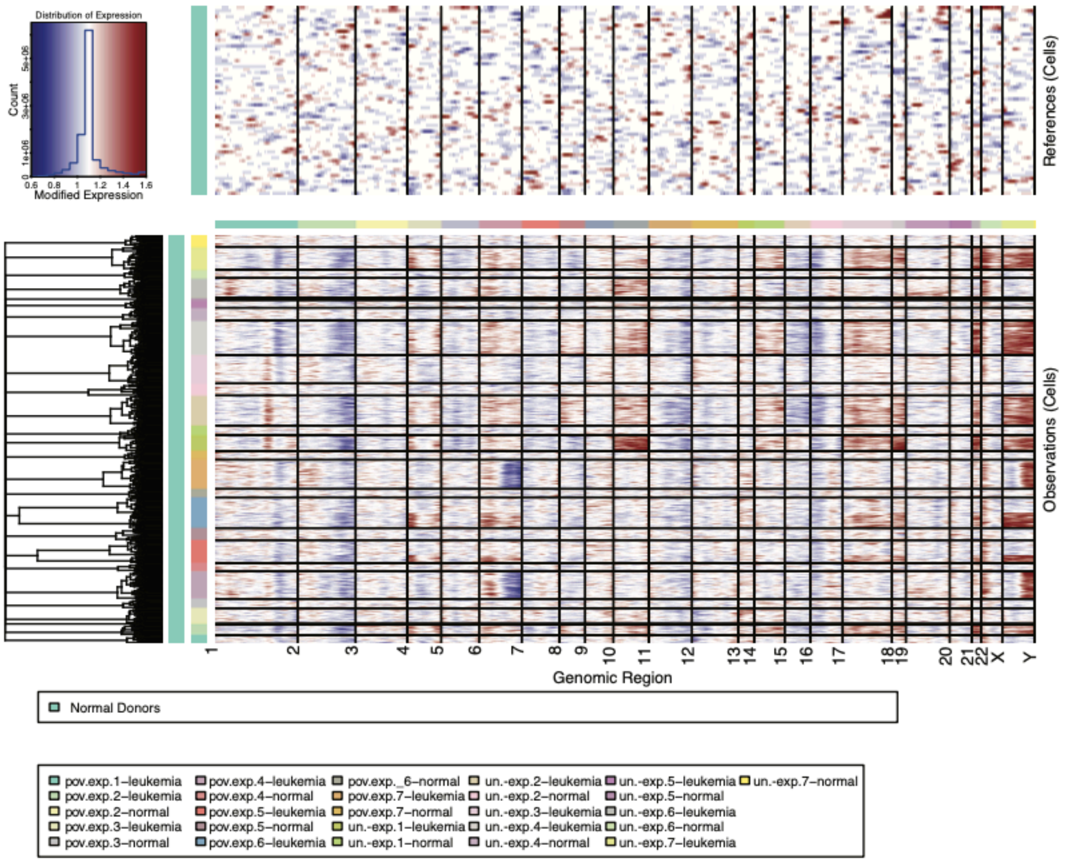


**Suppl Figure S4**

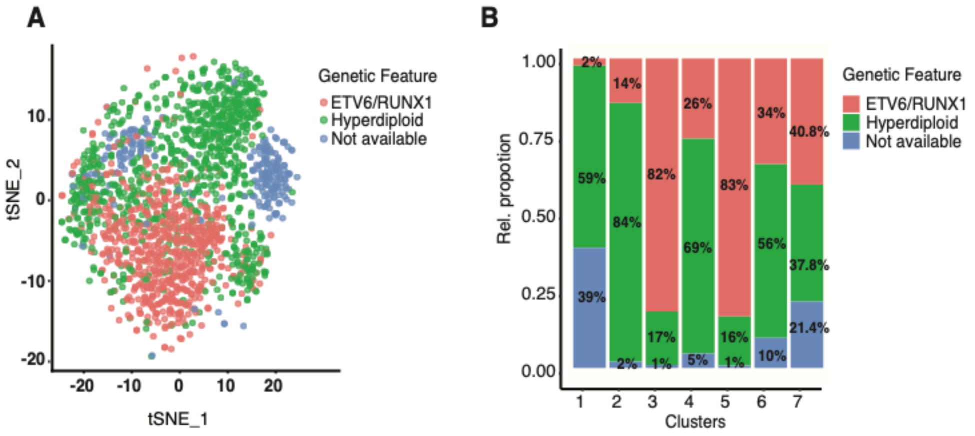


**Suppl Figure S5**

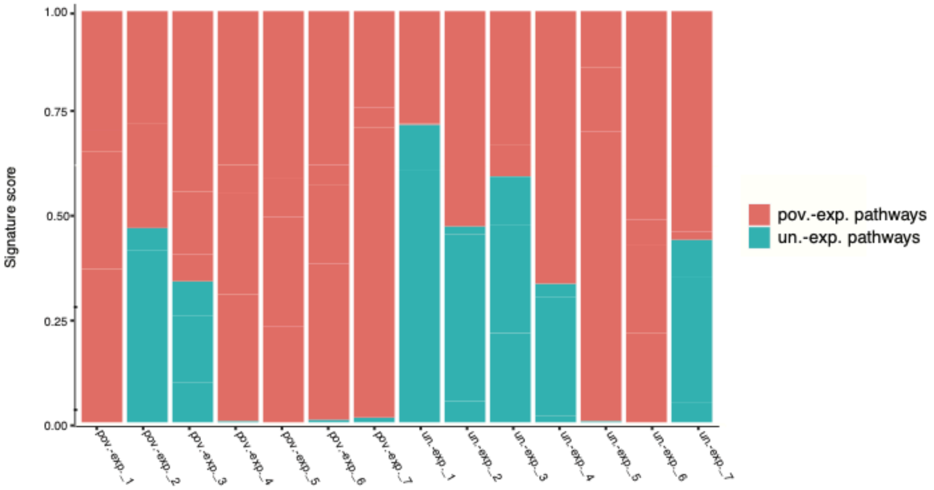


**Suppl Figure S6**

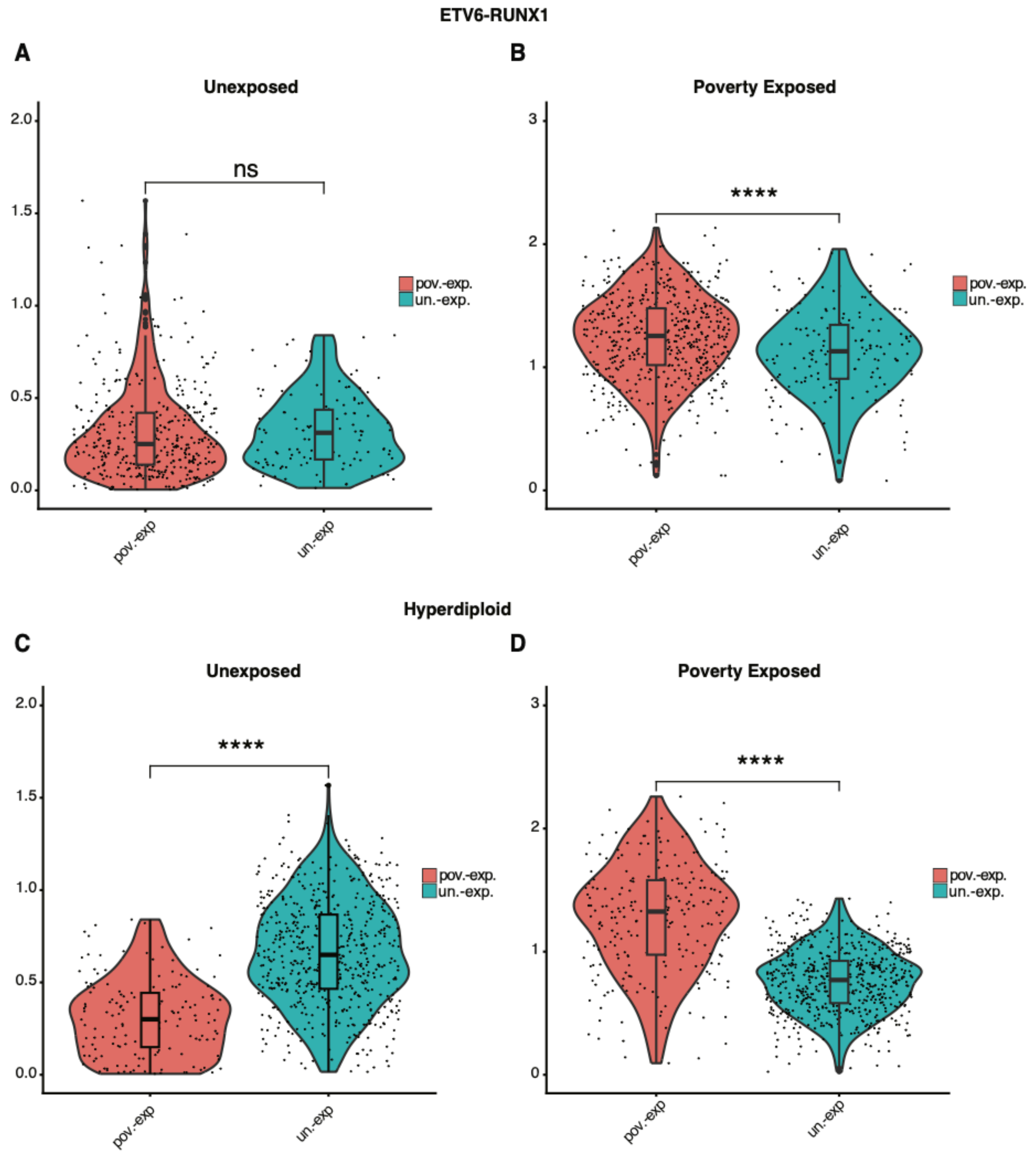


**Suppl Figure S7**

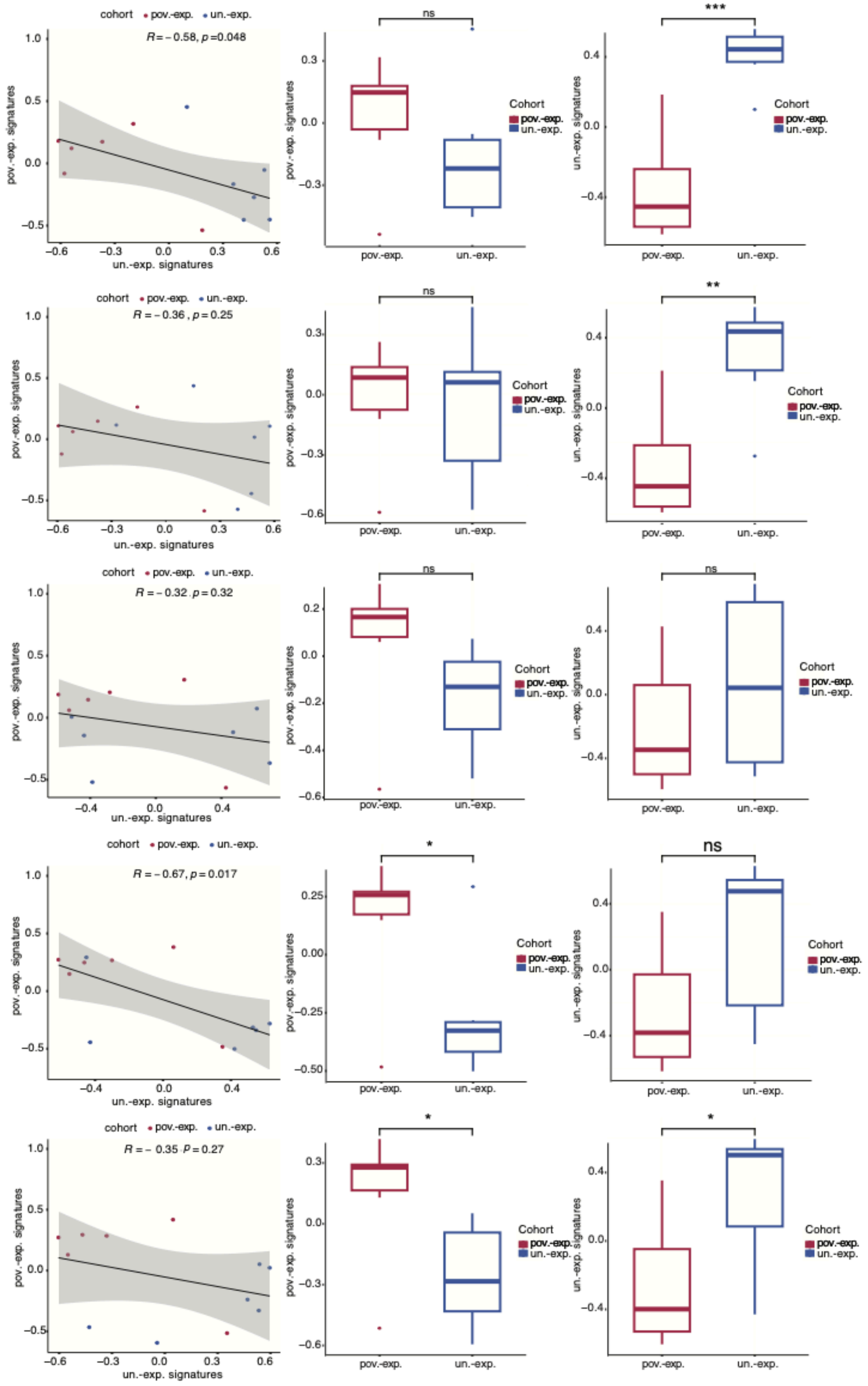


**Suppl Figure S8**

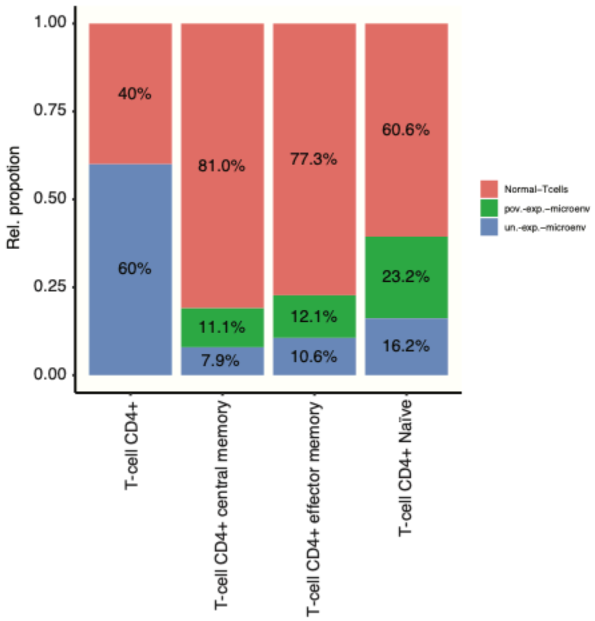


**Suppl Figure S9**

Table_S1_HCA signatures

| | hsc | clp | prob | immature | naiveb |
|---|---|---|---|---|---|
| 1 | AHSP | ADNP | AFM | ÔªøCD22 | BLK |
| 2 | ALAS2 | AIMP1 | ALPL | CYBB | CXCR5 |
| 3 | ATP8B4 | ASCC1 | CCR8 | FAM129C | CD19 |
| 4 | CA1 | ATP6V1G1 | CRX | FCRL1 | MS4A1 |
| 5 | CCDC121 | C11orf57 | DCC | FCRL3 | CD72 |
| 6 | CD34 | C19orf53 | DNTT | FCRL5 | SPIB |
| 7 | CRHBP | CALM1 | GABRA6 | FCRLA | TCL1A |
| 8 | ERG | COX6C | GCK | HDAC9 | FCRL2 |
| 9 | EXD2 | DNTT | GPR3 | HLA-DQA1 | BLK |
| 10 | FAM124B | DSTN | GPR4 | HVCN1 | CXCR5 |
| 11 | FLT3 | FAM76A | GRIK3 | KIAA0226 | CD19 |
| 12 | GSTM5 | GAPDH | GYS2 | NCF1 | MS4A1 |
| 13 | GYPA | GPN3 | TLX2 | NCF1B | CD22 |
| 14 | GYPE | H3F3B | KNG1 | P2RY10 | CD72 |
| 15 | KLHL9 | HIVEP3 | MEN1 | SP100 | CD79B |
| 16 | LAPTM4B | IDH3A | MUSK | TXNIP | CCR6 |
| 17 | LSM2 | IGBP1 | OMD | STAP1 | GPR18 |
| 18 | MMRN1 | IGLL1 | PARK2 | TAGAP | CD180 |
| 19 | MYCT1 | METAP2 | POU3F1 | ÔªøZCCHC2 | MGAT5 |
| 20 | PLS3 | MYL12B | RAG2 | NA | SPIB |
| 21 | SCARF1 | NGLY1 | RPE65 | NA | TCL1A |
| 22 | SLC4A1 | OXA1L | SLC2A2 | NA | MBD4 |
| 23 | TCEAL4 | PLP2 | SLC12A1 | NA | TSPAN13 |
| 24 | XPO7 | PSMA6 | SRY | NA | UTP6 |
| 25 | ZMYM3 | PSMB3 | VPREB1 | NA | FCRL2 |
| 26 | NA | RFC4 | CSRP3 | NA | TREML2 |
| 27 | NA | RPL8 | HIST1H2BM | NA | BLK |
| 28 | NA | SNRPD1 | SOX14 | NA | CXCR5 |
| 29 | NA | TOMM20 | CACNA1G | NA | CD19 |
| 30 | NA | VPREB1 | GREB1 | NA | MS4A1 |
| 31 | NA | WDR33 | CNKSR1 | NA | CD72 |
| 32 | NA | NA | ARPP21 | NA | CD180 |
| 33 | NA | NA | FRS3 | NA | SPIB |
| 34 | NA | NA | LILRB1 | NA | BLK |
| 35 | NA | NA | CLDN14 | NA | CD19 |
| 36 | NA | NA | PLA2G2D | NA | MS4A1 |
| 37 | NA | NA | OR7A5 | NA | CD22 |
| 38 | NA | NA | AKAP8L | NA | CD37 |
| 39 | NA | NA | STRN4 | NA | CD72 |

| | | | | | |
|---|---|---|---|---|---|
| 40 | NA | NA | GNL2 | NA | CD79A |
| 41 | NA | NA | TNFRSF12A | NA | CSNK1G3 |
| 42 | NA | NA | LARP7 | NA | DSP |
| 43 | NA | NA | BTBD7 | NA | FCER2 |
| 44 | NA | NA | SPATA7 | NA | GMFB |
| 45 | NA | NA | HAMP | NA | PNOC |
| 46 | NA | NA | C2orf49 | NA | CD1A |
| 47 | NA | NA | CXorf36 | NA | CD19 |
| 48 | NA | NA | LRRTM4 | NA | MS4A1 |
| 49 | NA | NA | FIP1L1 | NA | CD22 |
| 50 | NA | NA | SLC25A31 | NA | CD37 |
| 51 | NA | NA | SCRT1 | NA | CD72 |
| 52 | NA | NA | IMP4 | NA | CD79A |
| 53 | NA | NA | HSPB6 | NA | CSNK1G3 |
| 54 | NA | NA | ZNF81 | NA | DSP |
| 55 | NA | NA | ZNF674 | NA | FCER2 |
| 56 | NA | NA | ADARB2 | NA | GMFB |
| 57 | NA | NA | ADCY8 | NA | MGAT5 |
| 58 | NA | NA | AGXT | NA | PNOC |
| 59 | NA | NA | ALOX15B | NA | SNX2 |
| 60 | NA | NA | ANXA3 | NA | AP3B1 |
| 61 | NA | NA | APOC3 | NA | MBD4 |
| 62 | NA | NA | ARG1 | NA | STAG3 |
| 63 | NA | NA | ART4 | NA | PRDM4 |
| 64 | NA | NA | BMX | NA | PWP1 |
| 65 | NA | NA | CA1 | NA | RRAS2 |
| 66 | NA | NA | CCKAR | NA | GGA2 |
| 67 | NA | NA | SIGLEC6 | NA | SIPA1L3 |
| 68 | NA | NA | CETP | NA | STAP1 |
| 69 | NA | NA | CLCN1 | NA | P2RY10 |
| 70 | NA | NA | CNTFR | NA | VPREB3 |
| 71 | NA | NA | CRH | NA | DEF8 |
| 72 | NA | NA | CSHL1 | NA | MFN1 |
| 73 | NA | NA | CTSG | NA | FCRL2 |
| 74 | NA | NA | CYP2A7 | NA | BLK |
| 75 | NA | NA | DCC | NA | CAPN3 |
| 76 | NA | NA | DLX4 | NA | CD1A |
| 77 | NA | NA | DNTT | NA | CD19 |
| 78 | NA | NA | DPYS | NA | MS4A1 |
| 79 | NA | NA | DSP | NA | CD22 |
| 80 | NA | NA | EFNA2 | NA | CD37 |

| | | | | | |
|---|---|---|---|---|---|
| 81 | NA | NA | FCAR | NA | CD72 |
| 82 | NA | NA | FCN2 | NA | CD79A |
| 83 | NA | NA | FGF8 | NA | CD79B |
| 84 | NA | NA | MSTN | NA | CSNK1G3 |
| 85 | NA | NA | GDF10 | NA | DSP |
| 86 | NA | NA | GPR3 | NA | FCER2 |
| 87 | NA | NA | GPR4 | NA | GMFB |
| 88 | NA | NA | GRIK3 | NA | HSPA4 |
| 89 | NA | NA | GYS2 | NA | MGAT5 |
| 90 | NA | NA | HCRTR2 | NA | PNOC |
| 91 | NA | NA | ONECUT1 | NA | SNX2 |
| 92 | NA | NA | PRMT1 | NA | GCM1 |
| 93 | NA | NA | HTR1B | NA | AP3B1 |
| 94 | NA | NA | HTR5A | NA | MBD4 |
| 95 | NA | NA | IFNA1 | NA | STAG3 |
| 96 | NA | NA | IGLL1 | NA | PRDM4 |
| 97 | NA | NA | IL12B | NA | PWP1 |
| 98 | NA | NA | KCNJ9 | NA | SP140 |
| 99 | NA | NA | KCNJ13 | NA | RRAS2 |
| 100 | NA | NA | KRT12 | NA | GGA2 |
| 101 | NA | NA | KRT19 | NA | SIPA1L3 |
| 102 | NA | NA | LLGL1 | NA | STAP1 |
| 103 | NA | NA | LTC4S | NA | P2RY10 |
| 104 | NA | NA | MAG | NA | VPREB3 |
| 105 | NA | NA | MKI67 | NA | DEF8 |
| 106 | NA | NA | TRPM1 | NA | MFN1 |
| 107 | NA | NA | MUC6 | NA | FCRL2 |
| 108 | NA | NA | MYH4 | NA | C10orf76 |
| 109 | NA | NA | MYH8 | NA | SMC6 |
| 110 | NA | NA | NEUROG1 | NA | MCM9 |
| 111 | NA | NA | NOTCH4 | NA | EGOT |
| 112 | NA | NA | OMG | NA | CXCR5 |
| 113 | NA | NA | PARK2 | NA | BMP3 |
| 114 | NA | NA | PMP2 | NA | CACNA1F |
| 115 | NA | NA | POU1F1 | NA | CAPN3 |
| 116 | NA | NA | PPEF2 | NA | CD19 |
| 117 | NA | NA | PRB4 | NA | MS4A1 |
| 118 | NA | NA | PSG11 | NA | CD22 |
| 119 | NA | NA | RAD23A | NA | CD72 |
| 120 | NA | NA | RAG2 | NA | COL19A1 |
| 121 | NA | NA | RAPSN | NA | CSNK1G3 |

| | | | | | |
|---|---|---|---|---|---|
| 122 | NA | NA | RBP3 | NA | DAZL |
| 123 | NA | NA | MRPL12 | NA | DSP |
| 124 | NA | NA | RRM2 | NA | FCER2 |
| 125 | NA | NA | CCL17 | NA | GNG3 |
| 126 | NA | NA | TACR3 | NA | GPR18 |
| 127 | NA | NA | TCF3 | NA | MATN1 |
| 128 | NA | NA | TESK1 | NA | MAP3K9 |
| 129 | NA | NA | TGM3 | NA | MMP17 |
| 130 | NA | NA | THPO | NA | MYBPC2 |
| 131 | NA | NA | TNNT2 | NA | PAX5 |
| 132 | NA | NA | UCP3 | NA | PHKG1 |
| 133 | NA | NA | VPREB1 | NA | PYGM |
| 134 | NA | NA | ZNF155 | NA | ZNF154 |
| 135 | NA | NA | CSRP3 | NA | PRDM2 |
| 136 | NA | NA | HIST1H2BL | NA | USP7 |
| 137 | NA | NA | HIST1H2BM | NA | TCL1A |
| 138 | NA | NA | SOX14 | NA | SYN3 |
| 139 | NA | NA | EDF1 | NA | ADAM20 |
| 140 | NA | NA | ADAM21 | NA | USP6 |
| 141 | NA | NA | KCNQ4 | NA | AKAP6 |
| 142 | NA | NA | OTOF | NA | TCL1B |
| 143 | NA | NA | KIF23 | NA | BCL2L10 |
| 144 | NA | NA | CNKSR1 | NA | FRS2 |
| 145 | NA | NA | NMUR1 | NA | PRDM4 |
| 146 | NA | NA | PLK4 | NA | RRAS2 |
| 147 | NA | NA | ARPP21 | NA | GGA2 |
| 148 | NA | NA | FRS3 | NA | SIPA1L3 |
| 149 | NA | NA | LILRB1 | NA | TCL6 |
| 150 | NA | NA | STIP1 | NA | TSPAN13 |
| 151 | NA | NA | LILRB4 | NA | P2RY10 |
| 152 | NA | NA | KERA | NA | MYO3A |
| 153 | NA | NA | ADAMTS8 | NA | SDK2 |
| 154 | NA | NA | CLCA4 | NA | WDR74 |
| 155 | NA | NA | FSTL4 | NA | UBE2O |
| 156 | NA | NA | PMPCA | NA | RBM15 |
| 157 | NA | NA | ATP1B4 | NA | SMC6 |
| 158 | NA | NA | CLDN14 | NA | KHDRBS2 |
| 159 | NA | NA | PADI4 | NA | CXCR5 |
| 160 | NA | NA | TSSK2 | NA | BMP3 |
| 161 | NA | NA | CA14 | NA | CACNA1F |
| 162 | NA | NA | FBXO24 | NA | CAPN3 |

| | | | | | |
|---|---|---|---|---|---|
| 163 | NA | NA | SLC13A4 | NA | CD19 |
| 164 | NA | NA | OR7A5 | NA | MS4A1 |
| 165 | NA | NA | AKAP8L | NA | CD22 |
| 166 | NA | NA | DKKL1 | NA | CD72 |
| 167 | NA | NA | AHDC1 | NA | COL19A1 |
| 168 | NA | NA | BMP10 | NA | CSNK1G3 |
| 169 | NA | NA | VPREB3 | NA | DAZL |
| 170 | NA | NA | ANAPC2 | NA | FCER2 |
| 171 | NA | NA | VSX1 | NA | GNG3 |
| 172 | NA | NA | CALY | NA | MAP3K9 |
| 173 | NA | NA | HP1BP3 | NA | MMP17 |
| 174 | NA | NA | PDE11A | NA | MYBPC2 |
| 175 | NA | NA | CLEC1A | NA | PAX5 |
| 176 | NA | NA | GMIP | NA | PHKG1 |
| 177 | NA | NA | TNFRSF12A | NA | PRDM2 |
| 178 | NA | NA | SPTBN5 | NA | USP7 |
| 179 | NA | NA | ZMYND10 | NA | TCL1A |
| 180 | NA | NA | LARP7 | NA | SYN3 |
| 181 | NA | NA | COQ3 | NA | ADAM20 |
| 182 | NA | NA | CNGB3 | NA | USP6 |
| 183 | NA | NA | ZNF407 | NA | AKAP6 |
| 184 | NA | NA | IQCC | NA | TCL1B |
| 185 | NA | NA | BTBD7 | NA | FRS2 |
| 186 | NA | NA | SPATA7 | NA | PRDM4 |
| 187 | NA | NA | NXF3 | NA | RRAS2 |
| 188 | NA | NA | PAPOLB | NA | GGA2 |
| 189 | NA | NA | RPGRIP1 | NA | SIPA1L3 |
| 190 | NA | NA | SLURP1 | NA | TCL6 |
| 191 | NA | NA | SPC25 | NA | TSPAN13 |
| 192 | NA | NA | HAMP | NA | P2RY10 |
| 193 | NA | NA | MYL7 | NA | WDR74 |
| 194 | NA | NA | KLHL12 | NA | UBE2O |
| 195 | NA | NA | FBRS | NA | SMC6 |
| 196 | NA | NA | RNF25 | NA | KHDRBS2 |
| 197 | NA | NA | TUT1 | NA | RERE |
| 198 | NA | NA | MRPS15 | NA | CXCR5 |
| 199 | NA | NA | KRI1 | NA | BMP3 |
| 200 | NA | NA | NOL12 | NA | CACNA1F |
| 201 | NA | NA | CXorf36 | NA | CAPN3 |
| 202 | NA | NA | KRTAP1-3 | NA | CD1A |
| 203 | NA | NA | PCDH11Y | NA | CD19 |

| | | | | | |
|---|---|---|---|---|---|
| 204 | NA | NA | IMP4 | NA | MS4A1 |
| 205 | NA | NA | HSPB6 | NA | CD22 |
| 206 | NA | NA | PPP4R2 | NA | CD72 |
| 207 | NA | NA | KCNV2 | NA | COL19A1 |
| 208 | NA | NA | PSORS1C2 | NA | CSNK1G3 |
| 209 | NA | NA | R3HCC1 | NA | DAZL |
| 210 | NA | NA | ASPM | NA | DSP |
| 211 | NA | NA | NACA2 | NA | FCER2 |
| 212 | NA | NA | ARG1 | NA | GH1 |
| 213 | NA | NA | CLCN1 | NA | GNG3 |
| 214 | NA | NA | CNTFR | NA | HLA-DOA |
| 215 | NA | NA | CSHL1 | NA | LY9 |
| 216 | NA | NA | DCC | NA | MATN1 |
| 217 | NA | NA | DNTT | NA | CIITA |
| 218 | NA | NA | FCAR | NA | MAP3K9 |
| 219 | NA | NA | FGF8 | NA | MMP17 |
| 220 | NA | NA | GPR3 | NA | MYBPC2 |
| 221 | NA | NA | HTR1B | NA | PAX5 |
| 222 | NA | NA | IGLL1 | NA | PGAM2 |
| 223 | NA | NA | LTC4S | NA | PHKG1 |
| 224 | NA | NA | MEP1B | NA | POU2F1 |
| 225 | NA | NA | TRPM1 | NA | PRKCB |
| 226 | NA | NA | MUC6 | NA | PYGM |
| 227 | NA | NA | PRB4 | NA | RB1 |
| 228 | NA | NA | PSG11 | NA | TRA2B |
| 229 | NA | NA | RAG2 | NA | ZNF154 |
| 230 | NA | NA | MRPL12 | NA | SLC30A4 |
| 231 | NA | NA | SGCA | NA | PRDM2 |
| 232 | NA | NA | TACR3 | NA | USP7 |
| 233 | NA | NA | TCOF1 | NA | CUBN |
| 234 | NA | NA | TESK1 | NA | TCL1A |
| 235 | NA | NA | TGM3 | NA | SYN3 |
| 236 | NA | NA | VPREB1 | NA | CDK13 |
| 237 | NA | NA | ZNF155 | NA | PTCH2 |
| 238 | NA | NA | HIST1H2BL | NA | ADAM20 |
| 239 | NA | NA | SOX14 | NA | USP6 |
| 240 | NA | NA | ADAM21 | NA | AKAP6 |
| 241 | NA | NA | KCNQ4 | NA | TCL1B |
| 242 | NA | NA | ARPP21 | NA | TBC1D5 |
| 243 | NA | NA | FRS3 | NA | BCL2L11 |
| 244 | NA | NA | LILRB1 | NA | STAG3 |

| | | | | | |
|---|---|---|---|---|---|
| 245 | NA | NA | ADAMTS8 | NA | FRS2 |
| 246 | NA | NA | LAMB4 | NA | PRDM4 |
| 247 | NA | NA | CLDN14 | NA | RRAS2 |
| 248 | NA | NA | FBXO24 | NA | GGA2 |
| 249 | NA | NA | AKAP8L | NA | SIPA1L3 |
| 250 | NA | NA | AHDC1 | NA | N4BP3 |
| 251 | NA | NA | BMP10 | NA | TCL6 |
| 252 | NA | NA | VSX1 | NA | TSPAN13 |
| 253 | NA | NA | PCDHA5 | NA | P2RY10 |
| 254 | NA | NA | OTUD7B | NA | SNTG2 |
| 255 | NA | NA | SPC25 | NA | SDK2 |
| 256 | NA | NA | HAMP | NA | WDR74 |
| 257 | NA | NA | MYL7 | NA | UBE2O |
| 258 | NA | NA | DPEP3 | NA | NOC3L |
| 259 | NA | NA | C2orf49 | NA | RBM15 |
| 260 | NA | NA | CXorf36 | NA | FCRL2 |
| 261 | NA | NA | IMP4 | NA | SMC6 |
| 262 | NA | NA | HSPB6 | NA | TRAPPC9 |
| 263 | NA | NA | AZU1 | NA | PIKFYVE |
| 264 | NA | NA | BLK | NA | KHDRBS2 |
| 265 | NA | NA | CD72 | NA | 42802 |
| 266 | NA | NA | CD79B | NA | NA |
| 267 | NA | NA | CENPA | NA | NA |
| 268 | NA | NA | CETP | NA | NA |
| 269 | NA | NA | DNTT | NA | NA |
| 270 | NA | NA | FOXM1 | NA | NA |
| 271 | NA | NA | FLT3 | NA | NA |
| 272 | NA | NA | H2AFX | NA | NA |
| 273 | NA | NA | IGLL1 | NA | NA |
| 274 | NA | NA | KIF11 | NA | NA |
| 275 | NA | NA | LY6H | NA | NA |
| 276 | NA | NA | MKI67 | NA | NA |
| 277 | NA | NA | MYBL2 | NA | NA |
| 278 | NA | NA | PDE6D | NA | NA |
| 279 | NA | NA | POLA1 | NA | NA |
| 280 | NA | NA | PRTN3 | NA | NA |
| 281 | NA | NA | RAG2 | NA | NA |
| 282 | NA | NA | RFC2 | NA | NA |
| 283 | NA | NA | RFC5 | NA | NA |
| 284 | NA | NA | RRM2 | NA | NA |
| 285 | NA | NA | SMARCA4 | NA | NA |

| | | | | | |
|---|---|---|---|---|---|
| 286 | NA | NA | SNRPD1 | NA | NA |
| 287 | NA | NA | SPTA1 | NA | NA |
| 288 | NA | NA | TCF3 | NA | NA |
| 289 | NA | NA | TERT | NA | NA |
| 290 | NA | NA | TOP2B | NA | NA |
| 291 | NA | NA | TSSC1 | NA | NA |
| 292 | NA | NA | VPREB1 | NA | NA |
| 293 | NA | NA | XPNPEP2 | NA | NA |
| 294 | NA | NA | PTTG1 | NA | NA |
| 295 | NA | NA | TCL1B | NA | NA |
| 296 | NA | NA | ESPL1 | NA | NA |
| 297 | NA | NA | KIF14 | NA | NA |
| 298 | NA | NA | P2RY14 | NA | NA |
| 299 | NA | NA | SMC4 | NA | NA |
| 300 | NA | NA | TACC3 | NA | NA |
| 301 | NA | NA | NOP56 | NA | NA |
| 302 | NA | NA | SIVA1 | NA | NA |
| 303 | NA | NA | ARPP21 | NA | NA |
| 304 | NA | NA | HNRNPA0 | NA | NA |
| 305 | NA | NA | UBE2C | NA | NA |
| 306 | NA | NA | KIF4A | NA | NA |
| 307 | NA | NA | OR7A5 | NA | NA |
| 308 | NA | NA | VPREB3 | NA | NA |
| 309 | NA | NA | TRA2A | NA | NA |
| 310 | NA | NA | SAC3D1 | NA | NA |
| 311 | NA | NA | MRTO4 | NA | NA |
| 312 | NA | NA | NUSAP1 | NA | NA |
| 313 | NA | NA | AHSP | NA | NA |
| 314 | NA | NA | GTSE1 | NA | NA |
| 315 | NA | NA | CEP55 | NA | NA |
| 316 | NA | NA | QRSL1 | NA | NA |
| 317 | NA | NA | SPC25 | NA | NA |
| 318 | NA | NA | LSM2 | NA | NA |
| 319 | NA | NA | CCDC81 | NA | NA |
| 320 | NA | NA | SHCBP1 | NA | NA |
| 321 | NA | NA | C16orf59 | NA | NA |
| 322 | NA | NA | HPS4 | NA | NA |
| 323 | NA | NA | BLK | NA | NA |
| 324 | NA | NA | CD72 | NA | NA |
| 325 | NA | NA | CD79B | NA | NA |
| 326 | NA | NA | DNTT | NA | NA |

| | | | | | |
|---|---|---|---|---|---|
| 327 | NA | NA | FLT3 | NA | NA |
| 328 | NA | NA | IGLL1 | NA | NA |
| 329 | NA | NA | PRTN3 | NA | NA |
| 330 | NA | NA | RAG2 | NA | NA |
| 331 | NA | NA | VPREB1 | NA | NA |
| 332 | NA | NA | ARPP21 | NA | NA |
| 333 | NA | NA | VPREB3 | NA | NA |
| 334 | NA | NA | QRSL1 | NA | NA |
| 335 | NA | NA | CCDC81 | NA | NA |
| 336 | NA | NA | BLK | NA | NA |
| 337 | NA | NA | CD72 | NA | NA |
| 338 | NA | NA | CD79B | NA | NA |
| 339 | NA | NA | DNTT | NA | NA |
| 340 | NA | NA | FLT3 | NA | NA |
| 341 | NA | NA | IGLL1 | NA | NA |
| 342 | NA | NA | LY6H | NA | NA |
| 343 | NA | NA | MYBL2 | NA | NA |
| 344 | NA | NA | PRTN3 | NA | NA |
| 345 | NA | NA | RAG2 | NA | NA |
| 346 | NA | NA | VPREB1 | NA | NA |
| 347 | NA | NA | ARPP21 | NA | NA |
| 348 | NA | NA | VPREB3 | NA | NA |
| 349 | NA | NA | QRSL1 | NA | NA |
| 350 | NA | NA | CCDC81 | NA | NA |

Table_S2_PovertyEx_vs_UnExp_upr

| | Top | p.value | FDR | logFC |
|---|---|---|---|---|
| RAG1 | 1 | 2.25E-88 | 6.67E-84 | 3.554289072 |
| PCDH9 | 10 | 2.41E-61 | 7.14E-58 | 3.158456629 |
| TERF2 | 5 | 1.26E-71 | 7.44E-68 | 3.016238331 |
| IGF2BP1 | 2 | 1.11E-77 | 1.64E-73 | 2.664836213 |
| TCL1A | 15 | 3.25E-52 | 6.42E-49 | 2.480287049 |
| NRN1 | 11 | 7.41E-61 | 1.99E-57 | 2.394049148 |
| TP53INP1 | 22 | 2.69E-44 | 3.61E-41 | 2.362609764 |
| GNG11 | 12 | 2.28E-58 | 5.63E-55 | 2.309571438 |
| STAG3 | 21 | 1.10E-44 | 1.55E-41 | 2.283671314 |
| MME | 29 | 3.51E-42 | 3.58E-39 | 2.248374891 |
| FYB1 | 33 | 1.51E-40 | 1.35E-37 | 2.171166417 |
| HPS4 | 58 | 8.71E-34 | 4.45E-31 | 2.081803009 |
| PTPRK | 14 | 1.72E-54 | 3.64E-51 | 2.053654891 |
| MCTP2 | 17 | 5.70E-51 | 9.93E-48 | 1.938042477 |
| SEMA6A | 27 | 6.62E-43 | 7.25E-40 | 1.914447715 |
| TNS1 | 18 | 1.58E-49 | 2.60E-46 | 1.911330794 |
| SDC2 | 19 | 2.36E-47 | 3.67E-44 | 1.906945926 |
| ABHD3 | 64 | 5.32E-33 | 2.46E-30 | 1.90236241 |
| SCN3A | 31 | 2.33E-41 | 2.22E-38 | 1.862227646 |
| ERGIC1 | 74 | 5.42E-31 | 2.17E-28 | 1.807464459 |
| PLEKHG4B | 23 | 8.51E-44 | 1.09E-40 | 1.797918858 |
| INSR | 76 | 1.26E-30 | 4.91E-28 | 1.78219652 |
| SOCS2 | 132 | 3.03E-23 | 6.80E-21 | 1.763923656 |
| CD19 | 112 | 6.28E-25 | 1.66E-22 | 1.7449401 |
| SSBP2 | 104 | 7.68E-26 | 2.19E-23 | 1.723628636 |
| GAB1 | 110 | 2.96E-25 | 7.97E-23 | 1.713559795 |
| ARHGAP24 | 35 | 1.23E-39 | 1.04E-36 | 1.712228876 |
| CACNB2 | 50 | 8.36E-36 | 4.95E-33 | 1.681703199 |
| POU2AF1 | 47 | 1.23E-36 | 7.73E-34 | 1.626945799 |
| MYO10 | 30 | 1.41E-41 | 1.39E-38 | 1.611554137 |
| PHYH | 91 | 3.67E-28 | 1.19E-25 | 1.598110747 |
| IGLL1 | 242 | 2.83E-17 | 3.47E-15 | 1.590059179 |
| PCLO | 43 | 3.72E-37 | 2.56E-34 | 1.588537521 |
| FHIT | 114 | 8.45E-25 | 2.19E-22 | 1.565663649 |
| LINC01013 | 190 | 1.60E-19 | 2.50E-17 | 1.549685986 |
| EDEM1 | 138 | 6.67E-23 | 1.43E-20 | 1.54393183 |
| TMED6 | 44 | 7.41E-37 | 4.93E-34 | 1.538334838 |
| CMTM7 | 187 | 8.15E-20 | 1.29E-17 | 1.532507156 |
| IDI1 | 232 | 1.19E-17 | 1.52E-15 | 1.522251487 |
| CRELD2 | 67 | 9.87E-33 | 4.36E-30 | 1.480353732 |
| BEST3 | 49 | 4.59E-36 | 2.77E-33 | 1.477857809 |

| | | | | |
|---|---|---|---|---|
| MYADM | 85 | 1.13E-29 | 3.93E-27 | 1.477003699 |
| SMAD1 | 156 | 6.80E-22 | 1.29E-19 | 1.475411078 |
| TNFRSF21 | 87 | 7.72E-29 | 2.63E-26 | 1.471144129 |
| SPTBN1 | 169 | 5.33E-21 | 9.34E-19 | 1.453743143 |
| DPEP1 | 274 | 4.27E-16 | 4.62E-14 | 1.435085685 |
| SFXN1 | 149 | 4.28E-22 | 8.50E-20 | 1.422602757 |
| WASF2 | 78 | 1.93E-30 | 7.31E-28 | 1.41583946 |
| BLACE | 181 | 2.80E-20 | 4.58E-18 | 1.40251741 |
| MIB1 | 143 | 1.32E-22 | 2.73E-20 | 1.402427799 |
| PAX5 | 142 | 1.26E-22 | 2.63E-20 | 1.402400876 |
| NEIL1 | 317 | 1.08E-14 | 1.01E-12 | 1.393439873 |
| STK32B | 158 | 9.01E-22 | 1.69E-19 | 1.390087228 |
| CALN1 | 41 | 1.98E-37 | 1.43E-34 | 1.382201395 |
| CIITA | 105 | 1.58E-25 | 4.45E-23 | 1.380669674 |
| FBXW7 | 307 | 5.81E-15 | 5.60E-13 | 1.373621606 |
| SLC35E3 | 285 | 1.20E-15 | 1.25E-13 | 1.373057929 |
| ISG20 | 217 | 3.51E-18 | 4.79E-16 | 1.364679159 |
| LAIR1 | 243 | 3.06E-17 | 3.73E-15 | 1.363965475 |
| MDM2 | 183 | 3.04E-20 | 4.92E-18 | 1.341750835 |
| H1-0 | 197 | 3.34E-19 | 5.02E-17 | 1.340331425 |
| PEAK1 | 126 | 9.70E-24 | 2.28E-21 | 1.336445147 |
| FARP1 | 62 | 3.30E-33 | 1.58E-30 | 1.330934334 |
| BIRC7 | 69 | 5.33E-32 | 2.29E-29 | 1.282970311 |
| VPREB3 | 289 | 1.60E-15 | 1.63E-13 | 1.280094565 |
| SHROOM3 | 97 | 3.07E-27 | 9.38E-25 | 1.274728845 |
| UXS1 | 293 | 2.26E-15 | 2.28E-13 | 1.27395114 |
| SP4 | 260 | 1.16E-16 | 1.31E-14 | 1.265807639 |
| CBFA2T3 | 144 | 1.43E-22 | 2.93E-20 | 1.259948293 |
| TFPI | 173 | 9.02E-21 | 1.54E-18 | 1.255151353 |
| MDK | 204 | 1.09E-18 | 1.59E-16 | 1.25441367 |
| ITPR1 | 286 | 1.42E-15 | 1.47E-13 | 1.237783086 |
| BACH2 | 218 | 4.27E-18 | 5.78E-16 | 1.237361629 |
| KNTC1 | 265 | 1.89E-16 | 2.11E-14 | 1.232069484 |
| FCHSD2 | 256 | 9.06E-17 | 1.05E-14 | 1.210484086 |
| GNG7 | 226 | 6.61E-18 | 8.66E-16 | 1.199873364 |
| STMN1 | 570 | 5.33E-10 | 2.77E-08 | 1.197319131 |
| HLA-DOA | 386 | 4.64E-13 | 3.55E-11 | 1.186907905 |
| TSPO | 309 | 6.66E-15 | 6.38E-13 | 1.167579085 |
| HSPB1 | 180 | 1.73E-20 | 2.84E-18 | 1.163102802 |
| SH2D4B | 368 | 2.46E-13 | 1.97E-11 | 1.157828924 |
| DRAM1 | 207 | 1.23E-18 | 1.76E-16 | 1.157692364 |
| KCNN1 | 54 | 2.07E-34 | 1.14E-31 | 1.142608265 |
| PDE4D | 108 | 2.07E-25 | 5.66E-23 | 1.14151581 |

| | | | | |
|---|---|---|---|---|
| FADS3 | 224 | 6.28E-18 | 8.29E-16 | 1.12925717 |
| HAP1 | 68 | 5.20E-32 | 2.26E-29 | 1.12595743 |
| EGFL7 | 330 | 2.73E-14 | 2.45E-12 | 1.12157289 |
| SYK | 417 | 3.94E-12 | 2.79E-10 | 1.105162725 |
| DOK3 | 393 | 7.02E-13 | 5.29E-11 | 1.103023695 |
| ANKRD30B | 93 | 8.45E-28 | 2.69E-25 | 1.099309604 |
| MTCO1P12 | 266 | 2.01E-16 | 2.23E-14 | 1.095409006 |
| OVCH2 | 436 | 9.82E-12 | 6.65E-10 | 1.088683532 |
| TUSC3 | 82 | 3.40E-30 | 1.23E-27 | 1.087206132 |
| LINC01416 | 77 | 1.49E-30 | 5.74E-28 | 1.080805865 |
| FOXO1 | 452 | 1.78E-11 | 1.16E-09 | 1.076869121 |
| FCMR | 295 | 2.49E-15 | 2.50E-13 | 1.068192914 |
| NKAIN4 | 84 | 5.31E-30 | 1.87E-27 | 1.060596122 |
| AKAP12 | 163 | 1.99E-21 | 3.60E-19 | 1.056025629 |
| HEMGN | 396 | 1.06E-12 | 7.90E-11 | 1.051977499 |
| CDK6 | 482 | 4.36E-11 | 2.67E-09 | 1.05032313 |
| ATP1B3 | 668 | 5.97E-09 | 2.64E-07 | 1.048481604 |
| COL24A1 | 117 | 1.54E-24 | 3.89E-22 | 1.042756073 |
| FAM107B | 360 | 1.34E-13 | 1.10E-11 | 1.036668119 |
| FKBP1A | 495 | 6.84E-11 | 4.09E-09 | 1.034327569 |
| NASP | 656 | 4.53E-09 | 2.04E-07 | 1.026700909 |
| CD27 | 102 | 4.36E-26 | 1.26E-23 | 1.023768611 |
| NR3C2 | 107 | 2.00E-25 | 5.53E-23 | 1.022664025 |
| DSC3 | 103 | 4.57E-26 | 1.31E-23 | 1.010693464 |
| TCL1B | 46 | 1.04E-36 | 6.70E-34 | 1.00851187 |
| HLA-DRA | 202 | 7.64E-19 | 1.12E-16 | 1.003046135 |
| PRDX1 | 494 | 6.75E-11 | 4.05E-09 | 1.000735078 |
| INO80 | 522 | 1.70E-10 | 9.65E-09 | 0.999650257 |
| ARHGEF4 | 120 | 3.74E-24 | 9.22E-22 | 0.996857078 |
| NREP | 414 | 2.89E-12 | 2.06E-10 | 0.986852803 |
| RUBCNL | 318 | 1.21E-14 | 1.13E-12 | 0.98286308 |
| ADGRF1 | 195 | 3.23E-19 | 4.88E-17 | 0.982596508 |
| LOC1025462[illegible] | 116 | 1.12E-24 | 2.86E-22 | 0.979523568 |
| HLA-DPA1 | 468 | 2.77E-11 | 1.75E-09 | 0.972753662 |
| TCF3 | 407 | 2.21E-12 | 1.61E-10 | 0.972704194 |
| LINC00676 | 106 | 1.74E-25 | 4.86E-23 | 0.970058212 |
| UNC79 | 185 | 3.89E-20 | 6.22E-18 | 0.969717928 |
| MSR1 | 248 | 5.58E-17 | 6.66E-15 | 0.968717708 |
| LTB | 389 | 5.65E-13 | 4.29E-11 | 0.967287378 |
| ATP6V1G1 | 599 | 1.35E-09 | 6.66E-08 | 0.967179594 |
| LOC283299 | 496 | 7.17E-11 | 4.28E-09 | 0.962224433 |
| LINC00487 | 131 | 2.67E-23 | 6.04E-21 | 0.961733834 |
| KIF26B | 115 | 1.11E-24 | 2.86E-22 | 0.958568248 |

| | | | | |
|---|---|---|---|---|
| PTPN7 | 247 | 4.81E-17 | 5.76E-15 | 0.958551453 |
| FOXP1 | 563 | 4.70E-10 | 2.47E-08 | 0.956491892 |
| PIK3C3 | 608 | 1.54E-09 | 7.48E-08 | 0.956312435 |
| AK7 | 130 | 2.35E-23 | 5.36E-21 | 0.952244504 |
| CDCA7 | 598 | 1.26E-09 | 6.23E-08 | 0.948877933 |
| LINC00670 | 72 | 1.51E-31 | 6.22E-29 | 0.945928005 |
| CRMP1 | 196 | 3.23E-19 | 4.88E-17 | 0.944952775 |
| INO80C | 399 | 1.28E-12 | 9.48E-11 | 0.944139632 |
| MLXIP | 577 | 7.40E-10 | 3.79E-08 | 0.931416775 |
| UGP2 | 728 | 1.92E-08 | 7.80E-07 | 0.929960033 |
| TSPAN7 | 714 | 1.44E-08 | 5.98E-07 | 0.929559036 |
| ARPP21 | 849 | 1.15E-07 | 4.01E-06 | 0.9289535 |
| NOTCH1 | 322 | 1.84E-14 | 1.69E-12 | 0.928687936 |
| TCFL5 | 394 | 8.25E-13 | 6.20E-11 | 0.924842646 |
| ARHGEF12 | 193 | 2.87E-19 | 4.40E-17 | 0.922968551 |
| HLA-DQB1 | 475 | 3.23E-11 | 2.01E-09 | 0.920725084 |
| CALD1 | 150 | 4.70E-22 | 9.26E-20 | 0.918575977 |
| GNAS | 535 | 2.44E-10 | 1.35E-08 | 0.916025075 |
| SPTAN1 | 730 | 1.96E-08 | 7.96E-07 | 0.913942045 |
| ANKRD10 | 442 | 1.18E-11 | 7.93E-10 | 0.912655417 |
| NDUFAF6 | 444 | 1.31E-11 | 8.74E-10 | 0.909631048 |
| IKZF1 | 678 | 7.00E-09 | 3.05E-07 | 0.90864318 |
| HLA-DPB1 | 401 | 1.30E-12 | 9.57E-11 | 0.906179447 |
| GIMAP2 | 538 | 2.70E-10 | 1.49E-08 | 0.899099433 |
| ZNF608 | 361 | 1.45E-13 | 1.19E-11 | 0.897683871 |
| BMP3 | 512 | 1.24E-10 | 7.16E-09 | 0.897190977 |
| TXLNGY | 665 | 5.37E-09 | 2.39E-07 | 0.897148308 |
| TENM4 | 225 | 6.57E-18 | 8.63E-16 | 0.895873822 |
| ZNF827 | 305 | 5.17E-15 | 5.02E-13 | 0.890719947 |
| TRAF5 | 466 | 2.60E-11 | 1.65E-09 | 0.8844134 |
| CMTM8 | 315 | 9.46E-15 | 8.89E-13 | 0.883418018 |
| LINC02273 | 461 | 2.18E-11 | 1.40E-09 | 0.877502072 |
| MZB1 | 534 | 2.42E-10 | 1.34E-08 | 0.876314333 |
| CHD7 | 778 | 3.96E-08 | 1.51E-06 | 0.873995014 |
| JUN | 529 | 2.09E-10 | 1.17E-08 | 0.873617781 |
| PRKCB | 799 | 5.23E-08 | 1.94E-06 | 0.873292045 |
| NETO1 | 188 | 1.03E-19 | 1.62E-17 | 0.873196703 |
| TMEM243 | 707 | 1.31E-08 | 5.50E-07 | 0.867599081 |
| CIAO3 | 701 | 1.18E-08 | 5.00E-07 | 0.864863943 |
| SDHA | 961 | 3.92E-07 | 1.21E-05 | 0.864354634 |
| CEMIP2 | 595 | 1.06E-09 | 5.27E-08 | 0.862907605 |
| PSME2 | 886 | 1.71E-07 | 5.70E-06 | 0.8588975 |
| TPTEP1 | 241 | 2.62E-17 | 3.22E-15 | 0.853974448 |

| | | | | |
|---|---|---|---|---|
| CDH4 | 166 | 3.22E-21 | 5.74E-19 | 0.841536414 |
| SOCS2-AS1 | 698 | 1.12E-08 | 4.76E-07 | 0.841404451 |
| LIG4 | 583 | 8.14E-10 | 4.13E-08 | 0.838533562 |
| EBF1 | 645 | 3.72E-09 | 1.71E-07 | 0.8378749 |
| UCP2 | 1099 | 1.66E-06 | 4.46E-05 | 0.836733706 |
| POLE | 979 | 4.62E-07 | 1.40E-05 | 0.832101795 |
| UBA7 | 1012 | 6.44E-07 | 1.88E-05 | 0.830457022 |
| SORBS2 | 222 | 5.57E-18 | 7.43E-16 | 0.830036179 |
| ENOSF1 | 381 | 3.94E-13 | 3.06E-11 | 0.829517148 |
| NPY | 1004 | 5.98E-07 | 1.76E-05 | 0.827271841 |
| SLC12A2 | 568 | 5.24E-10 | 2.73E-08 | 0.827171039 |
| RBMS3 | 170 | 6.92E-21 | 1.20E-18 | 0.826107324 |
| CHCHD7 | 885 | 1.69E-07 | 5.65E-06 | 0.823940024 |
| CBX5 | 1093 | 1.54E-06 | 4.17E-05 | 0.823434002 |
| HLA-DQB2 | 346 | 6.17E-14 | 5.27E-12 | 0.822567789 |
| MYO18B | 451 | 1.77E-11 | 1.16E-09 | 0.819664105 |
| SCMH1 | 673 | 6.48E-09 | 2.85E-07 | 0.817149346 |
| CPXM1 | 460 | 2.11E-11 | 1.36E-09 | 0.812478869 |
| BMPR1B | 192 | 2.21E-19 | 3.41E-17 | 0.807796442 |
| RCSD1 | 752 | 2.78E-08 | 1.10E-06 | 0.805925522 |
| GOLGA8B | 240 | 2.43E-17 | 2.99E-15 | 0.80068343 |
| LPAR6 | 1176 | 3.81E-06 | 9.58E-05 | 0.80040069 |
| MTCO1P40 | 167 | 4.32E-21 | 7.66E-19 | 0.795850243 |
| COMMD4 | 510 | 1.09E-10 | 6.30E-09 | 0.793557722 |
| CHD9 | 1074 | 1.29E-06 | 3.57E-05 | 0.791407426 |
| UBE2I | 711 | 1.40E-08 | 5.81E-07 | 0.788625 |
| HBS1L | 1185 | 4.08E-06 | 0.0001018 | 0.788455353 |
| IGHV5-78 | 523 | 1.73E-10 | 9.81E-09 | 0.785219067 |
| FUS | 817 | 6.44E-08 | 2.33E-06 | 0.784058438 |
| LILRA2 | 1017 | 6.69E-07 | 1.95E-05 | 0.78162401 |
| ARID5B | 1410 | 2.13E-05 | 0.00044676 | 0.780981484 |
| CLIC5 | 134 | 5.28E-23 | 1.17E-20 | 0.780592179 |
| CD52 | 735 | 2.32E-08 | 9.33E-07 | 0.779289592 |
| PARP1 | 1285 | 9.86E-06 | 0.00022712 | 0.77620083 |
| ARHGDIB | 749 | 2.76E-08 | 1.09E-06 | 0.774712531 |
| FLI1 | 734 | 2.31E-08 | 9.30E-07 | 0.774616901 |
| TAPT1-AS1 | 717 | 1.60E-08 | 6.60E-07 | 0.762561955 |
| SLAMF6 | 587 | 9.37E-10 | 4.72E-08 | 0.761989573 |
| PSD3 | 1198 | 4.56E-06 | 0.00011256 | 0.761385464 |
| CUX1 | 989 | 5.16E-07 | 1.54E-05 | 0.755279172 |
| NUCB2 | 1196 | 4.51E-06 | 0.00011157 | 0.748960817 |
| CYB5R2 | 623 | 2.29E-09 | 1.09E-07 | 0.742780145 |
| PLEKHA5 | 474 | 3.03E-11 | 1.89E-09 | 0.741610187 |

| | | | | |
|---|---|---|---|---|
| PTGDR | 278 | 6.20E-16 | 6.60E-14 | 0.741180255 |
| GNPTAB | 1008 | 6.29E-07 | 1.85E-05 | 0.740872877 |
| TOP2B | 1223 | 5.76E-06 | 0.00013948 | 0.738919938 |
| HMGN3 | 1133 | 2.41E-06 | 6.29E-05 | 0.732576667 |
| INKA1 | 909 | 2.03E-07 | 6.61E-06 | 0.730602876 |
| RERGL | 209 | 1.63E-18 | 2.31E-16 | 0.729370963 |
| LITAF | 560 | 4.29E-10 | 2.27E-08 | 0.725449884 |
| FAM214A | 889 | 1.74E-07 | 5.79E-06 | 0.719758676 |
| MIR5195 | 434 | 9.56E-12 | 6.52E-10 | 0.716880282 |
| GBP4 | 1274 | 9.32E-06 | 0.00021656 | 0.715966443 |
| SCARB1 | 819 | 6.78E-08 | 2.45E-06 | 0.7157405 |
| TAFA1 | 194 | 3.23E-19 | 4.88E-17 | 0.715739727 |
| WBP1L | 948 | 3.29E-07 | 1.03E-05 | 0.713164545 |
| BID | 852 | 1.19E-07 | 4.12E-06 | 0.712454096 |
| BCL7A | 878 | 1.58E-07 | 5.32E-06 | 0.709253983 |
| OAZ1 | 860 | 1.25E-07 | 4.31E-06 | 0.705646211 |
| MYOCD | 182 | 2.83E-20 | 4.60E-18 | 0.701457966 |
| CALM3 | 721 | 1.80E-08 | 7.40E-07 | 0.69684031 |
| LINC01237 | 415 | 3.47E-12 | 2.47E-10 | 0.695429472 |
| PRX | 349 | 7.78E-14 | 6.60E-12 | 0.695334425 |
| H2AC6 | 1512 | 3.82E-05 | 0.00074701 | 0.692193148 |
| RAG2 | 501 | 7.95E-11 | 4.69E-09 | 0.691760768 |
| HLA-DOB | 258 | 9.55E-17 | 1.10E-14 | 0.690644798 |
| ARGLU1 | 787 | 4.43E-08 | 1.67E-06 | 0.69039446 |
| SNRNP27 | 676 | 6.90E-09 | 3.02E-07 | 0.689852586 |
| TRIM24 | 655 | 4.53E-09 | 2.04E-07 | 0.685740548 |
| TRIM27 | 1471 | 3.13E-05 | 0.00062902 | 0.682692173 |
| HLA-DMB | 1548 | 4.52E-05 | 0.00086348 | 0.682598865 |
| RSRC2 | 1257 | 7.73E-06 | 0.00018204 | 0.682025119 |
| PLCG1 | 339 | 3.91E-14 | 3.41E-12 | 0.682003663 |
| HHIP | 287 | 1.53E-15 | 1.58E-13 | 0.68057874 |
| GIMAP4 | 1360 | 1.64E-05 | 0.00035675 | 0.680484435 |
| ZNF107 | 1091 | 1.54E-06 | 4.17E-05 | 0.676022435 |
| TRIO | 755 | 2.87E-08 | 1.12E-06 | 0.672513232 |
| B2M | 382 | 4.11E-13 | 3.19E-11 | 0.671652045 |
| CBX2 | 362 | 1.49E-13 | 1.22E-11 | 0.671028933 |
| CCDC93 | 811 | 6.13E-08 | 2.23E-06 | 0.670796349 |
| MCM2 | 1363 | 1.67E-05 | 0.0003616 | 0.670258824 |
| PIK3IP1 | 1297 | 1.11E-05 | 0.00025217 | 0.669902386 |
| DLGAP2 | 343 | 5.31E-14 | 4.58E-12 | 0.660540054 |
| TSPYL5 | 297 | 3.41E-15 | 3.39E-13 | 0.660114342 |
| NUMA1 | 1218 | 5.41E-06 | 0.00013134 | 0.657840066 |
| LAPTM5 | 739 | 2.47E-08 | 9.88E-07 | 0.657777272 |

| | | | | |
|---|---|---|---|---|
| RASAL2 | 491 | 5.85E-11 | 3.52E-09 | 0.657231791 |
| VGLL4 | 702 | 1.21E-08 | 5.09E-07 | 0.655837756 |
| CPNE2 | 971 | 4.48E-07 | 1.37E-05 | 0.655483732 |
| PEG3 | 377 | 3.65E-13 | 2.87E-11 | 0.655105343 |
| TRIM38 | 1624 | 6.64E-05 | 0.0012098 | 0.65488439 |
| PCCA | 614 | 1.69E-09 | 8.17E-08 | 0.654710311 |
| TMSB4X | 569 | 5.28E-10 | 2.74E-08 | 0.652976335 |
| NCF4 | 1319 | 1.29E-05 | 0.00028951 | 0.651479118 |
| TMPO | 1649 | 7.40E-05 | 0.00132713 | 0.648488007 |
| ZCCHC7 | 1392 | 1.98E-05 | 0.00042101 | 0.647921996 |
| HNRNPA2B1 | 986 | 5.04E-07 | 1.51E-05 | 0.645944552 |
| THRAP3 | 1343 | 1.52E-05 | 0.00033523 | 0.644601717 |
| MYLK | 754 | 2.82E-08 | 1.11E-06 | 0.644353732 |
| STK39 | 519 | 1.61E-10 | 9.17E-09 | 0.64385282 |
| TIA1 | 1634 | 6.99E-05 | 0.00126673 | 0.643591864 |
| CD74 | 301 | 3.96E-15 | 3.89E-13 | 0.642742826 |
| SLFN13 | 427 | 7.21E-12 | 5.00E-10 | 0.639500626 |
| ACTB | 790 | 4.55E-08 | 1.71E-06 | 0.639077278 |
| ZMYND8 | 1421 | 2.31E-05 | 0.00048151 | 0.638264229 |
| NEK6 | 896 | 1.83E-07 | 6.05E-06 | 0.637374293 |
| GLG1 | 1504 | 3.67E-05 | 0.00072213 | 0.634032139 |
| H2BC12 | 1752 | 0.00011779 | 0.00198947 | 0.632795241 |
| VPS13C | 1516 | 3.86E-05 | 0.00075313 | 0.63262097 |
| NBPF1 | 914 | 2.15E-07 | 6.95E-06 | 0.628428599 |
| H3C4 | 606 | 1.53E-09 | 7.47E-08 | 0.626223262 |
| NIP7 | 1304 | 1.15E-05 | 0.0002614 | 0.625832416 |
| DPF3 | 263 | 1.65E-16 | 1.86E-14 | 0.625679093 |
| UBE2E3 | 499 | 7.58E-11 | 4.50E-09 | 0.624820709 |
| KHDRBS3 | 574 | 5.93E-10 | 3.05E-08 | 0.624737368 |
| RIMKLB | 1662 | 7.92E-05 | 0.00141065 | 0.623462318 |
| TMEM263 | 1932 | 0.00025254 | 0.00386794 | 0.622010159 |
| YEATS2 | 1333 | 1.40E-05 | 0.00031046 | 0.621356556 |
| TKT | 2097 | 0.00048623 | 0.00686124 | 0.620548023 |
| ALDH5A1 | 1317 | 1.28E-05 | 0.00028679 | 0.620479719 |
| HLA-DMA | 1373 | 1.74E-05 | 0.00037548 | 0.620447749 |
| UTY | 1086 | 1.44E-06 | 3.94E-05 | 0.619991067 |
| KLHL6 | 1528 | 4.08E-05 | 0.00078921 | 0.619509831 |
| PSMA5 | 1648 | 7.39E-05 | 0.00132713 | 0.61947284 |
| HSPA4 | 1608 | 6.11E-05 | 0.00112359 | 0.619289844 |
| SRP9 | 1738 | 0.00010851 | 0.00184745 | 0.619063921 |
| PID1 | 259 | 1.13E-16 | 1.29E-14 | 0.618975484 |
| MPP6 | 357 | 9.92E-14 | 8.22E-12 | 0.618704344 |
| ADGRA3 | 913 | 2.15E-07 | 6.95E-06 | 0.61818174 |

| | | | | |
|---|---|---|---|---|
| PRDM2 | 1489 | 3.35E-05 | 0.00066598 | 0.61799659 |
| RFTN1 | 1329 | 1.37E-05 | 0.00030444 | 0.617043903 |
| HGSNAT | 487 | 5.05E-11 | 3.07E-09 | 0.611446679 |
| DAGLB | 1522 | 3.93E-05 | 0.00076487 | 0.610518725 |
| MTF2 | 2060 | 0.00041664 | 0.00598489 | 0.609250989 |
| H2BC5 | 1208 | 4.99E-06 | 0.00012214 | 0.606776416 |
| ODF2L | 704 | 1.24E-08 | 5.23E-07 | 0.60584183 |
| MYB | 1913 | 0.00022533 | 0.00348543 | 0.603396486 |
| LDLRAD4 | 1477 | 3.19E-05 | 0.00063962 | 0.602688899 |
| ADARB2-AS1 | 335 | 3.61E-14 | 3.19E-12 | 0.602013947 |
| KIF16B | 939 | 2.79E-07 | 8.79E-06 | 0.597136477 |
| TLE1 | 1181 | 3.88E-06 | 9.72E-05 | 0.595657366 |
| PCNA | 2200 | 0.00067673 | 0.00910234 | 0.595234961 |
| SMARCA4 | 1701 | 9.43E-05 | 0.00164103 | 0.594632028 |
| VAV3 | 1222 | 5.70E-06 | 0.00013812 | 0.592147374 |
| MAML2 | 789 | 4.55E-08 | 1.70E-06 | 0.590139502 |
| ABCG2 | 363 | 1.57E-13 | 1.28E-11 | 0.588818605 |
| SYVN1 | 1059 | 1.14E-06 | 3.19E-05 | 0.588562767 |
| SOX11 | 298 | 3.42E-15 | 3.39E-13 | 0.588527775 |
| MBNL3 | 1398 | 2.01E-05 | 0.00042604 | 0.588397444 |
| GOLGA8A | 511 | 1.24E-10 | 7.16E-09 | 0.588318438 |
| VAV1 | 1647 | 7.38E-05 | 0.00132664 | 0.587587698 |
| TOX | 1154 | 3.03E-06 | 7.78E-05 | 0.586392961 |
| PARP14 | 2087 | 0.00047457 | 0.00672776 | 0.586148774 |
| GCSAM | 697 | 1.12E-08 | 4.75E-07 | 0.58611601 |
| OAS3 | 1656 | 7.65E-05 | 0.00136653 | 0.585510267 |
| HAUS5 | 1122 | 2.14E-06 | 5.65E-05 | 0.584207782 |
| NDFIP1 | 300 | 3.73E-15 | 3.68E-13 | 0.583832467 |
| SERF2 | 1109 | 1.86E-06 | 4.96E-05 | 0.583455356 |
| IGF1R | 1159 | 3.22E-06 | 8.23E-05 | 0.581448703 |
| ETV5 | 596 | 1.12E-09 | 5.57E-08 | 0.581241293 |
| RAB3IP | 449 | 1.68E-11 | 1.11E-09 | 0.580421004 |
| AGPAT2 | 967 | 4.21E-07 | 1.29E-05 | 0.580338918 |
| GSDME | 643 | 3.55E-09 | 1.64E-07 | 0.580285313 |

Table_S2_PovertyEx_vs_UnExp_dow

| | Top | p.value | FDR | logFC |
|---|---|---|---|---|
| LSP1 | 4 | 8.94E-74 | 6.61E-70 | -3.037010854 |
| FLT3 | 13 | 5.24E-55 | 1.19E-51 | -2.785873788 |
| S100A16 | 16 | 1.47E-51 | 2.73E-48 | -2.70382188 |
| ANXA2 | 6 | 2.56E-65 | 1.26E-61 | -2.499674632 |
| CD9 | 66 | 9.74E-33 | 4.36E-30 | -2.4403548 |
| DDIT4 | 7 | 2.92E-63 | 1.24E-59 | -2.406694883 |
| PLP2 | 3 | 1.08E-74 | 1.07E-70 | -2.388321336 |
| CD164 | 48 | 2.33E-36 | 1.44E-33 | -2.370690011 |
| FLNA | 8 | 4.46E-63 | 1.65E-59 | -2.368678289 |
| IRF8 | 25 | 3.09E-43 | 3.66E-40 | -2.306257139 |
| PDLIM1 | 61 | 2.04E-33 | 9.92E-31 | -2.290478335 |
| SRGN | 9 | 6.12E-62 | 2.01E-58 | -2.126843382 |
| IL3RA | 45 | 7.49E-37 | 4.93E-34 | -2.020848937 |
| DENND3 | 80 | 2.42E-30 | 8.95E-28 | -2.001652962 |
| PLEK | 24 | 2.81E-43 | 3.46E-40 | -1.980562594 |
| CD44 | 36 | 3.88E-39 | 3.19E-36 | -1.957697578 |
| MOB3A | 26 | 6.30E-43 | 7.17E-40 | -1.935162891 |
| KLF6 | 147 | 3.05E-22 | 6.13E-20 | -1.902254304 |
| MICAL1 | 79 | 2.18E-30 | 8.15E-28 | -1.856455652 |
| MS4A6A | 20 | 4.79E-46 | 7.09E-43 | -1.845336572 |
| CYTH1 | 63 | 3.44E-33 | 1.62E-30 | -1.836564477 |
| LYST | 55 | 2.98E-34 | 1.60E-31 | -1.834312628 |
| CDKN1A | 40 | 7.55E-38 | 5.58E-35 | -1.827575774 |
| VIM | 98 | 5.35E-27 | 1.62E-24 | -1.807283593 |
| FOS | 219 | 4.28E-18 | 5.78E-16 | -1.787216272 |
| CD82 | 70 | 9.75E-32 | 4.09E-29 | -1.763293747 |
| LCP1 | 171 | 7.41E-21 | 1.28E-18 | -1.719521263 |
| ACTN1 | 37 | 9.32E-39 | 7.46E-36 | -1.71837951 |
| FLNB | 59 | 1.50E-33 | 7.53E-31 | -1.676566984 |
| GAS7 | 39 | 1.39E-38 | 1.06E-35 | -1.667218342 |
| STK17B | 118 | 1.77E-24 | 4.43E-22 | -1.602343109 |
| SH3BP2 | 94 | 9.59E-28 | 3.02E-25 | -1.597100745 |
| CHST12 | 34 | 3.48E-40 | 3.03E-37 | -1.596265771 |
| CYBB | 88 | 1.48E-28 | 4.97E-26 | -1.577298014 |
| NKG7 | 28 | 7.56E-43 | 7.99E-40 | -1.560447585 |
| IL2RG | 191 | 1.74E-19 | 2.70E-17 | -1.557554561 |
| IL17RA | 56 | 7.01E-34 | 3.70E-31 | -1.555333274 |
| USP36 | 101 | 4.20E-26 | 1.23E-23 | -1.540252452 |
| KCNAB2 | 42 | 2.98E-37 | 2.10E-34 | -1.528223243 |
| BLK | 81 | 3.22E-30 | 1.18E-27 | -1.52367489 |
| ST3GAL1 | 162 | 1.50E-21 | 2.73E-19 | -1.515740927 |

| | | | | |
|---|---|---|---|---|
| NIBAN3 | 157 | 8.56E-22 | 1.61E-19 | -1.509113491 |
| RAP1GAP2 | 71 | 9.82E-32 | 4.09E-29 | -1.508486634 |
| NFE2 | 32 | 5.78E-41 | 5.34E-38 | -1.496496826 |
| FUT7 | 38 | 1.04E-38 | 8.09E-36 | -1.496394939 |
| LAT2 | 272 | 3.07E-16 | 3.34E-14 | -1.492214044 |
| KLF10 | 160 | 1.46E-21 | 2.70E-19 | -1.491762939 |
| TSC22D3 | 327 | 2.14E-14 | 1.94E-12 | -1.490801334 |
| CYTIP | 52 | 6.87E-35 | 3.91E-32 | -1.489268962 |
| EVI2B | 127 | 1.69E-23 | 3.95E-21 | -1.488360528 |
| DDIT4L | 57 | 7.53E-34 | 3.91E-31 | -1.474026768 |
| SERPINB1 | 231 | 1.16E-17 | 1.49E-15 | -1.473194233 |
| NFKBIZ | 95 | 1.19E-27 | 3.72E-25 | -1.462027386 |
| MYT1L | 60 | 1.87E-33 | 9.20E-31 | -1.443982286 |
| BLNK | 233 | 1.53E-17 | 1.94E-15 | -1.442843817 |
| MGAT4A | 154 | 6.54E-22 | 1.26E-19 | -1.428915635 |
| P2RX5 | 53 | 8.47E-35 | 4.73E-32 | -1.411313362 |
| HSH2D | 213 | 2.59E-18 | 3.60E-16 | -1.406955251 |
| ITGB2 | 137 | 6.52E-23 | 1.41E-20 | -1.390869545 |
| PDE4B | 239 | 2.39E-17 | 2.96E-15 | -1.386062961 |
| OGT | 223 | 5.70E-18 | 7.56E-16 | -1.378129255 |
| UBE2G2 | 211 | 2.32E-18 | 3.25E-16 | -1.375524286 |
| FKBP5 | 215 | 2.68E-18 | 3.69E-16 | -1.373843817 |
| NBEAL2 | 73 | 4.75E-31 | 1.93E-28 | -1.358727923 |
| MYO1F | 119 | 3.21E-24 | 7.99E-22 | -1.355089286 |
| PPM1F | 136 | 6.35E-23 | 1.38E-20 | -1.349421864 |
| PTTG1IP | 255 | 8.68E-17 | 1.01E-14 | -1.343621907 |
| NAV1 | 269 | 2.21E-16 | 2.43E-14 | -1.340937647 |
| S100A4 | 51 | 9.59E-36 | 5.57E-33 | -1.339679484 |
| CORO1A | 288 | 1.57E-15 | 1.62E-13 | -1.326651137 |
| SAT1 | 379 | 3.78E-13 | 2.95E-11 | -1.32616442 |
| LGALS3BP | 296 | 3.33E-15 | 3.32E-13 | -1.310829748 |
| STX7 | 268 | 2.15E-16 | 2.37E-14 | -1.305605801 |
| STING1 | 139 | 6.81E-23 | 1.45E-20 | -1.303040016 |
| GLUL | 210 | 2.01E-18 | 2.83E-16 | -1.298055639 |
| CNN2 | 184 | 3.30E-20 | 5.31E-18 | -1.297134974 |
| IQGAP2 | 141 | 9.58E-23 | 2.01E-20 | -1.284231346 |
| BCLAF1 | 333 | 2.99E-14 | 2.66E-12 | -1.282725261 |
| PPA1 | 135 | 5.44E-23 | 1.19E-20 | -1.279695157 |
| C12orf75 | 128 | 2.12E-23 | 4.90E-21 | -1.277293482 |
| MAP2K3 | 237 | 1.87E-17 | 2.34E-15 | -1.268549285 |
| RFLNB | 251 | 6.95E-17 | 8.20E-15 | -1.265803905 |
| TASL | 212 | 2.39E-18 | 3.34E-16 | -1.258072974 |
| IFITM3 | 254 | 8.35E-17 | 9.73E-15 | -1.255464932 |

| | | | | |
|---|---|---|---|---|
| AMD1 | 238 | 1.97E-17 | 2.45E-15 | -1.247352994 |
| VSIR | 178 | 1.14E-20 | 1.89E-18 | -1.226755151 |
| RAB37 | 246 | 4.80E-17 | 5.76E-15 | -1.207322107 |
| ANTXR2 | 83 | 4.30E-30 | 1.53E-27 | -1.20654684 |
| POU2F2 | 96 | 1.95E-27 | 6.01E-25 | -1.196023024 |
| FERMT3 | 319 | 1.22E-14 | 1.13E-12 | -1.194494399 |
| PTPN6 | 371 | 2.50E-13 | 1.99E-11 | -1.194235974 |
| ADGRG1 | 228 | 7.39E-18 | 9.59E-16 | -1.190413936 |
| C20orf27 | 65 | 9.53E-33 | 4.34E-30 | -1.179693533 |
| MSN | 408 | 2.30E-12 | 1.67E-10 | -1.178927142 |
| CAPG | 323 | 1.89E-14 | 1.73E-12 | -1.176902003 |
| PSAP | 520 | 1.70E-10 | 9.65E-09 | -1.173922497 |
| EVL | 284 | 1.04E-15 | 1.08E-13 | -1.165771317 |
| KLF2 | 129 | 2.26E-23 | 5.19E-21 | -1.159842763 |
| PLXNB2 | 270 | 2.74E-16 | 3.00E-14 | -1.159104308 |
| FOSL2 | 86 | 2.39E-29 | 8.22E-27 | -1.155027747 |
| SAMHD1 | 172 | 7.98E-21 | 1.37E-18 | -1.153136398 |
| ADGRE2 | 109 | 2.54E-25 | 6.89E-23 | -1.147734607 |
| SIK1 | 111 | 4.61E-25 | 1.23E-22 | -1.147176041 |
| B4GALT1 | 262 | 1.40E-16 | 1.58E-14 | -1.145211758 |
| PRNP | 277 | 5.84E-16 | 6.24E-14 | -1.137826795 |
| IGBP1 | 409 | 2.33E-12 | 1.69E-10 | -1.137447284 |
| RHBDF2 | 165 | 2.66E-21 | 4.76E-19 | -1.135776269 |
| PLVAP | 75 | 1.21E-30 | 4.77E-28 | -1.134963208 |
| CSF3R | 306 | 5.51E-15 | 5.33E-13 | -1.12756384 |
| SLC2A3 | 264 | 1.72E-16 | 1.93E-14 | -1.124508576 |
| CTNND1 | 152 | 5.45E-22 | 1.06E-19 | -1.121483856 |
| SH3BP5 | 100 | 3.51E-26 | 1.04E-23 | -1.112449542 |
| ANXA7 | 462 | 2.21E-11 | 1.41E-09 | -1.09460745 |
| SLC2A5 | 123 | 6.65E-24 | 1.60E-21 | -1.094405012 |
| ITGAE | 294 | 2.27E-15 | 2.28E-13 | -1.090463919 |
| SOD2 | 456 | 1.86E-11 | 1.21E-09 | -1.081207661 |
| CAST | 276 | 5.41E-16 | 5.80E-14 | -1.076812995 |
| RASD1 | 331 | 2.85E-14 | 2.55E-12 | -1.075718004 |
| LINC00707 | 92 | 7.98E-28 | 2.57E-25 | -1.071836793 |
| ADGRE5 | 480 | 4.00E-11 | 2.47E-09 | -1.071702616 |
| CSGALNACT1 | 359 | 1.26E-13 | 1.04E-11 | -1.069816003 |
| PFKFB3 | 121 | 3.91E-24 | 9.56E-22 | -1.065009189 |
| CD69 | 828 | 7.98E-08 | 2.85E-06 | -1.063660268 |
| LILRB2 | 344 | 5.48E-14 | 4.71E-12 | -1.056987334 |
| MX1 | 775 | 3.93E-08 | 1.50E-06 | -1.05457155 |
| C9orf72 | 261 | 1.22E-16 | 1.39E-14 | -1.050352556 |
| SFMBT2 | 273 | 3.19E-16 | 3.46E-14 | -1.045108806 |

| | | | | |
|---|---|---|---|---|
| MAP3K8 | 267 | 2.08E-16 | 2.31E-14 | -1.044003812 |
| STAB1 | 89 | 1.76E-28 | 5.84E-26 | -1.03830206 |
| ZFP36 | 410 | 2.36E-12 | 1.70E-10 | -1.033899005 |
| PROM1 | 279 | 6.27E-16 | 6.65E-14 | -1.026965497 |
| XIST | 504 | 9.08E-11 | 5.33E-09 | -1.024960023 |
| SPOCK2 | 133 | 3.32E-23 | 7.40E-21 | -1.023071715 |
| LIMD2 | 252 | 7.71E-17 | 9.05E-15 | -1.01080629 |
| DYNLT1 | 351 | 7.91E-14 | 6.67E-12 | -1.010245675 |
| C15orf39 | 90 | 2.69E-28 | 8.83E-26 | -1.008481905 |
| DDX21 | 455 | 1.85E-11 | 1.20E-09 | -1.005840816 |
| TCP1 | 722 | 1.83E-08 | 7.50E-07 | -0.991620661 |
| GPM6B | 376 | 3.50E-13 | 2.75E-11 | -0.988771329 |
| RBM3 | 732 | 2.16E-08 | 8.73E-07 | -0.983614173 |
| IGHD | 375 | 3.34E-13 | 2.64E-11 | -0.983208009 |
| FAM53B | 310 | 6.84E-15 | 6.53E-13 | -0.977843393 |
| SNHG5 | 435 | 9.69E-12 | 6.59E-10 | -0.974850846 |
| VNN1 | 99 | 1.64E-26 | 4.91E-24 | -0.973750273 |
| SYNGR1 | 507 | 9.78E-11 | 5.71E-09 | -0.970946453 |
| RNF24 | 332 | 2.90E-14 | 2.58E-12 | -0.97083995 |
| PREX1 | 465 | 2.58E-11 | 1.64E-09 | -0.970039153 |
| SLITRK4 | 174 | 9.21E-21 | 1.57E-18 | -0.967849468 |
| TACC3 | 325 | 2.03E-14 | 1.85E-12 | -0.967465017 |
| LAP3 | 649 | 3.81E-09 | 1.74E-07 | -0.963729449 |
| GOLIM4 | 148 | 4.07E-22 | 8.14E-20 | -0.961756217 |
| IL6R | 113 | 8.32E-25 | 2.18E-22 | -0.954943421 |
| EZR | 908 | 2.02E-07 | 6.58E-06 | -0.95475307 |
| TALDO1 | 744 | 2.57E-08 | 1.02E-06 | -0.952832802 |
| OGFRL1 | 313 | 7.99E-15 | 7.56E-13 | -0.948563565 |
| JPT1 | 549 | 3.38E-10 | 1.82E-08 | -0.946908111 |
| RIPOR2 | 675 | 6.76E-09 | 2.96E-07 | -0.94639335 |
| PFKP | 299 | 3.58E-15 | 3.54E-13 | -0.943507837 |
| PDCD4 | 514 | 1.44E-10 | 8.28E-09 | -0.941554729 |
| ARMH1 | 493 | 6.04E-11 | 3.63E-09 | -0.941038424 |
| CSGALNACT2 | 140 | 9.12E-23 | 1.93E-20 | -0.940520853 |
| NDE1 | 422 | 4.88E-12 | 3.42E-10 | -0.939515386 |
| AP1S2 | 575 | 6.94E-10 | 3.57E-08 | -0.935586397 |
| GLIPR1 | 348 | 6.96E-14 | 5.92E-12 | -0.933130793 |
| VPS26A | 486 | 5.00E-11 | 3.04E-09 | -0.931124985 |
| SERPINF1 | 199 | 4.52E-19 | 6.71E-17 | -0.931081527 |
| C18orf63 | 201 | 6.27E-19 | 9.23E-17 | -0.930738923 |
| ASCC1 | 342 | 4.44E-14 | 3.84E-12 | -0.928694166 |
| FAM30A | 186 | 4.55E-20 | 7.24E-18 | -0.925306988 |
| RASGRP2 | 321 | 1.71E-14 | 1.57E-12 | -0.925117859 |

| | | | | |
|---|---|---|---|---|
| NIN | 620 | 2.08E-09 | 9.95E-08 | -0.92310915 |
| RNF125 | 290 | 1.85E-15 | 1.89E-13 | -0.922426395 |
| ECHDC1 | 454 | 1.84E-11 | 1.20E-09 | -0.922366595 |
| ADA | 774 | 3.87E-08 | 1.48E-06 | -0.919964091 |
| RNF130 | 378 | 3.71E-13 | 2.91E-11 | -0.919956033 |
| EVI2A | 347 | 6.75E-14 | 5.76E-12 | -0.919248648 |
| LRRFIP1 | 509 | 1.05E-10 | 6.08E-09 | -0.91835941 |
| ARFGAP3 | 458 | 1.88E-11 | 1.21E-09 | -0.915413077 |
| AHNAK | 214 | 2.61E-18 | 3.60E-16 | -0.914437247 |
| EMP3 | 151 | 5.41E-22 | 1.06E-19 | -0.907154545 |
| RAB34 | 476 | 3.53E-11 | 2.20E-09 | -0.904803731 |
| GNA15 | 413 | 2.83E-12 | 2.03E-10 | -0.900281002 |
| TMED10 | 843 | 1.00E-07 | 3.52E-06 | -0.900177562 |
| SRP72 | 812 | 6.21E-08 | 2.26E-06 | -0.900084819 |
| XAF1 | 1079 | 1.33E-06 | 3.65E-05 | -0.899881343 |
| GRB2 | 600 | 1.35E-09 | 6.68E-08 | -0.89945424 |
| TNFAIP3 | 781 | 4.07E-08 | 1.54E-06 | -0.89790331 |
| FRMD4B | 751 | 2.78E-08 | 1.09E-06 | -0.89394358 |
| GAB3 | 518 | 1.55E-10 | 8.87E-09 | -0.893478893 |
| ANKRD44 | 432 | 9.32E-12 | 6.38E-10 | -0.8929276 |
| PLEC | 384 | 4.46E-13 | 3.44E-11 | -0.892442443 |
| APAF1 | 250 | 5.91E-17 | 7.00E-15 | -0.890961739 |
| WDFY4 | 505 | 9.17E-11 | 5.37E-09 | -0.889366757 |
| ENTPD1 | 235 | 1.78E-17 | 2.24E-15 | -0.880364395 |
| BRF1 | 403 | 1.74E-12 | 1.28E-10 | -0.878364422 |
| SH2D3C | 719 | 1.68E-08 | 6.91E-07 | -0.877038444 |
| NUFIP2 | 636 | 3.13E-09 | 1.46E-07 | -0.873633495 |
| PTP4A1 | 911 | 2.12E-07 | 6.89E-06 | -0.872592207 |
| SLC25A37 | 713 | 1.43E-08 | 5.94E-07 | -0.870355331 |
| CANX | 992 | 5.34E-07 | 1.59E-05 | -0.869467443 |
| GSTO1 | 463 | 2.30E-11 | 1.47E-09 | -0.86900805 |
| PLIN2 | 829 | 8.11E-08 | 2.89E-06 | -0.865321557 |
| PPP1R15A | 921 | 2.40E-07 | 7.70E-06 | -0.864456777 |
| SERINC1 | 760 | 3.11E-08 | 1.21E-06 | -0.86442597 |
| BTAF1 | 638 | 3.18E-09 | 1.47E-07 | -0.859579766 |
| SARAF | 1087 | 1.47E-06 | 3.99E-05 | -0.85917493 |
| NDRG1 | 153 | 6.49E-22 | 1.26E-19 | -0.858050158 |
| PMAIP1 | 1064 | 1.18E-06 | 3.29E-05 | -0.85776327 |
| METTL7A | 383 | 4.14E-13 | 3.20E-11 | -0.856742742 |
| ATP8A1 | 326 | 2.10E-14 | 1.91E-12 | -0.855273817 |
| NFATC3 | 537 | 2.56E-10 | 1.41E-08 | -0.85351008 |
| CARD19 | 200 | 4.82E-19 | 7.13E-17 | -0.852909206 |
| SNX27 | 709 | 1.36E-08 | 5.70E-07 | -0.852451623 |

| | | | | |
|---|---|---|---|---|
| ITGA4 | 919 | 2.35E-07 | 7.56E-06 | -0.851602706 |
| SNX6 | 808 | 5.92E-08 | 2.17E-06 | -0.850514923 |
| TMX1 | 580 | 7.79E-10 | 3.97E-08 | -0.850229378 |
| CTSS | 863 | 1.31E-07 | 4.49E-06 | -0.849659772 |
| ALOX5 | 963 | 3.95E-07 | 1.21E-05 | -0.848953113 |
| SMAD3 | 350 | 7.89E-14 | 6.67E-12 | -0.848734111 |
| SLC7A5 | 358 | 1.14E-13 | 9.44E-12 | -0.848226374 |
| RRP1B | 489 | 5.57E-11 | 3.37E-09 | -0.84479768 |
| S100A6 | 146 | 2.24E-22 | 4.53E-20 | -0.843946178 |
| ABCA2 | 236 | 1.81E-17 | 2.27E-15 | -0.843453873 |
| CCNC | 725 | 1.85E-08 | 7.56E-07 | -0.841362578 |
| MYC | 337 | 3.81E-14 | 3.34E-12 | -0.838674132 |
| ALDH3B1 | 159 | 1.22E-21 | 2.28E-19 | -0.837954368 |
| SMAGP | 354 | 8.79E-14 | 7.35E-12 | -0.833733975 |
| MGAT1 | 802 | 5.28E-08 | 1.95E-06 | -0.833587487 |
| SPPL2B | 586 | 9.20E-10 | 4.65E-08 | -0.833502448 |
| ANKRD28 | 651 | 4.22E-09 | 1.92E-07 | -0.833466438 |
| ADAM8 | 282 | 7.88E-16 | 8.26E-14 | -0.831600426 |
| NEAT1 | 658 | 4.74E-09 | 2.13E-07 | -0.831311845 |
| LINC-PINT | 308 | 6.14E-15 | 5.90E-13 | -0.831166856 |
| BTNL9 | 234 | 1.76E-17 | 2.23E-15 | -0.830901506 |
| NR4A2 | 291 | 1.88E-15 | 1.91E-13 | -0.830577143 |
| HBEGF | 385 | 4.57E-13 | 3.51E-11 | -0.829346574 |
| ARHGEF10 | 561 | 4.52E-10 | 2.38E-08 | -0.827006829 |
| PAN3 | 842 | 9.78E-08 | 3.44E-06 | -0.826229674 |
| NCF2 | 582 | 7.98E-10 | 4.06E-08 | -0.82460508 |
| CPNE3 | 805 | 5.68E-08 | 2.09E-06 | -0.81691377 |
| ASAP1 | 792 | 4.66E-08 | 1.74E-06 | -0.816491208 |
| SMIM24 | 374 | 3.12E-13 | 2.47E-11 | -0.814468255 |
| FHL1 | 1150 | 2.89E-06 | 7.42E-05 | -0.812067671 |
| DYNC1H1 | 920 | 2.35E-07 | 7.56E-06 | -0.811208805 |
| JAK3 | 602 | 1.39E-09 | 6.83E-08 | -0.811087546 |
| NAPSB | 198 | 3.91E-19 | 5.84E-17 | -0.808921224 |
| HSP90AB1 | 1423 | 2.36E-05 | 0.00049146 | -0.807642663 |
| SMAD2 | 855 | 1.22E-07 | 4.23E-06 | -0.807155299 |
| CYTH4 | 862 | 1.28E-07 | 4.40E-06 | -0.80617889 |
| TRPM2 | 687 | 8.42E-09 | 3.63E-07 | -0.805759854 |
| TIMP1 | 464 | 2.50E-11 | 1.59E-09 | -0.805197591 |
| CFP | 189 | 1.23E-19 | 1.93E-17 | -0.804611733 |
| UBQLN2 | 830 | 8.32E-08 | 2.97E-06 | -0.804196124 |
| ATP2B4 | 960 | 3.91E-07 | 1.21E-05 | -0.802584612 |
| MAPKBP1 | 388 | 5.10E-13 | 3.89E-11 | -0.801954554 |
| CASP1 | 302 | 4.15E-15 | 4.07E-13 | -0.801530904 |

| | | | | |
|---|---|---|---|---|
| ATP6V0D1 | 836 | 8.99E-08 | 3.18E-06 | -0.797169977 |
| CCND2 | 1060 | 1.14E-06 | 3.19E-05 | -0.79610333 |
| SCPEP1 | 831 | 8.45E-08 | 3.01E-06 | -0.795656334 |
| MYH11 | 345 | 5.52E-14 | 4.74E-12 | -0.795080847 |
| CD34 | 1085 | 1.44E-06 | 3.93E-05 | -0.795046165 |
| VMP1 | 901 | 1.89E-07 | 6.22E-06 | -0.794378175 |
| TNFSF13B | 161 | 1.48E-21 | 2.72E-19 | -0.793011166 |
| GAS5 | 981 | 4.78E-07 | 1.44E-05 | -0.792045339 |
| ETS2 | 1131 | 2.37E-06 | 6.19E-05 | -0.791224381 |
| PGD | 1028 | 8.41E-07 | 2.42E-05 | -0.789410125 |
| SEC14L1 | 395 | 8.66E-13 | 6.49E-11 | -0.788897954 |
| CD55 | 809 | 5.93E-08 | 2.17E-06 | -0.788154227 |
| UGT3A2 | 418 | 4.23E-12 | 2.99E-10 | -0.786143261 |
| EIF4A1 | 542 | 2.91E-10 | 1.59E-08 | -0.785350737 |
| SNHG29 | 710 | 1.39E-08 | 5.78E-07 | -0.78515897 |
| SLC9A3R1 | 392 | 6.50E-13 | 4.91E-11 | -0.78373007 |
| RIOK1 | 527 | 1.90E-10 | 1.07E-08 | -0.781648166 |
| SF3B5 | 508 | 1.00E-10 | 5.84E-09 | -0.780763063 |
| TRBC2 | 155 | 6.70E-22 | 1.28E-19 | -0.780538342 |
| TAGLN2 | 894 | 1.80E-07 | 5.96E-06 | -0.779056226 |
| PML | 584 | 8.46E-10 | 4.29E-08 | -0.778144236 |
| TSC22D1 | 791 | 4.56E-08 | 1.71E-06 | -0.774506229 |
| SYNCRIP | 1096 | 1.58E-06 | 4.28E-05 | -0.773023018 |
| KIF1B | 275 | 4.90E-16 | 5.27E-14 | -0.771360326 |
| USO1 | 748 | 2.76E-08 | 1.09E-06 | -0.771066231 |
| NPC2 | 619 | 2.08E-09 | 9.95E-08 | -0.768712198 |
| UBE2J1 | 1023 | 7.55E-07 | 2.18E-05 | -0.76645075 |
| PTPRC | 541 | 2.89E-10 | 1.58E-08 | -0.766326858 |
| FXYD5 | 970 | 4.42E-07 | 1.35E-05 | -0.765758264 |
| SETBP1 | 646 | 3.72E-09 | 1.71E-07 | -0.763894429 |
| GRN | 1160 | 3.24E-06 | 8.27E-05 | -0.763498108 |
| DYM | 940 | 2.88E-07 | 9.06E-06 | -0.762621388 |
| DDX3X | 1294 | 1.07E-05 | 0.00024472 | -0.761826104 |
| ATP2C1 | 367 | 2.19E-13 | 1.77E-11 | -0.759543525 |
| BIN2 | 221 | 5.04E-18 | 6.74E-16 | -0.759401752 |
| EIF4A3 | 955 | 3.68E-07 | 1.14E-05 | -0.756298256 |
| CLIP2 | 632 | 2.79E-09 | 1.31E-07 | -0.754342672 |
| KRI1 | 1062 | 1.16E-06 | 3.22E-05 | -0.754180303 |
| C1QBP | 1092 | 1.54E-06 | 4.17E-05 | -0.752974781 |
| SH3RF1 | 703 | 1.24E-08 | 5.23E-07 | -0.752932634 |
| LASP1 | 897 | 1.86E-07 | 6.12E-06 | -0.751919608 |
| ATP6AP2 | 1262 | 8.34E-06 | 0.00019539 | -0.751740858 |
| FES | 888 | 1.73E-07 | 5.77E-06 | -0.75121898 |

| | | | | |
|---|---|---|---|---|
| UTP6 | 984 | 4.91E-07 | 1.48E-05 | -0.750992514 |
| TINF2 | 1105 | 1.78E-06 | 4.77E-05 | -0.750601998 |
| EEF1B2 | 995 | 5.42E-07 | 1.61E-05 | -0.74875971 |
| MFSD1 | 803 | 5.42E-08 | 2.00E-06 | -0.74834106 |
| RAB9A | 603 | 1.44E-09 | 7.05E-08 | -0.747409239 |
| FLII | 1117 | 2.09E-06 | 5.55E-05 | -0.746464256 |
| CNN3 | 515 | 1.44E-10 | 8.28E-09 | -0.745559717 |
| OFD1 | 1339 | 1.49E-05 | 0.00032899 | -0.745322713 |
| HSDL2 | 411 | 2.36E-12 | 1.70E-10 | -0.742233738 |
| HOOK3 | 785 | 4.39E-08 | 1.65E-06 | -0.740451243 |
| ADPGK | 1047 | 9.69E-07 | 2.74E-05 | -0.738423705 |
| NUP210 | 572 | 5.49E-10 | 2.84E-08 | -0.738043388 |
| SNHG3 | 917 | 2.30E-07 | 7.41E-06 | -0.735715427 |
| SNHG8 | 705 | 1.27E-08 | 5.34E-07 | -0.733716372 |
| PQBP1 | 761 | 3.14E-08 | 1.22E-06 | -0.733139375 |
| GATB | 416 | 3.81E-12 | 2.71E-10 | -0.732794608 |
| GNG2 | 918 | 2.32E-07 | 7.49E-06 | -0.732292657 |
| NLRP1 | 1111 | 1.93E-06 | 5.13E-05 | -0.731326972 |
| PKIG | 280 | 6.88E-16 | 7.27E-14 | -0.730685206 |
| ACAT1 | 769 | 3.51E-08 | 1.35E-06 | -0.730478509 |
| HDAC9 | 437 | 9.83E-12 | 6.65E-10 | -0.73022244 |
| MS4A4E | 125 | 8.63E-24 | 2.04E-21 | -0.729548025 |
| MED23 | 814 | 6.24E-08 | 2.27E-06 | -0.729360871 |
| PSMC6 | 1174 | 3.80E-06 | 9.57E-05 | -0.728989749 |
| STAT3 | 1090 | 1.52E-06 | 4.14E-05 | -0.728751281 |
| SUN1 | 890 | 1.76E-07 | 5.84E-06 | -0.726116773 |
| PRSS57 | 124 | 7.80E-24 | 1.86E-21 | -0.725594609 |
| ABCE1 | 633 | 2.81E-09 | 1.31E-07 | -0.724741497 |
| B3GNTL1 | 445 | 1.31E-11 | 8.74E-10 | -0.724476513 |
| TCOF1 | 1226 | 5.90E-06 | 0.00014248 | -0.723779007 |
| HVCN1 | 544 | 3.05E-10 | 1.66E-08 | -0.722516666 |
| RABGAP1L | 910 | 2.11E-07 | 6.86E-06 | -0.721886662 |
| PIK3R5 | 762 | 3.15E-08 | 1.22E-06 | -0.7214575 |
| USP9X | 1255 | 7.62E-06 | 0.00017972 | -0.719184943 |
| DUSP3 | 304 | 4.59E-15 | 4.47E-13 | -0.718479548 |
| TRAF3IP3 | 825 | 7.54E-08 | 2.71E-06 | -0.717601337 |
| PDE3B | 324 | 2.00E-14 | 1.83E-12 | -0.716642101 |
| SEMA4A | 257 | 9.21E-17 | 1.06E-14 | -0.715425368 |
| KCNK17 | 316 | 1.08E-14 | 1.01E-12 | -0.714788856 |
| CALHM2 | 421 | 4.88E-12 | 3.42E-10 | -0.714615478 |
| GHITM | 1764 | 0.00012495 | 0.00209604 | -0.712424748 |
| NUMB | 1355 | 1.61E-05 | 0.00035087 | -0.711804367 |
| SH3KBP1 | 1167 | 3.39E-06 | 8.60E-05 | -0.710577432 |

| | | | | |
|---|---|---|---|---|
| PNISR | 1158 | 3.22E-06 | 8.22E-05 | -0.709337912 |
| PSMB1 | 1537 | 4.32E-05 | 0.00083079 | -0.709321304 |
| TSPAN13 | 1197 | 4.54E-06 | 0.00011231 | -0.707271715 |
| ITSN1 | 488 | 5.20E-11 | 3.16E-09 | -0.706831068 |
| LATS2 | 402 | 1.33E-12 | 9.82E-11 | -0.706652106 |
| TMEM123 | 1631 | 6.90E-05 | 0.00125231 | -0.704190277 |
| PDXK | 373 | 2.96E-13 | 2.35E-11 | -0.702132113 |
| GSTP1 | 1194 | 4.47E-06 | 0.00011068 | -0.700178653 |
| GPR132 | 571 | 5.49E-10 | 2.84E-08 | -0.699665197 |
| CCDC200 | 937 | 2.71E-07 | 8.56E-06 | -0.699484809 |
| AFF3 | 1228 | 6.01E-06 | 0.00014488 | -0.699262662 |
| CNIH1 | 965 | 4.07E-07 | 1.25E-05 | -0.698653319 |
| AKNA | 1024 | 7.69E-07 | 2.22E-05 | -0.698198977 |
| ELAC2 | 998 | 5.65E-07 | 1.67E-05 | -0.69656593 |
| PNP | 640 | 3.38E-09 | 1.56E-07 | -0.6952331 |
| ABR | 540 | 2.82E-10 | 1.54E-08 | -0.695195415 |
| ZYX | 227 | 7.16E-18 | 9.33E-16 | -0.694819825 |
| PSEN1 | 1089 | 1.49E-06 | 4.06E-05 | -0.694459975 |
| RGL2 | 816 | 6.43E-08 | 2.33E-06 | -0.694259305 |
| IL1B | 1666 | 8.10E-05 | 0.00143817 | -0.694165606 |
| C9orf139 | 206 | 1.20E-18 | 1.73E-16 | -0.694098726 |
| WAPL | 585 | 9.19E-10 | 4.65E-08 | -0.693623392 |
| MED13L | 1137 | 2.49E-06 | 6.49E-05 | -0.69287417 |
| LSM2 | 936 | 2.70E-07 | 8.55E-06 | -0.690406249 |
| ABCB7 | 1094 | 1.56E-06 | 4.21E-05 | -0.690307146 |
| SH3BGRL | 1867 | 0.00018397 | 0.00291538 | -0.6892195 |
| ADK | 663 | 5.26E-09 | 2.35E-07 | -0.687736503 |
| BTLA | 175 | 9.52E-21 | 1.61E-18 | -0.687484318 |
| LAMP2 | 1441 | 2.64E-05 | 0.0005411 | -0.687264893 |
| JMJD6 | 835 | 8.89E-08 | 3.15E-06 | -0.687074477 |
| HDAC6 | 899 | 1.89E-07 | 6.22E-06 | -0.686794667 |
| SERPINB6 | 1063 | 1.16E-06 | 3.24E-05 | -0.684662072 |
| C11orf21 | 177 | 1.12E-20 | 1.87E-18 | -0.684521543 |
| SLBP | 1166 | 3.36E-06 | 8.53E-05 | -0.683743316 |
| GCNT2 | 164 | 2.27E-21 | 4.09E-19 | -0.681496681 |
| ESYT1 | 756 | 2.92E-08 | 1.14E-06 | -0.681386747 |
| SETD3 | 1066 | 1.21E-06 | 3.36E-05 | -0.681383642 |
| CLEC2D | 959 | 3.88E-07 | 1.20E-05 | -0.680275313 |
| MYL12B | 1499 | 3.48E-05 | 0.00068754 | -0.67997955 |
| NUTM2A-AS | 576 | 7.22E-10 | 3.71E-08 | -0.678240556 |
| RPF2 | 439 | 1.04E-11 | 7.00E-10 | -0.676784047 |
| SPTLC2 | 1529 | 4.09E-05 | 0.00079235 | -0.675990072 |
| ADGRG5 | 176 | 1.06E-20 | 1.79E-18 | -0.675707565 |

| | | | | |
|---|---|---|---|---|
| ID2 | 1556 | 4.69E-05 | 0.00089179 | -0.6752458 |
| RBPJ | 1483 | 3.29E-05 | 0.00065741 | -0.674312412 |
| ARPC5 | 1840 | 0.00016802 | 0.00270208 | -0.673654368 |
| TMEM30A | 1053 | 1.02E-06 | 2.86E-05 | -0.672404876 |
| SNHG16 | 539 | 2.75E-10 | 1.51E-08 | -0.672169697 |
| FMNL1 | 727 | 1.88E-08 | 7.63E-07 | -0.67181691 |
| VAT1 | 750 | 2.77E-08 | 1.09E-06 | -0.671607966 |
| MOB3B | 581 | 7.88E-10 | 4.01E-08 | -0.670633272 |
| SYAP1 | 952 | 3.60E-07 | 1.12E-05 | -0.66961897 |
| GALNT3 | 372 | 2.91E-13 | 2.32E-11 | -0.669255147 |
| CCDC69 | 696 | 1.11E-08 | 4.71E-07 | -0.667466976 |
| EIF2A | 1276 | 9.37E-06 | 0.00021734 | -0.667325845 |
| ADD1 | 1435 | 2.54E-05 | 0.00052352 | -0.66732228 |
| COTL1 | 205 | 1.11E-18 | 1.60E-16 | -0.666972029 |
| MAML3 | 1051 | 1.00E-06 | 2.82E-05 | -0.666673121 |
| CTSB | 1519 | 3.90E-05 | 0.00075899 | -0.666441735 |
| RHOH | 1518 | 3.87E-05 | 0.00075497 | -0.666247535 |
| IVD | 1056 | 1.05E-06 | 2.95E-05 | -0.666155464 |
| LDHA | 2157 | 0.00059348 | 0.00814009 | -0.666089522 |
| MSL3 | 927 | 2.51E-07 | 8.01E-06 | -0.665311226 |
| LMO2 | 397 | 1.16E-12 | 8.62E-11 | -0.665246751 |
| CCDC26 | 168 | 4.62E-21 | 8.14E-19 | -0.663511449 |
| HDHD5 | 609 | 1.56E-09 | 7.59E-08 | -0.662748874 |
| ASPH | 369 | 2.46E-13 | 1.98E-11 | -0.662326604 |
| CTC1 | 1186 | 4.21E-06 | 0.0001051 | -0.66200198 |
| CRYZL1 | 1532 | 4.16E-05 | 0.00080313 | -0.660189288 |
| RSL1D1 | 1416 | 2.27E-05 | 0.00047414 | -0.659866828 |
| FTSJ3 | 1127 | 2.24E-06 | 5.89E-05 | -0.65867626 |
| MEF2D | 715 | 1.50E-08 | 6.21E-07 | -0.656221328 |
| KIN | 846 | 1.06E-07 | 3.72E-06 | -0.656212299 |
| STK10 | 1206 | 4.90E-06 | 0.00012017 | -0.655862215 |
| RASSF4 | 551 | 3.59E-10 | 1.93E-08 | -0.655394658 |
| USP38 | 536 | 2.55E-10 | 1.41E-08 | -0.654739641 |
| SH3BP5-AS1 | 564 | 4.71E-10 | 2.47E-08 | -0.654181255 |
| ARAF | 833 | 8.69E-08 | 3.09E-06 | -0.654176979 |
| GM2A | 525 | 1.77E-10 | 9.96E-09 | -0.65321021 |
| IDH3A | 453 | 1.81E-11 | 1.18E-09 | -0.652777123 |
| GDI2 | 1879 | 0.00019254 | 0.00303101 | -0.652343527 |
| TBC1D10C | 1180 | 3.83E-06 | 9.62E-05 | -0.651957544 |
| PRKD2 | 974 | 4.54E-07 | 1.38E-05 | -0.650675736 |
| QKI | 903 | 1.94E-07 | 6.34E-06 | -0.649928269 |
| STN1 | 674 | 6.73E-09 | 2.96E-07 | -0.648697323 |
| RGS18 | 412 | 2.68E-12 | 1.92E-10 | -0.647796295 |

| | | | | |
|---|---|---|---|---|
| SPECC1 | 550 | 3.53E-10 | 1.90E-08 | -0.646705765 |
| VCL | 1376 | 1.76E-05 | 0.00037888 | -0.646223431 |
| CD86 | 253 | 8.13E-17 | 9.51E-15 | -0.645093543 |
| CLEC2B | 777 | 3.95E-08 | 1.50E-06 | -0.643627912 |
| SBF1 | 823 | 7.26E-08 | 2.61E-06 | -0.643378273 |
| MACROH2A1 | 1249 | 7.28E-06 | 0.00017252 | -0.64325173 |
| HADHA | 1857 | 0.00017538 | 0.00279457 | -0.64232458 |
| CDH11 | 220 | 4.60E-18 | 6.18E-16 | -0.641404739 |
| CCDC12 | 1310 | 1.18E-05 | 0.000267 | -0.641371027 |
| TACC1 | 1371 | 1.73E-05 | 0.00037416 | -0.640770197 |
| UXT | 771 | 3.73E-08 | 1.43E-06 | -0.640583764 |
| TRPV2 | 691 | 8.99E-09 | 3.85E-07 | -0.640373657 |
| SFT2D1 | 997 | 5.59E-07 | 1.66E-05 | -0.640370452 |
| SSR1 | 1260 | 8.06E-06 | 0.00018924 | -0.640270793 |
| FOXO3 | 788 | 4.49E-08 | 1.69E-06 | -0.640228112 |
| ITGAL | 440 | 1.08E-11 | 7.28E-10 | -0.640120616 |
| MED14 | 1098 | 1.62E-06 | 4.36E-05 | -0.638945396 |
| CSTB | 938 | 2.78E-07 | 8.76E-06 | -0.638941041 |
| CYRIB | 1663 | 8.01E-05 | 0.00142462 | -0.638158205 |
| BNIP2 | 1665 | 8.09E-05 | 0.00143796 | -0.63669339 |
| MTHFD1 | 1072 | 1.28E-06 | 3.54E-05 | -0.635548762 |
| MAGT1 | 1401 | 2.06E-05 | 0.00043539 | -0.635503696 |
| RGS1 | 419 | 4.28E-12 | 3.02E-10 | -0.63540926 |
| HSP90B1 | 2047 | 0.00040261 | 0.00581998 | -0.635209568 |
| RLIM | 1169 | 3.59E-06 | 9.09E-05 | -0.634287955 |
| OLFML2A | 356 | 9.80E-14 | 8.14E-12 | -0.633968653 |
| PPP2R5C | 1628 | 6.79E-05 | 0.00123356 | -0.633926517 |
| CHD3 | 1083 | 1.40E-06 | 3.83E-05 | -0.633506322 |
| MYL12A | 1330 | 1.37E-05 | 0.00030502 | -0.633442882 |
| BCAT1 | 630 | 2.63E-09 | 1.24E-07 | -0.633065105 |
| TMCC3 | 216 | 3.09E-18 | 4.23E-16 | -0.631702081 |
| ANXA4 | 718 | 1.64E-08 | 6.77E-07 | -0.631390356 |
| RNASE6 | 244 | 3.20E-17 | 3.88E-15 | -0.630928928 |
| NDUFB11 | 1231 | 6.32E-06 | 0.00015199 | -0.630770222 |
| AHCTF1 | 887 | 1.72E-07 | 5.75E-06 | -0.628880871 |
| RBM25 | 1588 | 5.41E-05 | 0.00100739 | -0.628115573 |
| LPXN | 650 | 3.89E-09 | 1.77E-07 | -0.627921557 |
| RIN3 | 179 | 1.24E-20 | 2.04E-18 | -0.626002442 |
| HSPA5 | 2281 | 0.00084586 | 0.0109732 | -0.625586777 |
| IKZF3 | 352 | 8.42E-14 | 7.07E-12 | -0.625127005 |
| MYO15B | 1595 | 5.68E-05 | 0.00105285 | -0.624050025 |
| ANAPC16 | 1585 | 5.32E-05 | 0.00099276 | -0.623005049 |
| MED13 | 1544 | 4.42E-05 | 0.00084699 | -0.622889866 |

| | | | | |
|---|---|---|---|---|
| SYTL1 | 716 | 1.53E-08 | 6.32E-07 | -0.62180037 |
| DNAJC1 | 430 | 7.94E-12 | 5.46E-10 | -0.62161875 |
| DERL1 | 1241 | 6.93E-06 | 0.00016519 | -0.62103878 |
| DIPK1C | 768 | 3.47E-08 | 1.34E-06 | -0.620603374 |
| MS4A1 | 1375 | 1.76E-05 | 0.00037887 | -0.620319058 |
| WDR13 | 1130 | 2.34E-06 | 6.13E-05 | -0.620311371 |
| CCAR1 | 1756 | 0.00012091 | 0.00203752 | -0.61877468 |
| PPP2R5A | 844 | 1.03E-07 | 3.62E-06 | -0.617952331 |
| RAB11FIP1 | 543 | 3.01E-10 | 1.64E-08 | -0.617699667 |
| MTMR14 | 1271 | 9.15E-06 | 0.00021307 | -0.617221235 |
| RSU1 | 1009 | 6.37E-07 | 1.87E-05 | -0.617194183 |
| GNPTG | 941 | 2.89E-07 | 9.09E-06 | -0.616206109 |
| POLR2B | 1568 | 4.91E-05 | 0.00092699 | -0.614522974 |
| UVRAG | 625 | 2.39E-09 | 1.13E-07 | -0.614418831 |
| MAST3 | 647 | 3.74E-09 | 1.71E-07 | -0.614023636 |
| SH3TC1 | 1896 | 0.00021282 | 0.00332147 | -0.61385196 |
| TMBIM1 | 870 | 1.43E-07 | 4.87E-06 | -0.613542948 |
| ADCY7 | 292 | 2.19E-15 | 2.21E-13 | -0.613501332 |
| TLR1 | 796 | 4.82E-08 | 1.79E-06 | -0.613316631 |
| ASF1A | 1050 | 9.91E-07 | 2.79E-05 | -0.613082748 |
| NSMAF | 1193 | 4.47E-06 | 0.00011068 | -0.612944057 |
| PUDP | 654 | 4.31E-09 | 1.95E-07 | -0.612398693 |
| GINM1 | 731 | 1.98E-08 | 8.01E-07 | -0.61183909 |
| TNFAIP8L2 | 483 | 4.55E-11 | 2.79E-09 | -0.611474967 |
| LIPA | 1253 | 7.43E-06 | 0.0001755 | -0.611340303 |
| MTHFD2 | 641 | 3.38E-09 | 1.56E-07 | -0.609496249 |
| NPC1 | 1054 | 1.04E-06 | 2.92E-05 | -0.609196543 |
| NDRG3 | 1041 | 9.37E-07 | 2.66E-05 | -0.607819589 |
| EIF5A | 1120 | 2.12E-06 | 5.59E-05 | -0.607460313 |
| RSL24D1 | 1853 | 0.00017411 | 0.00278041 | -0.606135467 |
| ETF1 | 1433 | 2.51E-05 | 0.00051791 | -0.606102033 |
| IPCEF1 | 880 | 1.61E-07 | 5.42E-06 | -0.606049084 |
| WDR45B | 1514 | 3.85E-05 | 0.0007516 | -0.605695008 |
| HOPX | 271 | 2.96E-16 | 3.23E-14 | -0.605651737 |
| HDAC2 | 2177 | 0.00063963 | 0.00869421 | -0.60541108 |
| LINC01410 | 336 | 3.77E-14 | 3.32E-12 | -0.60474786 |
| PAICS | 1458 | 2.96E-05 | 0.00060116 | -0.604276804 |
| TP53 | 1290 | 1.04E-05 | 0.00023879 | -0.603329955 |
| DOCK11 | 1527 | 4.07E-05 | 0.00078816 | -0.603062626 |
| SLC43A2 | 450 | 1.68E-11 | 1.11E-09 | -0.602641662 |
| RUNX2 | 341 | 4.34E-14 | 3.77E-12 | -0.601847459 |
| SUSD3 | 431 | 8.34E-12 | 5.72E-10 | -0.601828054 |
| MARCHF5 | 1340 | 1.49E-05 | 0.00032984 | -0.601693112 |

| | | | | |
|---|---|---|---|---|
| MYD88 | 2083 | 0.00046493 | 0.00660476 | -0.600234941 |
| NLRC5 | 1757 | 0.00012125 | 0.00204213 | -0.599727404 |
| SHOC2 | 1767 | 0.00012603 | 0.00211048 | -0.599647955 |
| RNF149 | 1412 | 2.16E-05 | 0.00045166 | -0.598953081 |
| ASCC3 | 1110 | 1.90E-06 | 5.07E-05 | -0.597933871 |
| CD83 | 666 | 5.64E-09 | 2.51E-07 | -0.59785927 |
| CA5B | 706 | 1.30E-08 | 5.44E-07 | -0.597630317 |
| DAXX | 1562 | 4.78E-05 | 0.00090633 | -0.597450802 |
| NMD3 | 1002 | 5.92E-07 | 1.75E-05 | -0.597364842 |
| MTDH | 1486 | 3.31E-05 | 0.00066009 | -0.596597282 |
| ATG4C | 804 | 5.50E-08 | 2.02E-06 | -0.596423086 |
| MCL1 | 1082 | 1.39E-06 | 3.80E-05 | -0.594933947 |
| CCT8 | 2575 | 0.00166065 | 0.0190836 | -0.593556239 |
| LOC1009963[illegible] | 328 | 2.37E-14 | 2.14E-12 | -0.590948328 |
| EEF1B2P3 | 800 | 5.23E-08 | 1.94E-06 | -0.590143735 |
| ECM1 | 1778 | 0.00013029 | 0.00216835 | -0.588515432 |
| FPGS | 1915 | 0.0002259 | 0.00349063 | -0.588403385 |
| ADAM10 | 1800 | 0.00014235 | 0.00234023 | -0.587818332 |
| NEK9 | 1484 | 3.30E-05 | 0.00065749 | -0.587317753 |
| EIF3B | 1224 | 5.77E-06 | 0.00013954 | -0.586723881 |
| KPNB1 | 1704 | 9.57E-05 | 0.00166211 | -0.586127389 |
| MPZL1 | 1006 | 6.06E-07 | 1.78E-05 | -0.585895875 |
| LCN8 | 1246 | 7.18E-06 | 0.00017041 | -0.58552408 |
| ZBTB21 | 670 | 6.26E-09 | 2.77E-07 | -0.585269337 |
| PAPSS1 | 779 | 4.04E-08 | 1.53E-06 | -0.584799057 |
| PTPRD | 624 | 2.31E-09 | 1.09E-07 | -0.584614681 |
| IRAK1 | 1337 | 1.46E-05 | 0.0003229 | -0.584354774 |
| TMEM104 | 528 | 2.08E-10 | 1.17E-08 | -0.58320189 |
| CDKN1B | 925 | 2.47E-07 | 7.90E-06 | -0.58246425 |
| AP2B1 | 1758 | 0.00012175 | 0.00204925 | -0.58158094 |
| FTSJ1 | 1184 | 4.08E-06 | 0.0001018 | -0.581537653 |
| EIF3A | 2184 | 0.00066206 | 0.00897031 | -0.581226955 |

TableS3

| Genes in poverty-exposed signature | Genes in unexposed signature |
| --- | --- |
| ACTB | SMAD2 |
| PIK3C3 | BNIP2 |
| NCF4 | PSMB1 |
| PSME2 | ADA |
| GSDME | SLBP |
| PLCG1 | ABCE1 |
| VAV3 | RRP1B |
| ATP6V1G1 | UVRAG |
| VAV1 | MS4A1 |
| PSMA5 | EIF3B |
| MYO10 | TCP1 |
| DOK3 | PSMC6 |
| ITPR1 | TMEM123 |
| WASF2 | BLK |
| SYK | BTAF1 |
| HGSNAT | B3GNTL1 |
| B2M | POLR2B |
| LAIR1 | LDHA |
| AGPAT2 | ADAM10 |
| CD19 | PPP2R5C |
| JUN | CAST |
| UBA7 | EEF1B2 |
| MME | KPNB1 |
| SPTAN1 | MTHFD2 |
| LAPTM5 | GDI2 |
| PIK3IP1 | CORO1A |
| ARHGDIB | ETF1 |
| CPXM1 | RHOH |
| SDC2 | MACROH2A1 |
| BIRC7 | SSR1 |
| KLHL6 | IGBP1 |
| FADS3 | DAXX |
| CALM3 | TMX1 |
| CD74 | HDAC2 |
| SLAMF6 | NSMAF |
| PCNA | PFKFB3 |
| MSR1 | MYC |
| HLA-DMB | PUDP |
| RBMS3 | IRF8 |

DRAM1
TMSB4X
HBS1L
FKBP1A
FOXP1
CHCHD7
RUBCNL
MYADM
TRAF5
PHYH
LITAF
NDFIP1
CD52
TSPO
ZNF608
OAS3
TRIM38
SOCS2
HNRNPA2B1
SMARCA4
CD27
PTPN7
FOXO1
GBP4
RAG1
HLA-DPA1
ISG20
RAG2
HLA-DQB1
CIITA
HLA-DQB2
H3C4
HLA-DRA
LTB
HLA-DPB1

PAICS
EIF4A3
WAPL
CNIH1
LSM2
ASF1A
UBE2J1
SYNCRIP
ABCB7
STK17B
GRB2
ITGA4
NFATC3
SRP72
BLNK
GLIPR1
FTSJ1
MSL3
EIF4A1
DDX21
IL2RG
RBM3
C1QBP
BCAT1
PLEK
NUP210
IRAK1
CCNC
SLC7A5
TP53
ELAC2
RSL1D1
LPXN
MTDH
ADK
CCT8
TCOF1
RABGAP1L
PMAIP1
LCP1
SH3TC1

CSGALNACT1
LILRB2
GPM6B
ALOX5
PNISR
P2RX5
CD9
CD69
EZR
SPPL2B
CDKN1B
SOD2
PGD
FRMD4B
MTHFD1
FHL1
PDLIM1
ACAT1
FOS
SH3BGRL
HSP90AB1
TALDO1
CANX
LAT2
RSU1
CPNE3
CD34
PPA1

TableS4_Flow Percentage

| Sample Number | %CD45low | # of sequenced cells | % CD45high | # of sequenced cells | % CD45highCD3+ | % CD45highCD19+ | % CD45highCD14+ |
|---|---|---|---|---|---|---|---|
| Unexposed 1 | 88.10% | 185 | 11.80% | 176 | 37.30% | 27.70% | 32.70% |
| Unexposed 2 | 30.80% | 188 | 56.40% | 88 | 10.30% | 76.60% | 3.20% |
| Unexposed 3 | 98.80% | 185 | 0.98% | 0 | 64.60% | 8.04% | 27.30% |
| Unexposed 4 | 95.80% | 282 | 4.03% | 88 | 32.20% | 30.60% | 33.70% |
| Unexposed 5 | 93.30% | 187 | 6.08% | 88 | 23.20% | 56.80% | 19.50% |
| Unexposed 6 | 87.50% | 122 | 11.60% | 87 | 39.30% | 38.50% | 10.70% |
| Unexposed 7 | 79.80% | 137 | 18.80% | 88 | 16.00% | 70.90% | 6.19% |
| Poverty-exposed 1 | 65.50% | 188 | 31.90% | NA | 49.80% | 41.10% | 5.63% |
| Poverty-exposed 2 | 89.90% | 188 | 8.51% | 88 | 31.20% | 18.80% | 43.80% |
| Poverty-exposed 3 | 3.21% | 94 | 95.50% | 88 | 16.85% | 75.70% | 1.01% |
| Poverty-exposed 4 | 85.50% | 188 | 14.10% | 87 | 54.40% | 28.40% | 7.55% |
| Poverty-exposed 5 | 86.90% | 140 | 12.30% | 88 | 24.10% | 63.60% | 1.80% |
| Poverty-exposed 6 | 95.50% | 186 | 4.02% | 88 | 52.70% | 36.60% | 4.30% |
| Poverty-exposed 7 | 79.40% | 186 | 20.00% | 88 | 28.00% | 50.10% | 13.60% |

TableS5

| GSE181157_ETV6-RUNX1 | GSE181157_Hyperdiploid |
|---|---|
| Poverty-exposed_16-003 | Poverty-exposed_16-049 |
| Poverty-exposed_16-106 | Poverty-exposed_16-257 |
| Poverty-exposed_16-115 | Poverty-exposed_16-291 |
| Poverty-exposed_16-304 | Poverty-exposed_16-335 |
| Poverty-exposed_16-312 | Poverty-exposed_16-342 |
| Unexposed_16-001 | Unexposed_16-048 |
| Unexposed_16-002 | Unexposed_16-090 |
| Unexposed_16-006 | Unexposed_16-131 |
| Unexposed_16-012 | Unexposed_16-141 |
| Unexposed_16-050 | Unexposed_16-158 |
| Unexposed_16-065 | Unexposed_16-161 |
| Unexposed_16-070 | Unexposed_16-176 |
| Unexposed_16-074 | Unexposed_16-182 |
| Unexposed_16-076 | Unexposed_16-186 |
| Unexposed_16-094 | Unexposed_16-194 |
| Unexposed_16-100 | Unexposed_16-199 |
| Unexposed_16-108 | Unexposed_16-215 |
| Unexposed_16-140 | Unexposed_16-239 |
| Unexposed_16-152 | Unexposed_16-249 |
| Unexposed_16-221 | Unexposed_16-272 |
| Unexposed_16-238 | Unexposed_16-273 |
| Unexposed_16-267 | Unexposed_16-279 |
| Unexposed_16-293 | Unexposed_16-287 |
| Unexposed_16-322 | Unexposed_16-302 |
| Unexposed_16-350 | Unexposed_16-306 |
| Unexposed_16-352 | Unexposed_16-315 |
| Unexposed_16-354 | Unexposed_16-338 |
| Unexposed_16-363 | Unexposed_16-339 |
| | Unexposed_16-374 |
